# Deciphering the Timing and Impact of Life-extending Interventions: Temporal Efficacy Profiler Distinguishes Early, Midlife, and Senescence Phase Efficacies

**DOI:** 10.1101/2024.03.27.585737

**Authors:** Nisi Jiang, Catherine J. Cheng, Qianqian Liu, Randy Strong, Jonathan Gelfond, James F. Nelson

## Abstract

Evidence that life-extending interventions are not uniformly effective across the lifespan calls for an analytic tool that can estimate age-specific treatment effects on mortality hazards. Here we report such a tool, applying it to mouse data from 42 compounds tested in the NIA Interventions Testing Program. This tool identified agents that either reduced (22) or increased (15) mortality hazards or did both (2) in at least one sex, most with marked variation in the duration of efficacy and magnitude of effect size. Only 8 reduced mortality hazards after 90% mortality, when the burden of senescence is the greatest. Sex differences were common. This new analytic tool complements the commonly used log-rank test. It detects more potential life-extending candidates (22 versus 10) and indicates when during the life course they are effective. It also uncovers adverse effects.

## Introduction

The search for pharmacological interventions that extend the healthy lifespan has increased markedly in recent years, spurred by the discovery of a wide range of compounds, such as rapamycin and acarbose, that lengthen life of model organisms^1-3^. Whether these life-extending agents act broadly by reducing mortality hazard throughout the lifespan or only affect mortality during part of the life course remains unclear, in part due to the limitations of statistical tests usually used in aging research. The log-rank test^4^ is the most commonly used statistical tool to determine whether an intervention, be it pharmacologic, genetic, or nutritional, is life-extending. However, its use as the primary and often only tool for this purpose is questionable for several reasons. First is its requirement for proportional hazards (PH) between compared groups, implying that treatment effects on mortality remain constant over time^5,6^. This assumption does not align with the evidence that many interventions exert varying impacts at different life stages^7^. For example, in an earlier analysis of data from the Interventions Testing Program (ITP), we found that many interventions do not adhere to the PH assumption, thus challenging the applicability of the log-rank test in these contexts^7^. When the PH assumption is not met, there are many analytic tools that can be used^6-9^. Our previous approach for these interventions used the Gehan test, which is more robust to the PH consistency requirement and more sensitive to effects during early adulthood^6^. Despite its strengths, the Gehan test has its own drawback: a diminished sensitivity to effects manifesting at later life stages, when mortality and morbidity rates are highest^10^.

To assess the effects of interventions on the final phase of the aging process, methods like the Wang-Allison and the Gao-Allison test have been developed to determine if treatments extend the “maximum” lifespan^11,12^. However, these tests do not evaluate whether an intervention specifically reduces age-specific mortality in the last phase of life when frailty, cognitive impairment, chronic disease, and other burdens of senescence peak. Although the Gompertz model has been used for evaluating age-specific or time-varying effects, it is limited by its strict parametric assumptions about the shape of the hazard function^13^. The limitations of these approaches underscore the need for a more flexible tool for evaluating longevity interventions, especially one that accommodates potentially variable impacts of treatments across an organism’s lifespan. Such methods should pinpoint when, for how long, and to what extent an intervention significantly alters the mortality risk. This capability is particularly crucial for identifying interventions that mitigate mortality toward the end of life when the exponential increase in the burden of senescence is greatest.

Numerous statistical methods for assessing the time-varying efficacy of drugs, including chemotherapeutics, have been developed and published^6,14-21^. However, these approaches have not seen widespread adoption in clinical trials, nor have they been applied in longevity studies^22^. A key barrier to their use is the lack of accessible implementations, coupled with the need for substantial user expertise to effectively tune these models. In response to these challenges, we introduce a nonparametric method termed the Temporal Efficacy Profiler (TEP), which estimates the time-varying hazard ratio and visualizes age-specific effects on mortality risk. TEP can identify when, for how long, and to what extent an intervention significantly influences mortality risk, thereby overcoming the major limitations of traditional methods like the log-rank test.

In our approach, we employ the Rebora method, implemented in the *‘bshazard’* R package, to calculate the age-specific mortality risk for the treatment (λ_*Treatment*_(*t*)) and control (λ_*Control*_(*t*)) arms separately^23^. Rebora et al. utilized B-splines within generalized linear Poisson models,incorporating a robust model selection process that automates tuning and eliminates the need for users to manually select smoothing parameters. TEP is defined as:

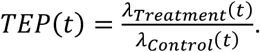

We considered two approaches to estimate confidence intervals (CIs) of the hazard ratios: the asymptotic method and the bootstrap method. Although bootstrap methods are broadly applicable, they are computationally intensive. On the other hand, simulation studies demonstrated that the asymptotic method is generally more conservative in terms of the coverage probability, especially at age extremes (Online methods and Figure S1). For the asymptotic method, we derived pointwise analytical CIs for the hazard ratio as the sum of two asymptotically normal estimates based on the variance of the difference the log hazards for each group as shown below

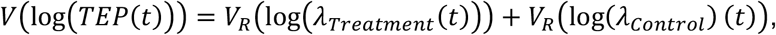

where *V*_*R*_ represents the Rebora estimator variance^23,24^. In addition, we used pointwise bootstrap CIs to describe the time-varying hazard profile^25,26^. The bootstrap method estimates CIs that are more sensitive to differences at age extremes, but it also has slightly lower coverage probability under the null hypothesis (Online methods and Figure S1). For conciseness, we only present the results from the asymptotic method, and only when corroborated by the bootstrap method. All significant findings, along with their corresponding mortality hazard ratios, are provided in the supplementary materials (Table S1, Figures S3 and S4).

In addition, we developed a color-coded visualization system to better communicate the statistical results. The code is publicly available, and we provide a user-friendly R script with instructions to facilitate its use by any investigator. (Github link: https://github.com/liu-dada/Temporal-Efficacy-Profiler).

To assess the utility of this approach, we utilized publicly available data from the ITP up to 2022, comprising 42 compounds evaluated in over 27,000 genetically heterogeneous mice at 3 geographically distinct sites^27^. These agents were tested alone or in combination in 132 trials, examining the effects of sex, dosage, and age of treatment initiation. Ten of these agents have been identified by log-rank testing to significantly extend lifespan in at least one sex^28^. This is the largest publicly available compendium of mouse survival data from tests of compounds with lifespan-extension potential, an exemplary resource for testing the efficacy of our analytic tool.

## Results

### Development and validation of the TEP to determine the timing and impact of life-extending candidates

Figure 1 illustrates how the TEP identifies age-specific effects of an intervention on the mortality hazard, using the ITP test of green tea extract (GTE) in females as an example^29^. Details of the analysis are described in “Online Methods”. It should be noted that GTE had no significant effect on survival by log-rank testing^29^. Figure 1A shows the Kaplan-Meier survival plots for treatment and control groups. These plots indicate that the proportional hazard assumption is likely violated due to the crossing survival curves, which was confirmed by the z-test^7^.

**Figure 1.**
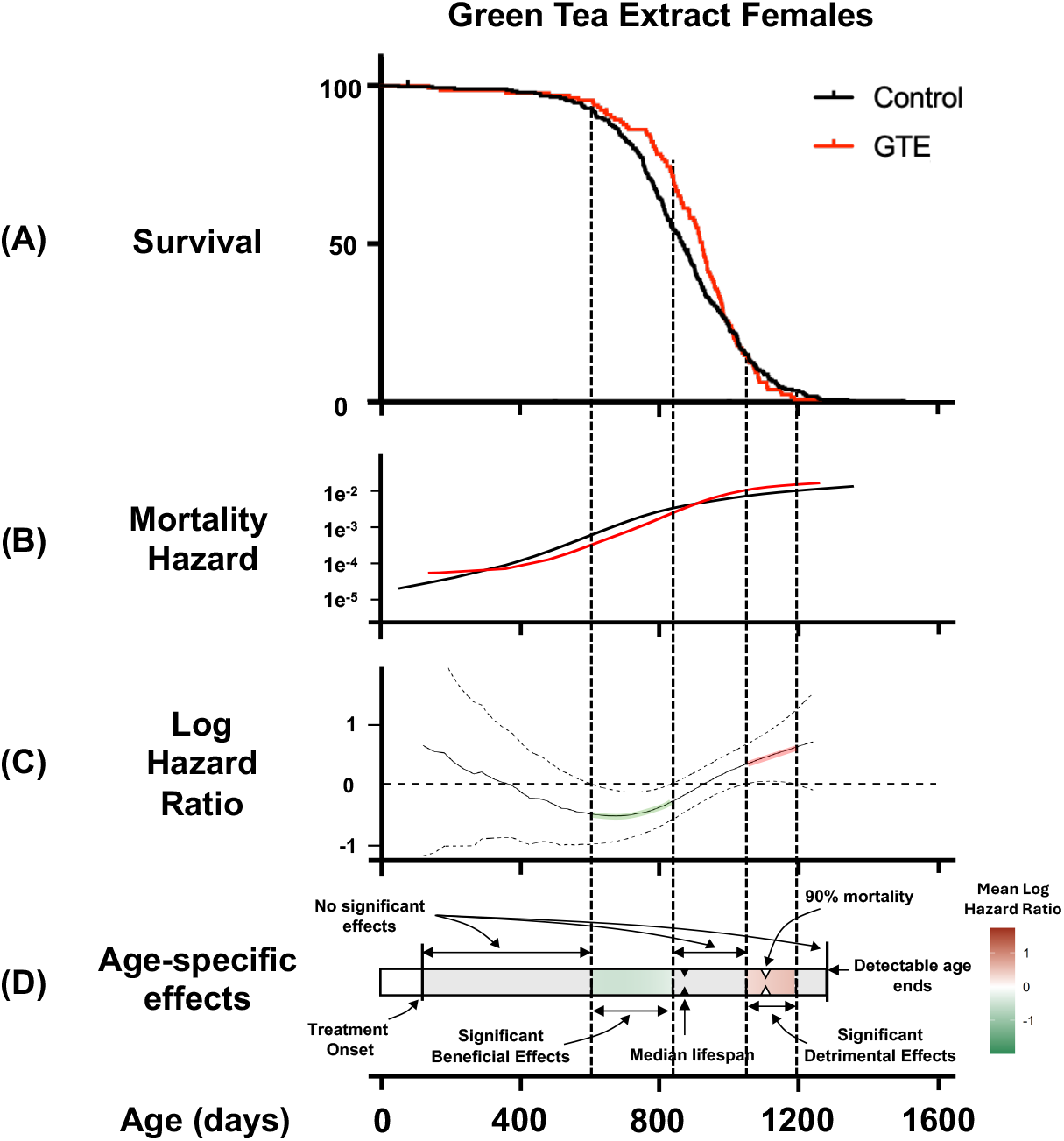
Graphical representation of the TEP. Survival data are from the test of Green Tea Extract in females^29^. **A)** Kaplan-Meier survival curves of the GTE-treated female mice (Red) and control female mice (Black); **B)** Age-specific mortality hazards of GTE-treated and control mice groups; **C)** Mortality hazard ratio between GTE-treated and control mice groups and 95% confidence intervals shown as dashed lines; **D)** Life course heat map visualization of the age-specific effects of GTE on the mortality hazard ratio. Vertical dashed lines mark the boundaries of significant effects on the mortality hazard ratio based on the ages when the 95% confidence intervals in Figure C cross 0.

Figure 1B is a graphical representation of the mortality hazards of the control and GTE-treated groups throughout the period of testing, using the Rebora method^30,31^. The mortality hazard of the GTE-treated group is reduced relative to that of the control group before the median lifespan, but shortly thereafter crosses over, exceeding that of the control group.

Figure 1C shows the application of the TEP to the GTE data. The log ratios of the mortality hazards of GTE-treated and control groups shown in Figure 1B are calculated based on the mortality hazard estimated by Rebora method^23^. Negative log hazard ratios indicate beneficial effects of GTE (lower mortality hazard), while positive values suggest detrimental effects. The 95% CIs for the mortality hazard ratio were estimated by asymptotic and bootstrap methods, with the asymptotic CI shown as dashed lines. Significant beneficial effects are marked by upper 95% CIs remaining below zero (marked in green), whereas significant adverse effects are indicated when lower 95% CIs exceed zero (marked in red). The duration (age range) of significance is bounded by the ages when the 95% confidence limit crosses 0, as illustrated.

This analysis reveals that GTE reduced mortality hazards during midlife but increased mortality hazards toward the end of life.

Figure 1D integrates the features of Figure 1C into an annotated horizontal heatmap to assist in cross-compound comparisons. The heatmap ranges from birth to the death of the last subject in either control or treated group, starting blank and transitioning to color with the onset of treatment. Gray indicates no significant effect, green marks periods of significant mortality reduction, and red denotes significant increases. The color intensity correlates with the effect size (log HR), allowing for a direct comparison of intervention impacts across different timelines as illustrated in Figure 2. In this example, TEP complements and adds value to the log-rank test, pinpointing the specific ages and durations over which GTE significantly alters age-specific mortality hazards.

**Figure 2.**
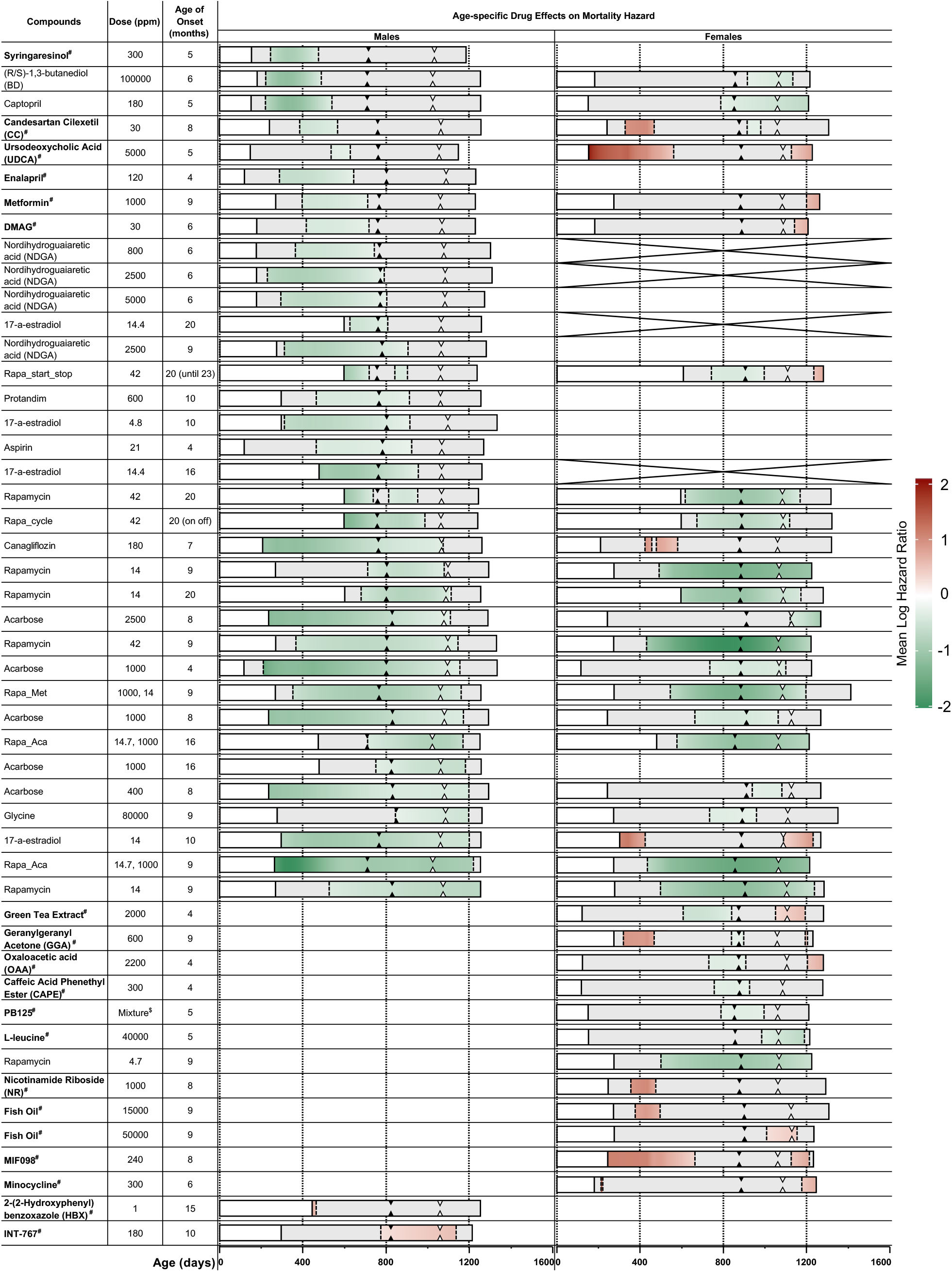
Life course heat maps of interventions that significantly modified mortality hazard. Only interventions confirmed by both bootstrap and asymptotic methods are displayed, with the asymptotic method results used as the representative data. Interventions are ranked by the age when beneficial effects ceased in males (from earliest to latest). The remaining interventions are ranked by cessation age of beneficial effects in females, followed by detrimental effects in females (ranked by cessation age of effect), and the remaining ranked from longest to shortest duration of detrimental effects in males. Each row represents an individual trial of one intervention in a single cohort. Each intervention involved one compound or a combination of two, with dosage and starting age of treatment listed. The color-coded bands denote the temporal significance of drug effects: white indicates the period before treatment onset, gray marks periods with no significant effects, green indicates periods of significant beneficial effects, and red denotes intervals of significant detrimental effects. The solid black triangle indicates the median lifespan of the control group for each trial, and the open triangle marks the age of 90% mortality of the control group. Empty cells indicate no significant effects detected by both methods, while cells crossed indicate there is no trial tested. Note that the reader can rearrange trials in any order using an Excel spreadsheet (Table S1). Footnotes:# New Compounds that significantly affect mortality (i.e., not identified by the log-rank test) are also noted in **bold** font. $ PB125 is a mixture of luteolin, withaferin A, and carnosol, dosages refer to publication^40^.

Since we added features into the established *‘bshazard’* function, we conducted two simulation scenarios assessing the TEP performance. Under null and alternative hypotheses, simulation results show the method’s robustness in estimating accurately coverage probabilities for confidence intervals across life spans (Online Methods and Figure S1). The first scenario confirmed the accuracy of the asymptotic confidence intervals and the conservative nature over an entire lifespan, while the second scenario demonstrated TEP’s ability to correctly indicate treatment effects, outperforming the log-rank test in detecting non-proportional hazards (Online Methods and Figure S1).

### Greater sensitivity and precision in identifying mortality-modifying interventions

Figure 2 presents heatmaps of interventions identified by the TEP that significantly reduced or increased the age-specific mortality hazard during treatment. Comprehensive heatmaps generated by both the asymptotic and bootstrap methods are provided in Table S1. The hazard ratio plots, which underlie these heatmaps and were calculated via time-varying hazard ratio analysis, are displayed in Figures S3 and S4 for males and females, respectively.

Interventions in Figure 2 are ranked based on the age at which their beneficial effects ceased in males, from earliest to latest. For interventions that were identified differently by the two methods, further investigation may be required. Readers can reorder these data as they see fit using the spreadsheet in Table S1. In this discussion, we focus on the interventions identified to be significant by both methods.

Twenty-eight compounds, consisting either of a single agent or a combination of two agents, at one or more doses, initiated at varying ages, significantly modified the mortality hazard in one or both sexes at one or more periods during the treatment period. This analysis identified 11 new compounds that significantly reduced mortality in at least one sex during treatment but were overlooked by the log-rank test: namely, candesartan cilexetil (CC), caffeic acid phenethyl ester (CAPE), 17-dimethylaminoethylamino-17-demethoxygeldanamycin hydrochloride (DMAG), enalapril, GTE, L-leucine, metformin, oxaloacetic acid (OAA), PB125, syringaresinol (Syr), and ursodeoxycholic acid (UDCA). The new analysis also identified 14 compounds that were detrimental (i.e., increased mortality) in one or both sexes at one or more periods of treatment. The duration of significant benefit or detriment varied markedly from weeks (e.g., H2-(2-Hydroxyphenyl) benzoxazole (HBX)) to almost the entire treatment period (e.g., rapamycin + acarbose). Most compounds only reduced mortality or only increased mortality. Two exceptions were CC and GTE in females. Effect sizes, indicated by the color intensity, varied markedly during the periods of benefit and detriment. Acarbose had its greatest benefit at the initiation of treatment, waning progressively thereafter. Effect sizes of other compounds, such as butanediol and captopril in males and many of the different rapamycin trials in females peaked during the middle of treatment. A few interventions showed a steady increase in effect with continued treatment (e.g., glycine in males and leucine in females).

### Only a fraction of interventions reduced mortality at later ages

A strength of the TEP is its ability to estimate when during the life course and for how long an agent exerts its effect on survival. In males, 16 interventions reduced mortality hazards at some period during the life course (Figure 2). Of these, 9 compounds only reduced mortality risk in early and mid-adulthood (i.e., before the median lifespan): Syr, (R/S)-1,3-butanediol (BD), CC, captopril, enalapril, UDCA, metformin, DMAG, and nordihydroguaiaretic acid (NDGA) at 800 ppm. The two higher doses of NDGA had a slightly longer period of benefit, but only for a short period beyond the median lifespan. By contrast, in females, of the 11 agents that reduced mortality risk at some stage of life, only GTE reduced mortality during early-to mid-adulthood.

In males, five compounds tested in 11 trials demonstrated reduced mortality after attainment of median lifespan, although these effects vanished before mice attained the 90% mortality benchmark: 17α-estradiol, aspirin at 21 ppm, Protandim, high doses of NDGA, and 3 of 4 late-onset (20 mo) rapamycin treatments. Notably, only 6 of the 17 compounds that reduced mortality in males did so at ages beyond the 90% mortality threshold: canagliflozin, acarbose, 17α-estradiol, glycine, simvastatin, rapamycin, and cocktails of either acarbose or metformin with rapamycin. In females, in contrast to males, 10 of 11 beneficial interventions reduced mortality mainly at ages after attainment of median lifespan. Five compounds reduced mortality after 90% mortality, including most trials involving rapamycin, acarbose, BD, L-leucine, and captopril.

### Some compounds have adverse effects on mortality hazards

One goal of the ITP has been to ensure against possible deleterious side effects of potential life-extending interventions, especially those already being marketed. Until now, the ITP has only identified two life shortening interventions using the log-rank test^32^. Here the TEP revealed 15 trials with 14 compounds that increased the mortality hazard at one or more periods of treatment: 2 in males (HBX and INT-767) and 12 in females (CC, metformin, DMAG, canagliflozin, 17α-estradiol, GTE, minocycline, geranylgeranyl acetone (GGA), fish oil, nicotinamide riboside (NR), UDCA, and MIF098) (Figure S2).

### Sex differences in the effect of pharmacological interventions

Marked sex differences in the responses to life-extending compounds are one of the key outcomes of the ITP^28^. The TEP unveiled even more sex differences. It identified 5 additional compounds that only benefited males: Syr, enalapril, metformin, DMAG, and UDCA, and 7 compounds that only reduced mortality in females: OAA, CAPE, PB125, Leu, and GTE. Notably, 6 interventions, UDCA, CC, metformin, DMAG, canagliflozin, and 17α-estradiol, exhibited beneficial effects in males but detrimental effects in females (Figure S2). More compounds adversely affected survival in females (12) than in males (2). Moreover, most compounds with negative effects exerted their effect on females almost from the beginning of treatment. The detrimental effects waned during the 2^nd^ year of life but sometimes reappeared in the final stage of life.

## Discussion

The TEP presented here promises to be broadly useful and impactful for aging research. Rather than repeatedly testing different quantiles (i.e., median, 90^th^ percentile, etc.), we have introduced a descriptive approach that reveals the age-specific effects of interventions on the mortality hazard. This opens the door to more granular insights about the actions of an intervention. Such insights can lead to more targeted application of interventions and a better understanding of the underlying mechanisms. The analysis does this by providing estimates of when and for how long during the life course an intervention reduces (or in the case of detrimental effects, increases) age-specific mortality. It also provides an estimate of the effect size of an intervention and how the strength of its effect changes over the course of treatment. None of this information is attainable by the log-rank test, the current standard for evaluating longevity interventions.

The TEP can distinguish interventions that specifically reduce mortality during senescence from those that only affect survival during midlife or earlier. This is important in the search for therapeutic interventions that benefit individuals of advanced age when the burdens of senescence are greatest. The TEP is also sensitive to adverse effects, which is critical for pre-clinical models that aim to be translatable. Furthermore, the method is sensitive to sex differences in timing, duration, and efficacy of interventions, providing further impetus to probe the mechanisms underlying the growing number of sexually dimorphic traits in aging. Here we discuss some of the ways the new information provided by this analytic tool that can assist drug discovery and the search for the underlying mechanisms that drive aging. Additional applications will likely emerge as its adoption spreads within the geroscience community.

A major discovery using this tool is that most interventions exhibited age-related changes in drug efficacy across the detectable treatment period (Figure 2). This observation is not readily apparent by visual inspection of most Kaplan-Meier plots and is not obtainable from the log-rank tests. Very few interventions significantly reduced (or increased) mortality through the entire course of treatment. Most were only effective for less than half of the treatment duration. This calls for explanation, and the answers are likely to lead to better interventions and greater insight into the mechanisms of aging. The age-specific decrease, increase, or loss of efficacy of an intervention may reflect age-related changes in pharmacokinetics or pharmacodynamics, leading to suboptimal dosage. This finding opens the door to developing age-specific doses to sustain efficacy for longer periods and raises awareness of the importance of understanding the role of aging in pharmacokinetics. It is plausible that the aging processes or causes of mortality change with age and the intervention loses efficacy because it no longer targets the underlying pathways. Whatever the reason, this tool has uncovered a critical variable that needs to be considered in interventional geroscience.

Another important outcome of the application of the TEP to longevity data is the finding that only a subset of the interventions in the ITP database affected age-specific mortality rates in the last half of the lifespan, and even fewer affected mortality rates after the age when 90% of the control cohort has died^11^. Diet restriction has long been considered an example of an intervention that retards aging processes broadly, because it extends the age of 90% mortality, distinguishing it from many interventions that only extend the median lifespan^33,34^. Many studies, including the ITP, use the Wang-Allison test as a discriminator for interventions that do or do not extend the maximum lifespan based on the 90% mortality measure. However, this test does not distinguish whether an increase in the age of 90% mortality reflects the effects of reduced mortality accumulated during earlier ages from the effects of the age-specific mortality reduction at or near the age of 90% mortality. This distinction is of particular importance to a major goal of geroscience: namely, to identify compounds and discover the underlying mechanisms that extend the maximum lifespan by reducing age-specific mortality during the later stages of life when the burden of senescence is greatest. The TEP provides such a measure by indicating whether the intervention specifically reduces mortality rates in the final stage of life. Only a subset of the interventions reported by the ITP as “lifespan extending” using log-rank analysis reduced mortality hazard after the median lifespan, and even fewer did so at later ages.

Nevertheless, compounds that only reduce mortality during the first half of adult life should not be discounted. Reducing mortality at any stage of life can be impactful, especially when considering translatability to humans. For example, the male mortality disadvantage, compared to females, is greatest in the first half of adult life in both humans and UM-HET3 mice^31^. It is noteworthy that most of the compounds that are only effective in males are also only effective during the first half of the lifespan. Castration of UM-HET3 males before puberty eliminates this mortality disadvantage^30^. If any of the compounds that only eliminate the male mortality disadvantage during this period without interfering with male reproductive function, the societal impact if clinically translatable would be great^28,35^.

Not only is this method more sensitive to agents that reduce age-specific mortality, it is also more sensitive to those that increase mortality. The ITP has never identified adverse effects using the log-rank test until recently^32,36^. This new tool revealed 15 trials involving 14 compounds that increased mortality hazards in at least one gender. There was a marked sex difference. Only 2 trials showed detrimental effects in males compared to 13 trials in females. Some compounds, including canagliflozin and high doses of 17α-estradiol markedly reduced mortality in males but were harmful in females. This finding has been confirmed in a recent ITP trial, where canagliflozin significantly prolonged lifespan in males, but shortened lifespan in females^32,36^. These findings underscore the need for sex-specific testing of life-extending candidates.

The TEP can detect reversals of the benefit of compounds across the life course. GTE in females reduced mortality before the median lifespan but increased mortality at later ages— another discriminator not possible using the log-rank test. There is precedence for this reversal. In humans, individuals reporting the lowest intake of dietary protein had reduced mortality from cardiovascular disease and cancer before 65 years of age, but this relationship reversed after 65^37^. Mice with reduced branch chain amino acid intake had extended life when the diet began in early adulthood, but their lifespan was unaffected when the diet was initiated at a later age^38^.

Age-related changes in pharmacokinetics and pharmacodynamics may play a role here. For example, blood levels of canagliflozin, whose beneficial effects in males diminish with age, are 2-3-fold higher in older males^39^.

Another strength of the TEP is its heightened sensitivity to potential life-extending candidates. It identified over twice as many as the log-rank test. This is due in part to its ability to identify age-specific effects on the mortality hazard unimpeded by the requirement of the log-rank test for consistent proportional hazard across the duration of treatment. The newly identified compounds generally have smaller effect sizes and shorter durations of positive effect compared to those identified by the log-rank test. However, given their geroprotective potential and the fact that most trials have only used one dose, they deserve further study. It is important to emphasize that this statistical tool should not be used as a final arbiter of any candidate for mortality reduction and lifespan extension (or adverse effect), but rather should be considered a screening tool for identifying potential candidates that deserve follow-up—for example with different doses. Type 1 errors (i.e., false positives) during initial screens are more acceptable and preferable to false negatives.

It is important to acknowledge the limitations of this method. Although we employed both asymptotic and bootstrap methods to identify significant effects and we only present findings that were consistently identified by both methods, effects that emerge at age extremes (after 90% mortality) may require further validation due to the relatively small sample size during that period. For example, the detrimental effects observed in DMAG and metformin treatments in females were only evident during a brief window after 90% mortality. On the other hand, we emphasize that these detrimental effects warrant close attention, especially since many of the compounds studied are readily available over the counter, raising potential safety concerns.

Compounds showing detrimental effects, even if detected by only one method, deserve further investigation. Another limitation is that the TEP may require larger sample size than traditional log-rank test. While there is no specific sample size requirement for TEP analysis, we recommend using a sample size that meets the requirements of the log-rank test to ensure more reliable interpretation of the results.

The method currently does not explicitly consider uncertainty in the Time axis so the ages at which the treatment effect becomes nonzero are presented as point-estimates without confidence intervals. However, this limitation did not preclude consistent findings between similar treatments across several cohorts, such as the early effects of ACE inhibitors (Enalapril and Captopril) or early effects of different doses of NDGA. Statistically testing whether two different treatments have the same effect relative to control is more complex (testing whether the ratio of two hazard ratios is 1) and may require comparisons across cohorts.

While this methodology facilitates the estimation of time-varying treatment effects in comparison to a control, future enhancements could include explicit testing and quantification of differences between active treatments in terms of both timing and the extent of changes in mortality hazard ratios. It would be particularly insightful to assess different dosages of a single compound to pinpoint the optimal dosage for specific age intervals.

In conclusion, this new analytic tool will lead to a better understanding of the impact of interventions on survival, especially in the field of aging research. Testing interventions on survival across the life course is not only time-consuming but also expensive. From such studies, we should derive not just a p-value but also gain insight into the ages when interventions are effective or deleterious. This method can provide a more comprehensive evaluation of lifespan interventions, thereby enhancing our understanding of the mechanisms of aging and age-related risk factors.

## Author contributions

Conceptualization: CJC, NJ, JG, QL, JN, RS

Methodology: CJC, NJ, JG, QL, JN

Funding acquisition: RS, JN

Data analysis: QL, NJ, CJC, JG, JN

Writing: NJ, JN, JG

## Acknowledgments

Dr. Strong has been honored with the Senior Research Career Scientist award (# IK6 BX006289) from the Department of Veterans Affairs. The funding for this research was provided by the Center for Testing Potential Anti-Aging Interventions (5U01AG022307), the Nathan Shock Center of Excellence in Basic Biology of Aging (5P30AG013319), as well as NIH grant 5T32AG021890-15, alongside a fellowship from the Glenn Foundation awarded to CJC.

## Data availability

All raw data can be found in Mouse Phenome Database (MPD; https://phenome.jax.org/projects/ITP1) and also will be available upon request.

## Code availability

Code is uploaded to Github link: https://github.com/liu-dada/Temporal-Efficacy-Profiler

## Online Methods

### Data availability, mouse model, and husbandry

The datasets employed in this study are sourced from the Mouse Phenome Database (MPD; phenome.jax.org), encompassing all data from the Interventions Testing Program (ITP) spanning from 2004 to 2022. This dataset incorporates 13 distinct cohorts, integrating data across three research facilities to ensure the robustness and reproducibility of the findings. The ITP employed the UM-HET3 mouse line, a genetically heterogeneous model, chosen for its relevance to the genetically diverse human population. UM-HET3 mice are bred according to a specific crossbreeding protocol: BALB/cByJ females are mated with C57BL/6J males to produce F1 hybrid females, which are then bred with F1 hybrid males derived from mating C3H/HeJ females with DBA/2J males. This breeding strategy is designed to maximize genetic diversity within the model, thereby approximating the genetic variability inherent in human populations and increasing the translational value of the research findings. The mice designated for longevity assays were maintained under controlled environmental conditions, with a constant ambient temperature of 25°C and a regulated photoperiod of 12 hours light/12 hours darkness. Nutritional needs were met with ad libitum access to the Purina 5LG6 diet, alongside specific drugged food formulations as per experimental requirements. Housing protocols were optimized for social enrichment and welfare, accommodating up to three males or four females per standard laboratory enclosure, in accordance with established ethical guidelines. Rigorous daily health assessments were conducted by trained staff to monitor the well-being of the subjects, promptly identify morbidity signs, and implement early intervention strategies as necessary. This proactive health management approach minimized unnecessary suffering and ensured the reliability of longevity data. The specifics of drug administration, including dosage, frequency, and duration, as well as the rationale behind the selection of intervention agents, are detailed in the original published reports, providing a comprehensive overview of the therapeutic strategies explored in this body of research.

### Description of the Temporal Efficacy Profiler

TEP adapted the Rebora method (implemented in *‘bshazard’* package in R^23^) to generate a nonparametric smoothed estimate of the baseline hazard rate for both treatment and control groups separately. We included site as an adjustment covariate within the models. We considered male and female as different groups, since most pro-longevity interventions exhibit significant sex differences, with more than half demonstrating efficacy exclusively in males^28^. In our analysis, mortality events occurring prior to the initiation of treatment were excluded to ensure that the hazard ratio estimates accurately reflect the treatment’s effect on survival. This exclusion criterion is crucial for eliminating bias arising from pre-treatment mortality, thus enhancing the validity of our findings.

The confidence intervals for the treatment hazard ratio were estimated using asymptotic and bootstrap methods. First, we used 1,000 bootstrapped replications to estimate the confidence intervals^25,26^, this method is similar to that reported previously in the analysis of the age-specific effects of sex^31^. Second, we employed the asymptotic method to derive pointwise analytical confidence intervals (CIs) for the hazard ratio. This was achieved by summing two asymptotically normal estimates based on the variance of the difference in log hazards between the groups, which was estimated by the Rebora method^23^.

### Data Visualization

The visualization method uses a color-coded band to depict treatment effects on hazard ratios, with the pre-treatment phase shown as a blank band. Upon treatment initiation, a gray color indicates no detectable effect, while significant effects are represented by changes in color intensity: beneficial effects cause the band to turn green, with the intensity reflecting the magnitude of negative log hazard ratios, and detrimental effects are shown in red, with intensity corresponding to positive log hazard ratios. The transition points where significant effects begin, or end are marked by dashed lines. Additionally, key lifespan metrics for the control group, such as median and maximum lifespan (when 90% have died), are highlighted to facilitate interpretation. All computational analyses were conducted in R (version 4.3, Vienna, Austria).

### Simulation

We conducted two simulation scenarios to validate the performance of the method under the null and alternative hypotheses. The simulations demonstrate the model’s accuracy under known conditions with the goal of demonstrating that the coverage of the confidence intervals was accurate and sensitive to variation. The first scenario used simulated datasets of similar sample size as the ITP case study with 300 controls and 150 treated under the null hypothesis, we used a same Gompertz distribution for both the treatment and control groups (Figure S1A). The specific Gompertz density was *f*(*t*∣*a*,*b*)=*be*^*at*^exp(−*b*/*a*(*e*^*at*^−1)) where a=log(300)/1200 and b=0.001/a. These values were chosen to reflect a similar hazard as female mice with a median survival of 741 days and a censoring rate of 10%. We conducted 500 simulations (Figure S1C) and estimated the *TEP*(t) hazard over the lifespan with the bootstrap (500 resamples each) and asymptotic 95% confidence intervals and computed the coverage probabilities of each. The results are shown in Figure S1E and S1G. The bootstrap confidence intervals exhibit close to nominal coverage until the later part of the lifespan where the coverage rate goes below 90% (Figure S1E). The asymptotic confidence intervals have >95% coverage and are conservative over the full lifespan (Figure S1G). This indicates good accuracy of the TEP method under the null hypothesis for the asymptotic confidence intervals, and the lower computational burden makes this an attractive option.

In the second scenario, we examined the alternative hypothesis where the control group (n=300) maintained the same Gompertz parameters as previously described, while the treatment group (n=150) was characterized by parameters a=log(50)/1200, b=0.0003/a. This resulted in a time-varying treatment effect, with an early hazard ratio greater than 1 (harm) and a later hazard ratio less than 1 (benefit), with the transition occurring around the median lifespan (Figure S1B and S1D). To evaluate the performance of the TEP and the log-rank test, we analyzed 100 simulated datasets by comparing the proportion of estimated hazard ratio 95% CIs that did not include 1 for TEP, against the proportion of log-rank tests rejecting the null hypothesis at alpha = .05. The CIs estimated using both bootstrap and asymptotic methods did not contain HR = 1 and correctly indicated the direction of the treatment effect in the early part of the curve in over 90% of simulations (Figure S1F and S1H). However, in the later part of the curve, bootstrap CIs did not contain HR = 1 and correctly indicated the treatment effect direction in approximately 60% to 90% of simulations (Figure S1F), whereas asymptotic CIs did so in about 60% of cases (Figure S1H). Around the median lifespan, where the hazards crossed, TEP appropriately covered HR = 1 at a near-nominal rate. As expected, the log-rank test exhibited reduced power (21%) to reject the null hypothesis in this scenario of nonproportional, crossing hazards.

---

**Figure S1.**
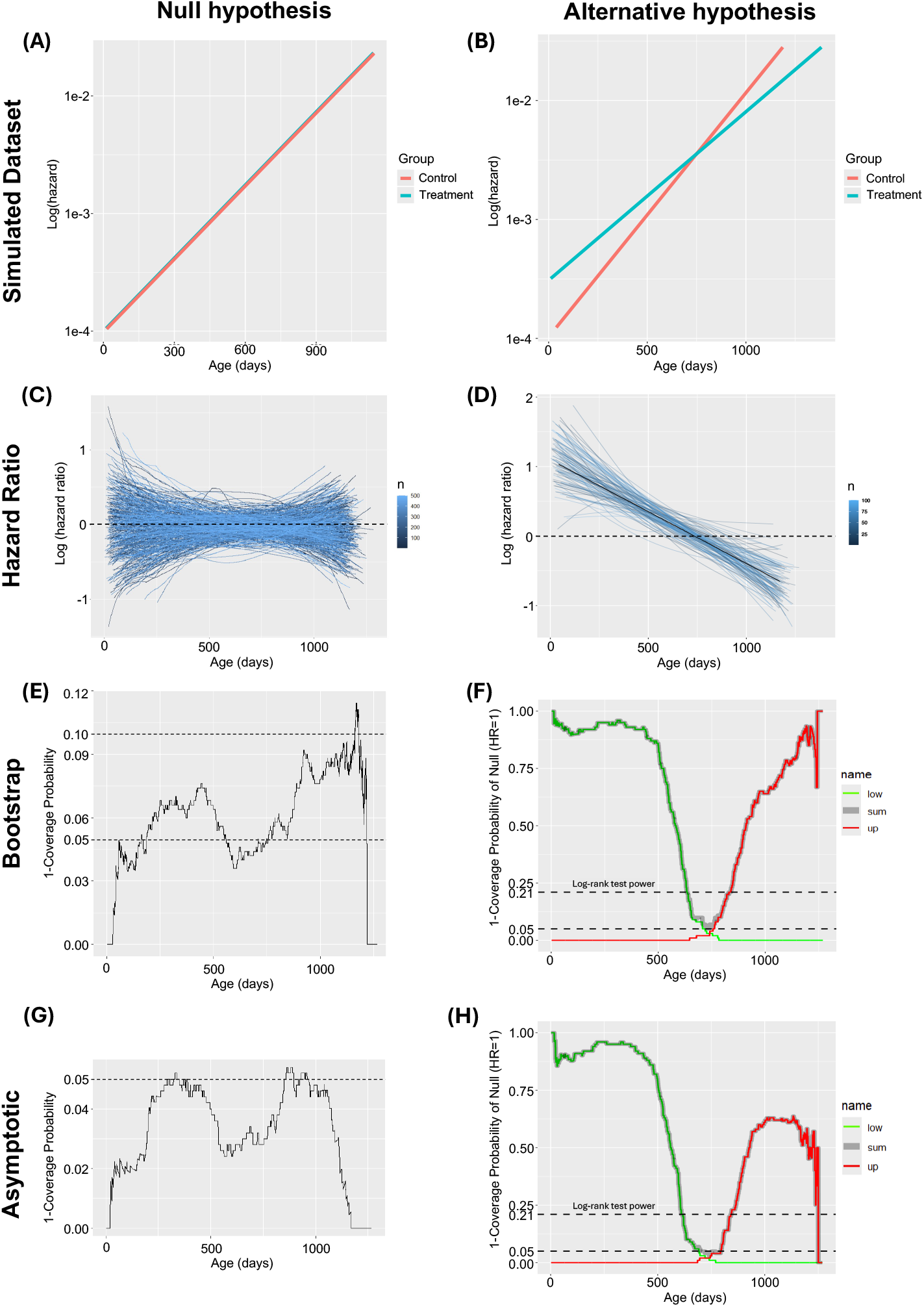
Evaluation of the performance of the asymptotic and bootstrap methods under null and alternative scenarios. (A) The Log mortality hazard for creating the 500 cases for null hypothesis scenario, the control and treatment group share the same age-specific mortality hazard; (B) The Log mortality hazard for creating the 100 cases for alternative hypothesis scenario, treatment group have high mortality hazard early in life, but lower mortality hazard later in life, which switch around the median lifespan; (C) Case specific Log hazard ratio of treatment to control under null-hypothesis. (D) Case specific Log hazard ratio of treatment to control under alternative hypothesis. (E) Summary of the performance of the bootstrap method under null hypothesis. (F) Summary of the performance of the bootstrap method under alternative hypothesis. (G) Summary of the performance of the asymptotic method under null hypothesis. (H) Summary of the performance of the asymptotic method under alternative hypothesis.

**Figure S2.**
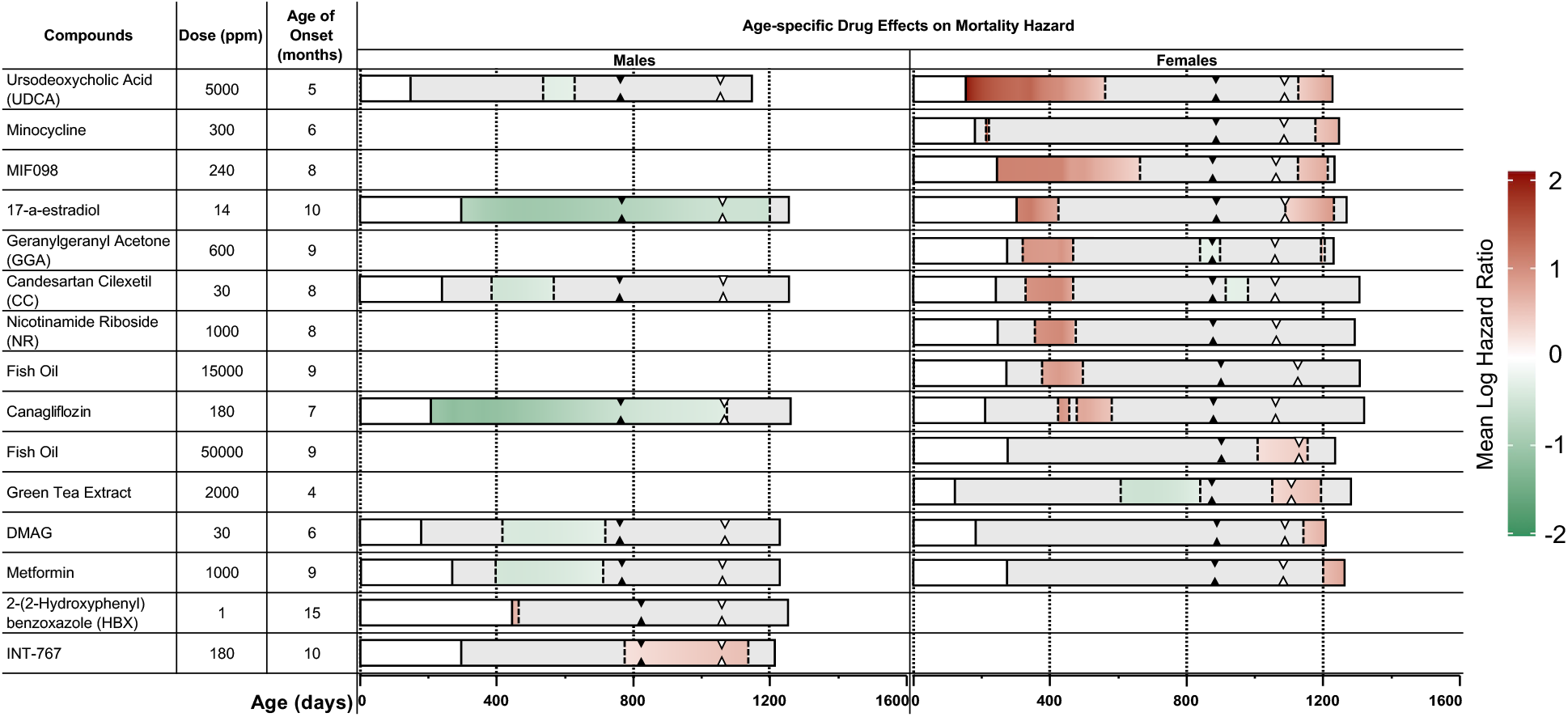
Life course heat maps of trials with drug-induced detrimental effects on mortality hazard. Only interventions confirmed by both bootstrap and asymptotic methods are displayed, with the asymptotic method results used as the representative data. These trials are shown ranked by the age when detrimental effects started in females (earliest to oldest), then by males. Other information is same as Figure 2.

**Figure S3. Mortality Hazard ratio plots for males.** Each graph shows a single intervention as well as the cohort year (the year the intervention was begun). For example, “C2017_Capt_180_5” indicates: Cohort year_compound(s)_dose (ppm)_age when treatment was initiated (month). Abbreviations are listed in Table S2. Y axis is the log mortality hazard ratio. Negative values indicate that the treatment group has lower mortality rate (beneficial).Positive values indicate the treatment group has a higher mortality rate (detrimental). Zero indicated there is no difference between treated and control groups. X axis is the age in days. The dashed line indicates the age of the treatment onset. The upper and lower dashed lines indicated the 95% confidence interval.

**Figure S4.Mortality Hazard ratio plots for females.**(see Figure S3 legend for details)

**Table S1.Interactive life course heat maps of interventions that significantly modified mortality hazard estimated by both bootstrap and asymptotic methods.**(see Figure 2 legend for details)

**Table S2.Summary of sample sizes of all groups.**

## References

1. Harrison, D. E. et al. Rapamycin fed late in life extends lifespan in genetically heterogeneous mice. Nature 460, 392–395, doi:10.1038/nature08221 (2009).

2. Harrison, D. E. et al. Acarbose, 17-alpha-estradiol, and nordihydroguaiaretic acid extend mouse lifespan preferentially in males. Aging Cell 13, 273–282, doi:10.1111/acel.12170 (2014).

3. Knufinke, M., MacArthur, M. R., Ewald, C. Y. & Mitchell, S. J. Sex differences in pharmacological interventions and their effects on lifespan and healthspan outcomes: a systematic review. Front Aging 4, 1172789, doi:10.3389/fragi.2023.1172789 (2023).

4. Mantel, N. Evaluation of survival data and two new rank order statistics arising in its consideration. Cancer Chemother Rep 50, 163–170 (1966).

5. Bouliotis, G. & Billingham, L. Crossing survival curves: alternatives to the log-rank test. Trials 12, A137, doi:10.1186/1745-6215-12-S1-A137 (2011).

6. Bradburn, M. J., Clark, T. G., Love, S. B. & Altman, D. G. Survival analysis part II: multivariate data analysis--an introduction to concepts and methods. Br J Cancer 89, 431–436, doi:10.1038/sj.bjc.6601119 (2003).

7. Jiang, N., Gelfond, J., Liu, Q., Strong, R. & Nelson, J. F. The Gehan test identifies life-extending compounds overlooked by the log-rank test in the NIA Interventions Testing Program: Metformin, Enalapril, caffeic acid phenethyl ester, green tea extract, and 17-dimethylaminoethylamino-17-demethoxygeldanamycin hydrochloride. Geroscience, doi:10.1007/s11357-024-01161-9 (2024).

8. Gehan, E. A. A Generalized Wilcoxon Test for Comparing Arbitrarily Singly-Censored Samples. Biometrika 52, 203–223 (1965).

9. Bradburn, M. J., Clark, T. G., Love, S. B. & Altman, D. G. Survival analysis Part III: multivariate data analysis -- choosing a model and assessing its adequacy and fit. Br J Cancer 89, 605–611, doi:10.1038/sj.bjc.6601120 (2003).

10. Harrington, D. P. & Fleming, T. R. A class of rank test procedures for censored survival data. Biometrika 69, 553–566 (1982).

11. Wang, C., Li, Q., Redden, D. T., Weindruch, R. & Allison, D. B. Statistical methods for testing effects on “maximum lifespan”. Mech Ageing Dev 125, 629–632, doi:10.1016/j.mad.2004.07.003 (2004).

12. Gao, G., Wan, W., Zhang, S., Redden, D. T. & Allison, D. B. Testing for differences in distribution tails to test for differences in ‘maximum’ lifespan. BMC Med Res Methodol 8, 49, doi:10.1186/1471-2288-8-49 (2008).

13. Pletcher, S. D., Khazaeli, A. A. & Curtsinger, J. W. Why do life spans differ? Partitioning mean longevity differences in terms of age-specific mortality parameters. J Gerontol A Biol Sci Med Sci 55, B381–389, doi:10.1093/gerona/55.8.b381 (2000).

14. Zhang, Z., Reinikainen, J., Adeleke, K. A., Pieterse, M. E. & Groothuis-Oudshoorn, C. G. Time-varying covariates and coefficients in Cox regression models. Annals of translational medicine 6 (2018).

15. Gilbert, P. B., Wei, L. J., Kosorok, M. R. & Clemens, J. D. Simultaneous inferences on the contrast of two hazard functions with censored observations. Biometrics 58, 773–780, doi:10.1111/j.0006-341x.2002.00773.x (2002).

16. Swindell, W. R. Accelerated failure time models provide a useful statistical framework for aging research. Exp Gerontol 44, 190–200, doi:10.1016/j.exger.2008.10.005 (2009).

17. Wei, L. J. The accelerated failure time model: a useful alternative to the Cox regression model in survival analysis. Stat Med 11, 1871–1879, doi:10.1002/sim.4780111409 (1992).

18. Royston, P. & Parmar, M. K. Restricted mean survival time: an alternative to the hazard ratio for the design and analysis of randomized trials with a time-to-event outcome. BMC Med Res Methodol 13, 152, doi:10.1186/1471-2288-13-152 (2013).

19. Liao, J. J. Z., Liu, G. F. & Wu, W. C. Dynamic RMST curves for survival analysis in clinical trials. BMC Med Res Methodol 20, 218, doi:10.1186/s12874-020-01098-5 (2020).

20. Kronborg, D. & Aaby, P. Piecewise comparison of survival functions in stratified proportional hazards models. Biometrics 46, 375–380 (1990).

21. Fisher, L. D. & Lin, D. Y. Time-dependent covariates in the Cox proportional-hazards regression model. Annu Rev Public Health 20, 145–157, doi:10.1146/annurev.publhealth.20.1.145 (1999).

22. Gould, S. J. The median isn’t the message. AMA Journal of Ethics 15, 77–81 (2013).

23. Rebora, P., Salim, A. & Reilly, M. Bshazard: a flexible tool for nonparametric smoothing of the hazard function. The R Journal 6, 114–122 (2014).

24. Van der Vaart, A. W. Asymptotic statistics. Vol. 3 (Cambridge university press, 2000).

25. Efron, B. The jackknife, the bootstrap and other resampling plans. (SIAM, 1982).

26. Efron, B. & Tibshirani, R. J. An introduction to the bootstrap. (Chapman and Hall/CRC, 1994).

27. Miller, R. A. et al. An Aging Interventions Testing Program: study design and interim report. Aging Cell 6, 565–575, doi:10.1111/j.1474-9726.2007.00311.x (2007).

28. Jiang, N. & Nelson, J. F. Sex differences in mouse longevity and responses to geroprotective drugs: implications for human intervention. Public Policy Aging Rep 33, 120–124, doi:10.1093/ppar/prad026 (2023).

29. Strong, R. et al. Evaluation of resveratrol, green tea extract, curcumin, oxaloacetic acid, and medium-chain triglyceride oil on life span of genetically heterogeneous mice. J Gerontol A Biol Sci Med Sci 68, 6–16, doi:10.1093/gerona/gls070 (2013).

30. Jiang, N. et al. Prepubertal castration eliminates sex differences in lifespan and growth trajectories in genetically heterogeneous mice. Aging Cell 22, e13891, doi:10.1111/acel.13891 (2023).

31. Cheng, C. J., Gelfond, J. A. L., Strong, R. & Nelson, J. F. Genetically heterogeneous mice exhibit a female survival advantage that is age- and site-specific: Results from a large multi-site study. Aging Cell 18, e12905, doi:10.1111/acel.12905 (2019).

32. Miller, R. A. et al. Lifespan effects in male UM-HET3 mice treated with sodium thiosulfate, 16-hydroxyestriol, and late-start canagliflozin. Geroscience, doi:10.1007/s11357-024-01176-2 (2024).

33. Yu, B. P., Masoro, E. J. & McMahan, C. A. Nutritional influences on aging of Fischer 344 rats: I. Physical, metabolic, and longevity characteristics. J Gerontol 40, 657–670, doi:10.1093/geronj/40.6.657 (1985).

34. Masoro, E. J. Biology of Aging: Current State of Knowledge. Archives of Internal Medicine 147, 166–169, doi:10.1001/archinte.1987.00370010164033 (1987).

35. Jiang, N., Cheng, C. J., Strong, R. & Nelson, J. F. Castration reduces mortality and increases resilience in male mice: what is next? Geroscience 46, 2787–2790, doi:10.1007/s11357-023-00973-5 (2024).

36. Miller, R. A. et al. Correction to: Lifespan effects in male UM-HET3 mice treated with sodium thiosulfate, 16-hydroxyestriol, and late-start canagliflozin. Geroscience, doi:10.1007/s11357-024-01283-0 (2024).

37. Levine, M. E. et al. Low protein intake is associated with a major reduction in IGF-1, cancer, and overall mortality in the 65 and younger but not older population. Cell Metab 19, 407–417, doi:10.1016/j.cmet.2014.02.006 (2014).

38. Richardson, N. E. et al. Lifelong restriction of dietary branched-chain amino acids has sex-specific benefits for frailty and lifespan in mice. Nat Aging 1, 73–86, doi:10.1038/s43587-020-00006-2 (2021).

39. Miller, R. A. et al. Canagliflozin extends life span in genetically heterogeneous male but not female mice. JCI Insight 5, doi:10.1172/jci.insight.140019 (2020).

40. Strong, R. et al. Lifespan benefits for the combination of rapamycin plus acarbose and for captopril in genetically heterogeneous mice. Aging Cell 21, e13724, doi:10.1111/acel.13724 (2022).

