## Supplemental Figure 4 for "Deciphering the Timing and Impact of Life-extending Interventions: Temporal Efficacy Profiler Distinguishes Early, Midlife, and Senescence Phase Efficacies"

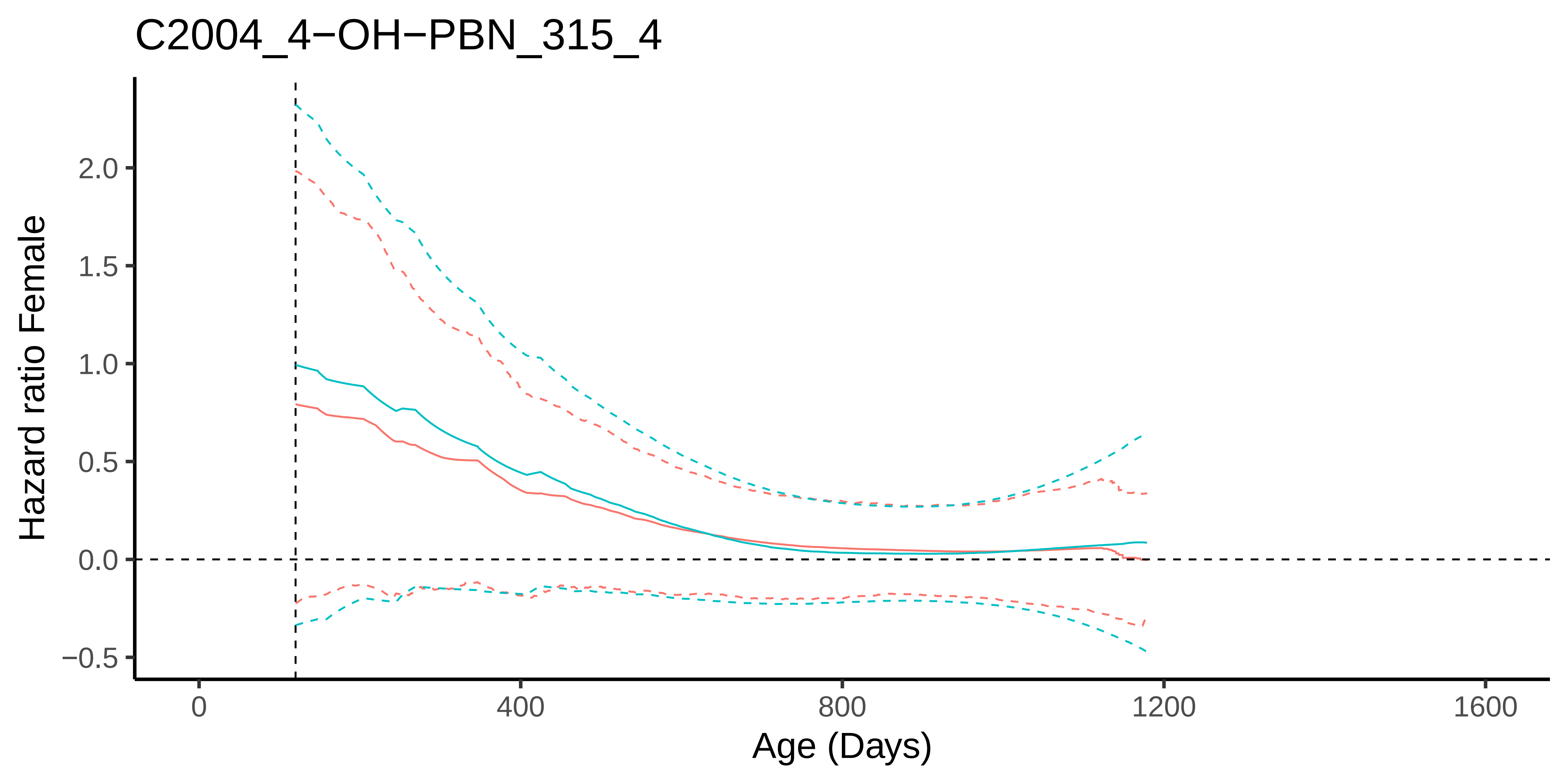
C2004_4−OH−PBN_315_4
Hazard ratio Female
2.0
1.5
1.0
0.5
0.0
−0.5
0
400
800
1200
1600
Age (Days)

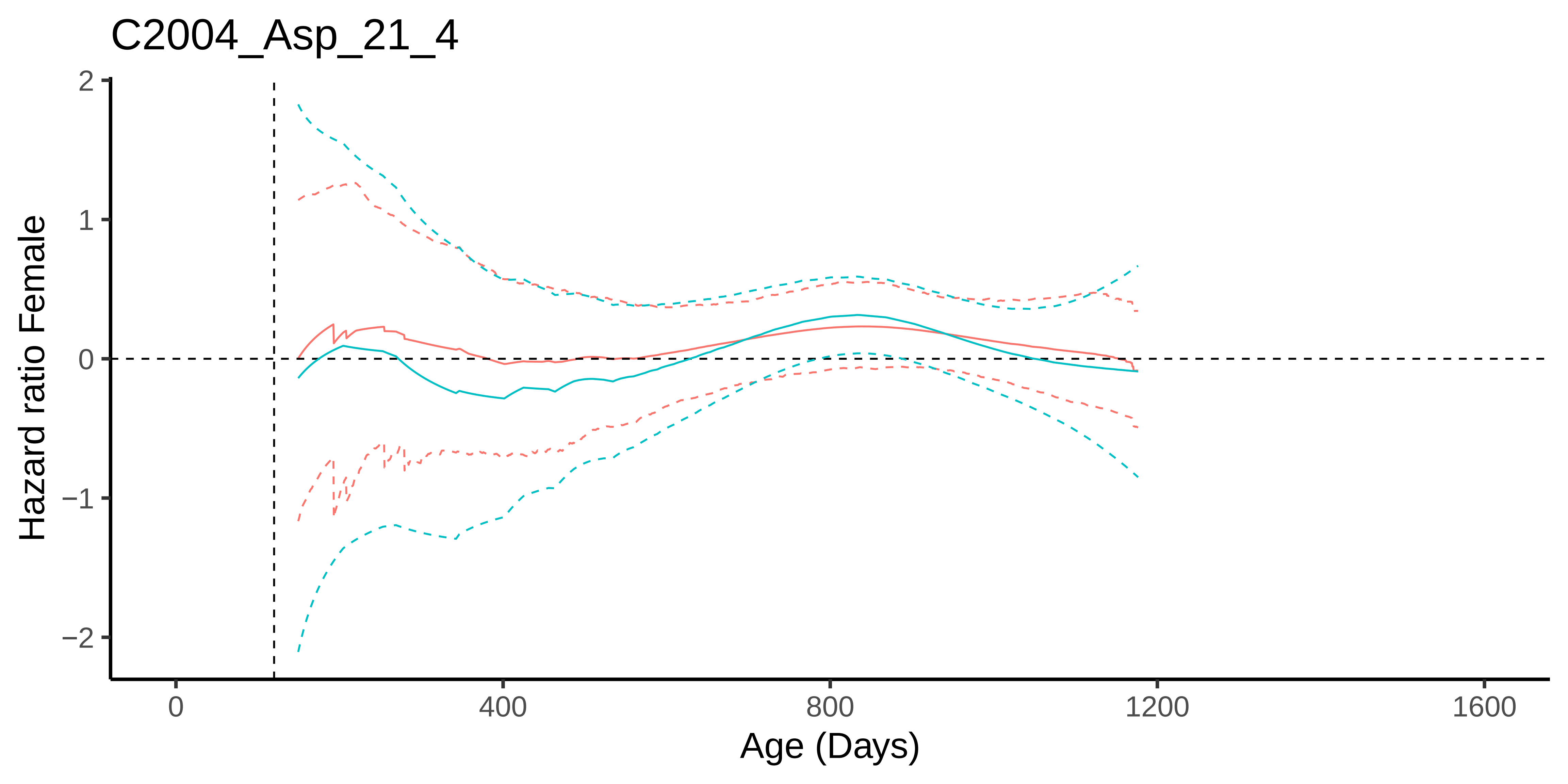
C2004_Asp_21_4
Hazard ratio Female
2
1
0
−1
−2
0
400
800
1200
1600
Age (Days)

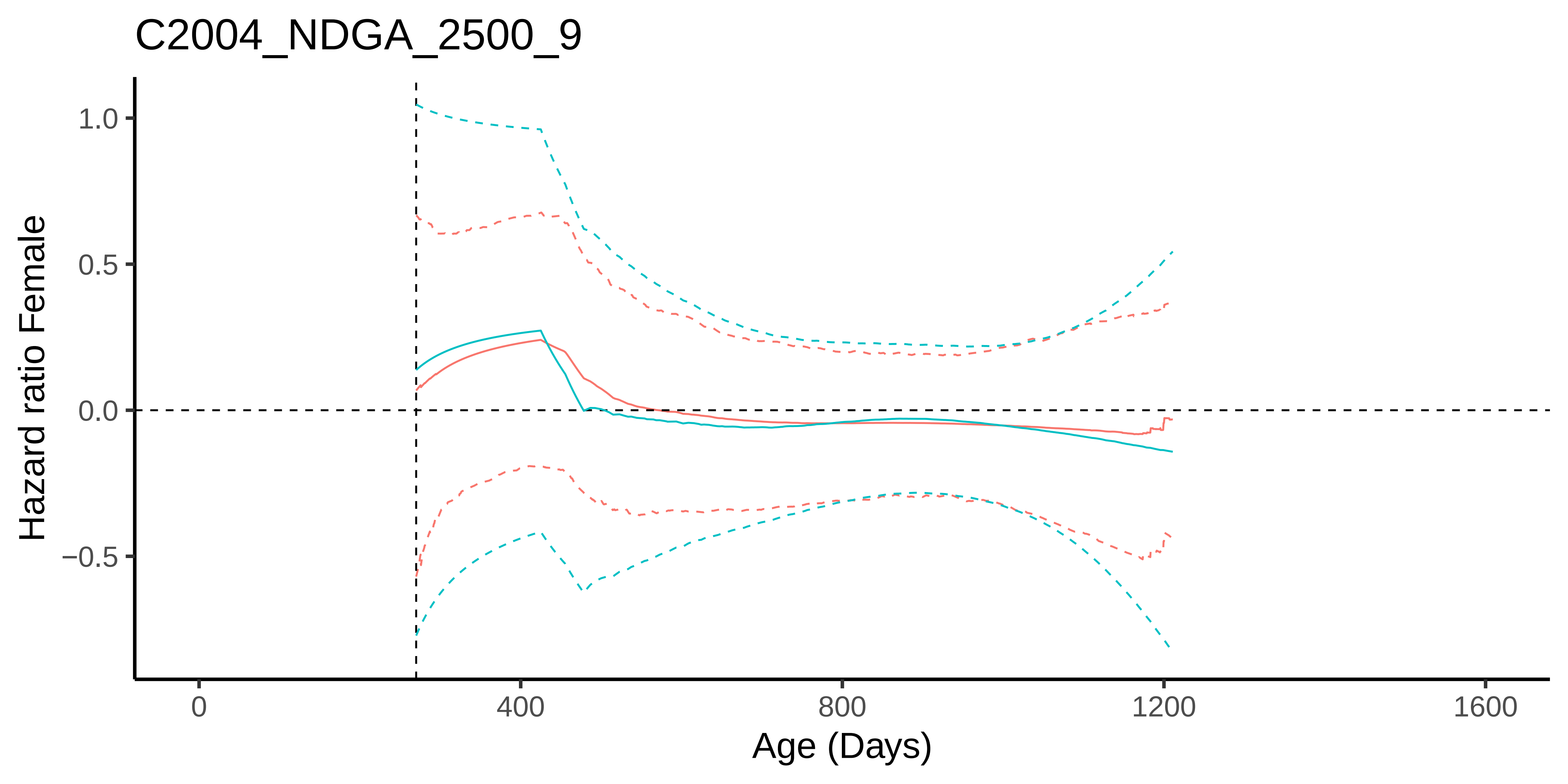
C2004_NDGA_2500_9
Hazard ratio Female
1.0
0.5
0.0
−0.5
0
400
800
1200
1600
Age (Days)

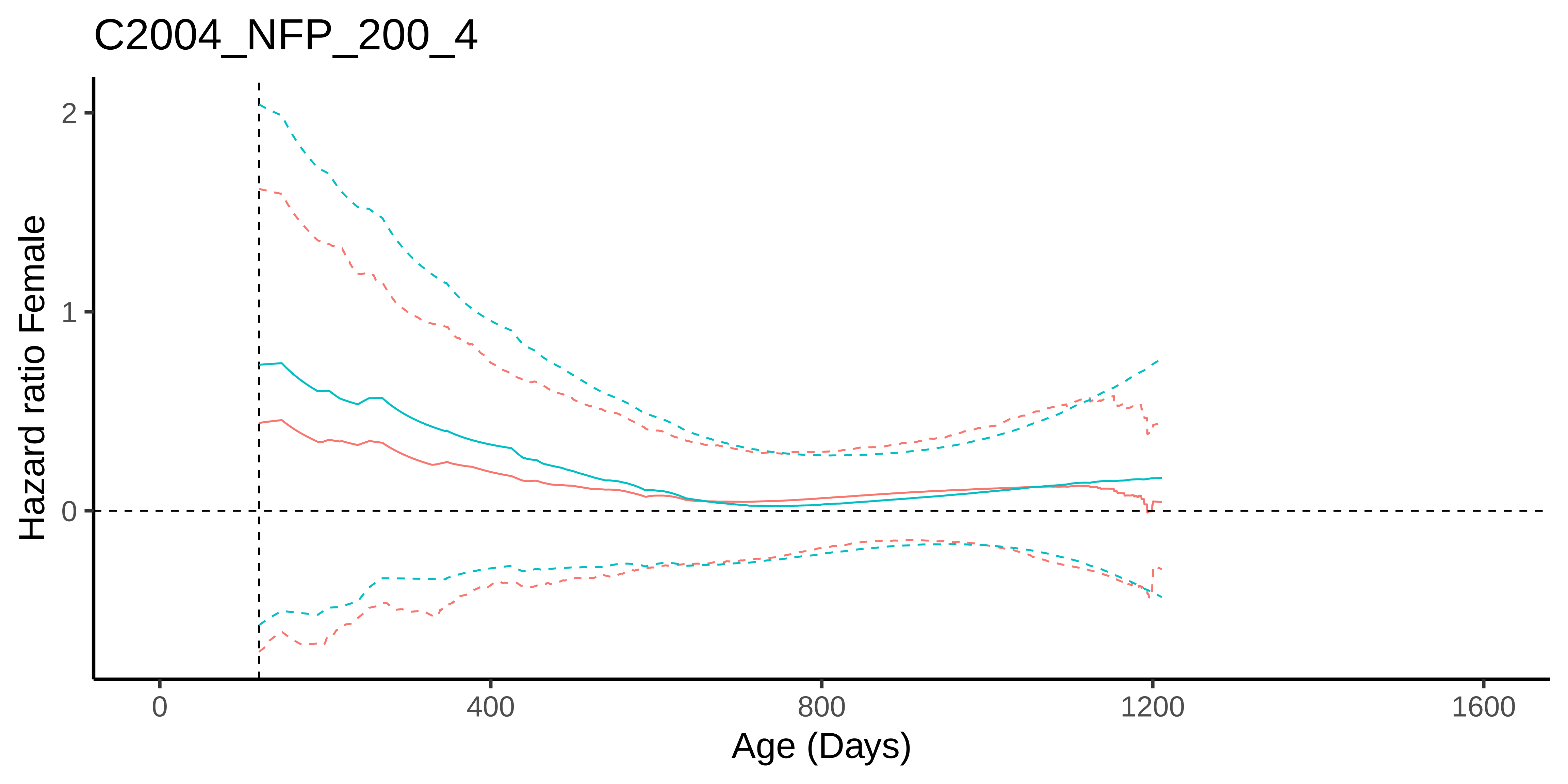
C2004_NFP_200_4
Hazard ratio Female
2
1
0
0
400
800
1200
1600
Age (Days)

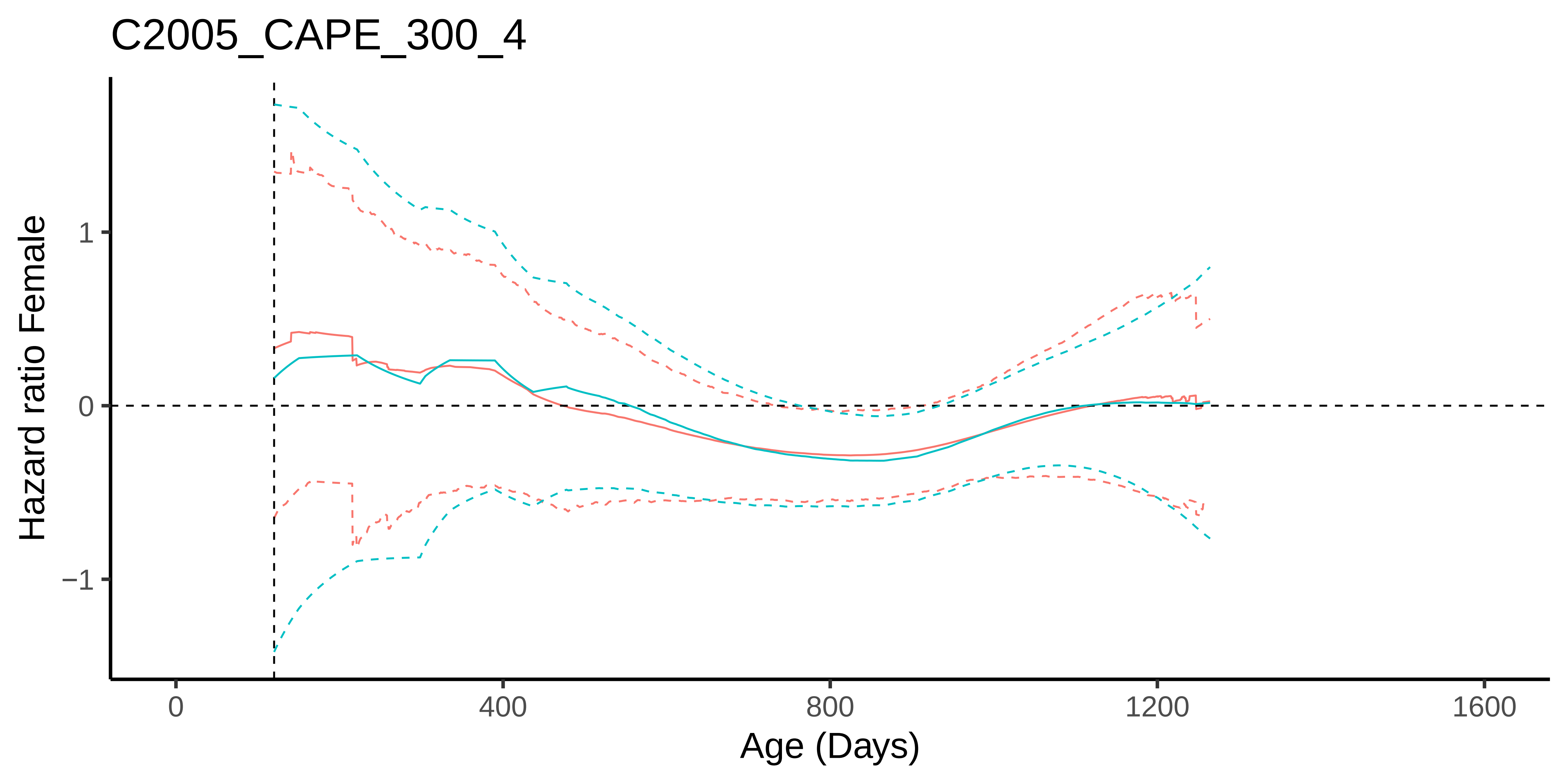
C2005_CAPE_300_4
Hazard ratio Female
1
0
−1
0
400
800
1200
1600
Age (Days)

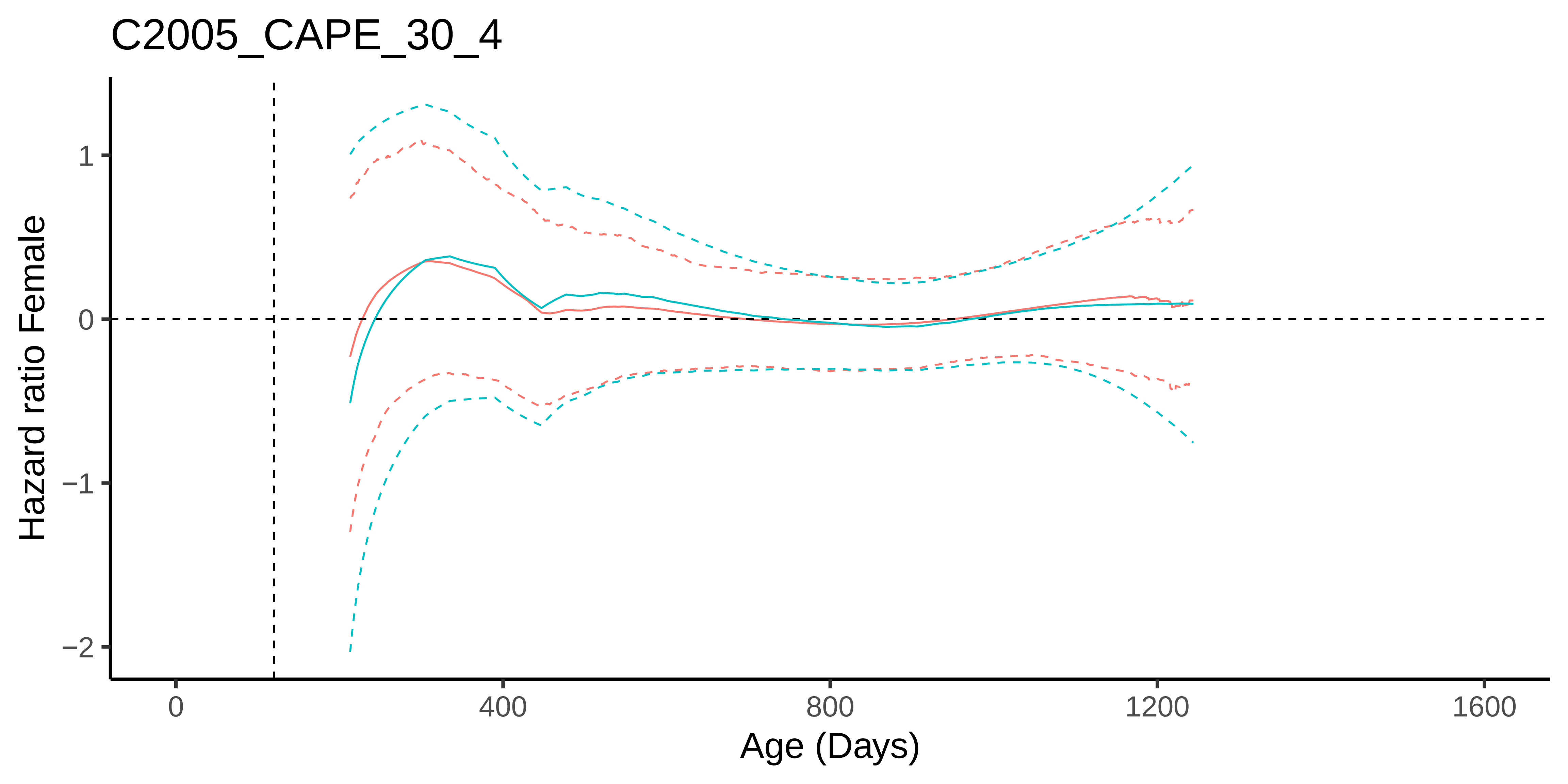

C2005_CAPE_30_4
Hazard ratio Female
1
0
−1
−2
0
400
800
1200
1600
Age (Days)

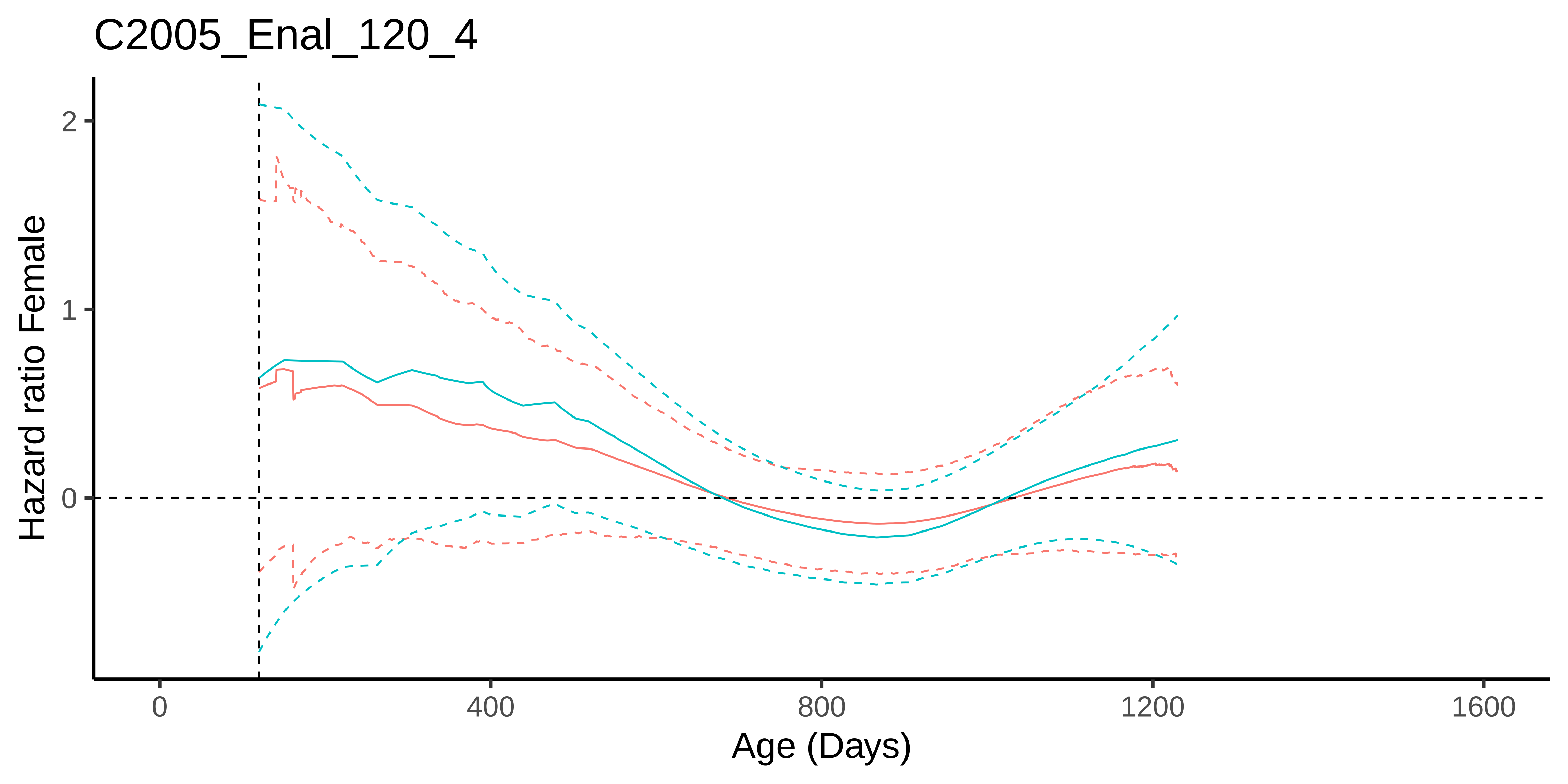
C2005_Enal_120_4
Hazard ratio Female
2
1
0
0
400
800
1200
1600
Age (Days)

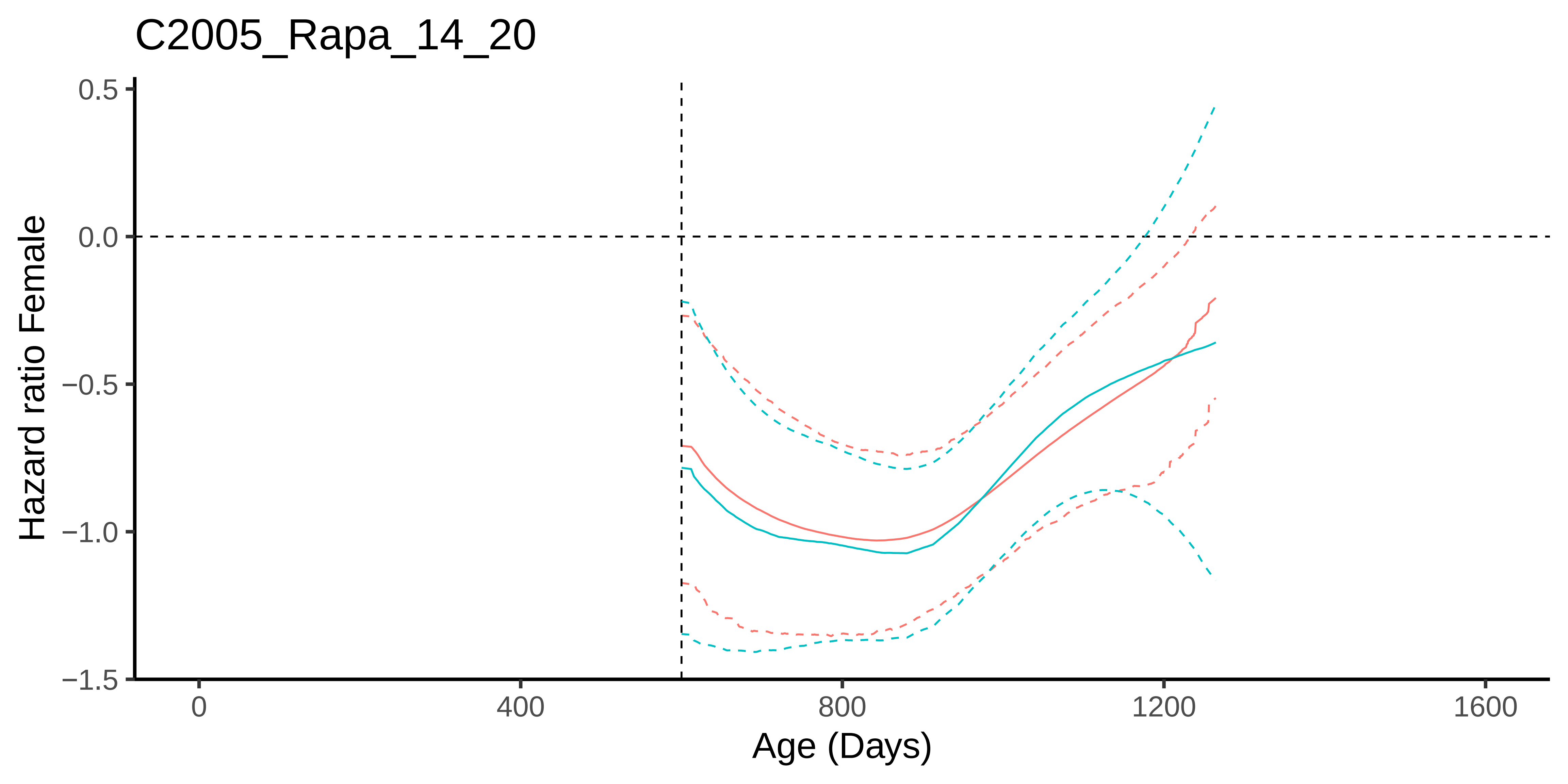

C2005_Rapa_14_20
Hazard ratio Female
0.5
0.0
−0.5
−1.0
−1.5
0
400
800
1200
1600
Age (Days)

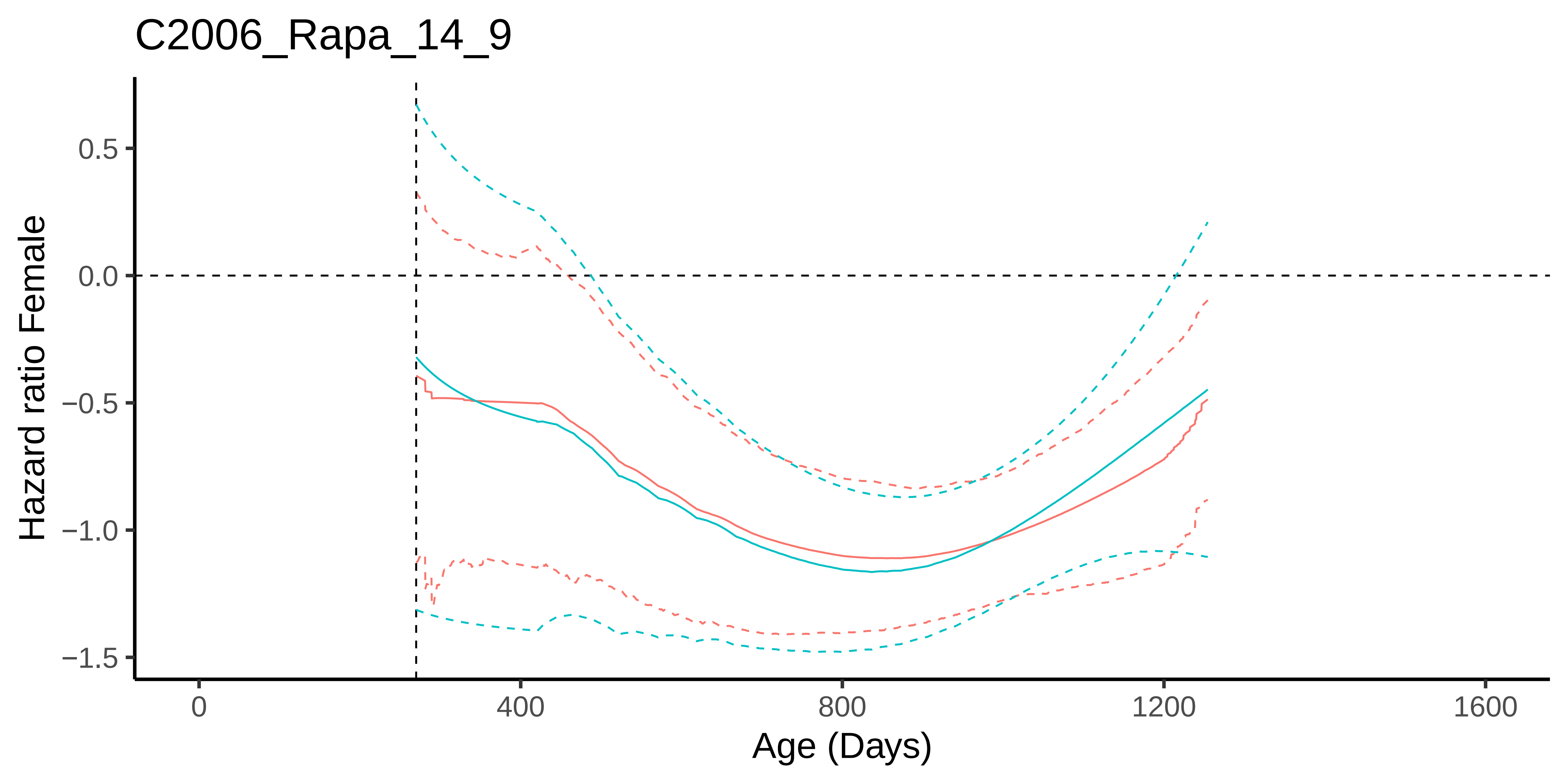

C2006_Rapa_14_9
Hazard ratio Female
0.5
0.0
−0.5
−1.0
−1.5
0
400
800
1200
1600
Age (Days)

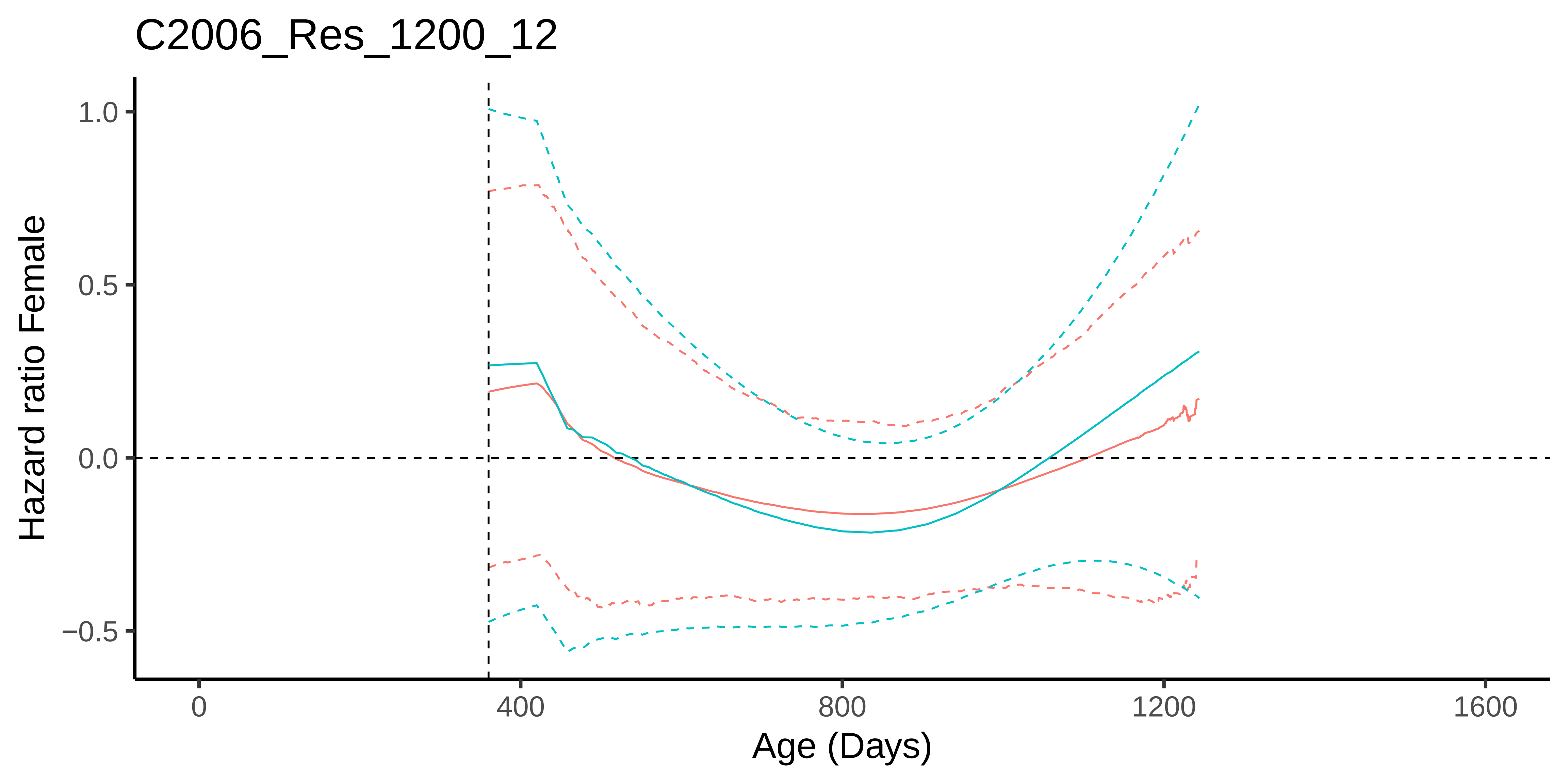

C2006_Res_1200_12
Hazard ratio Female
1.0
0.5
0.0
−0.5
0
400
800
1200
1600
Age (Days)

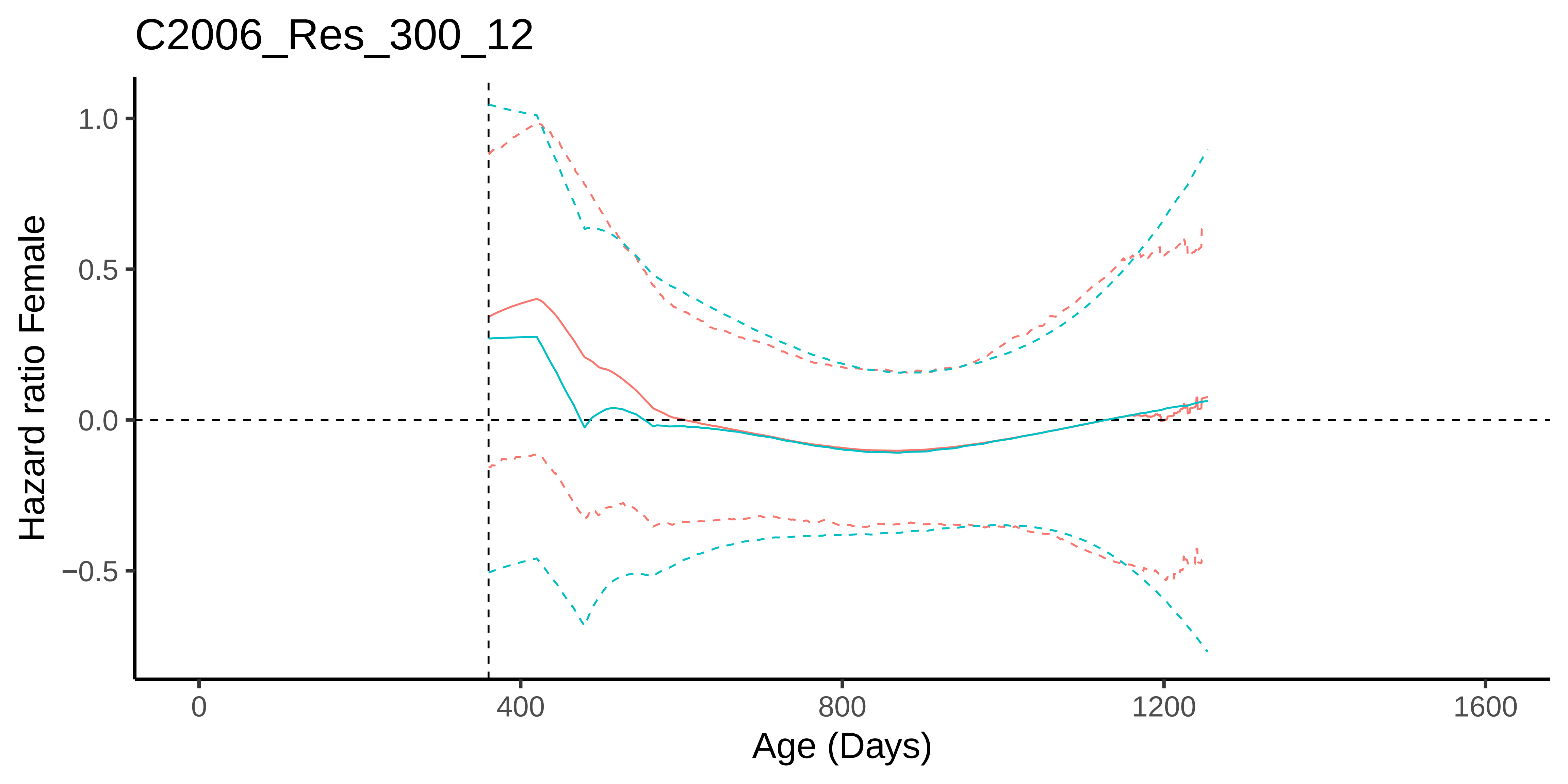
C2006_Res_300_12
Hazard ratio Female
1.0
0.5
0.0
−0.5
0
400
800
1200
1600
Age (Days)

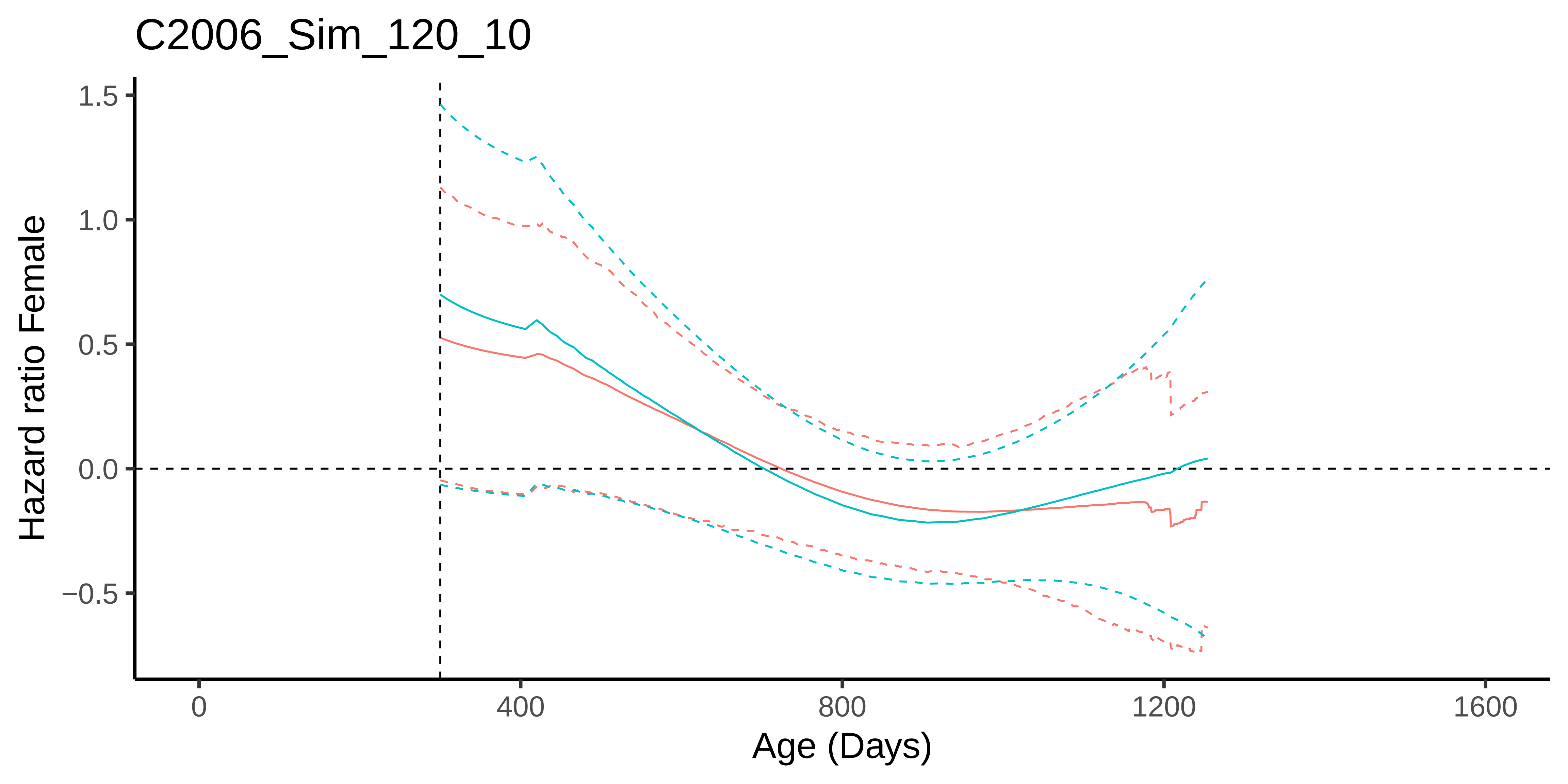

C2006_Sim_120_10
Hazard ratio Female
1.5
1.0
0.5
0.0
−0.5
0
400
800
1200
1600
Age (Days)

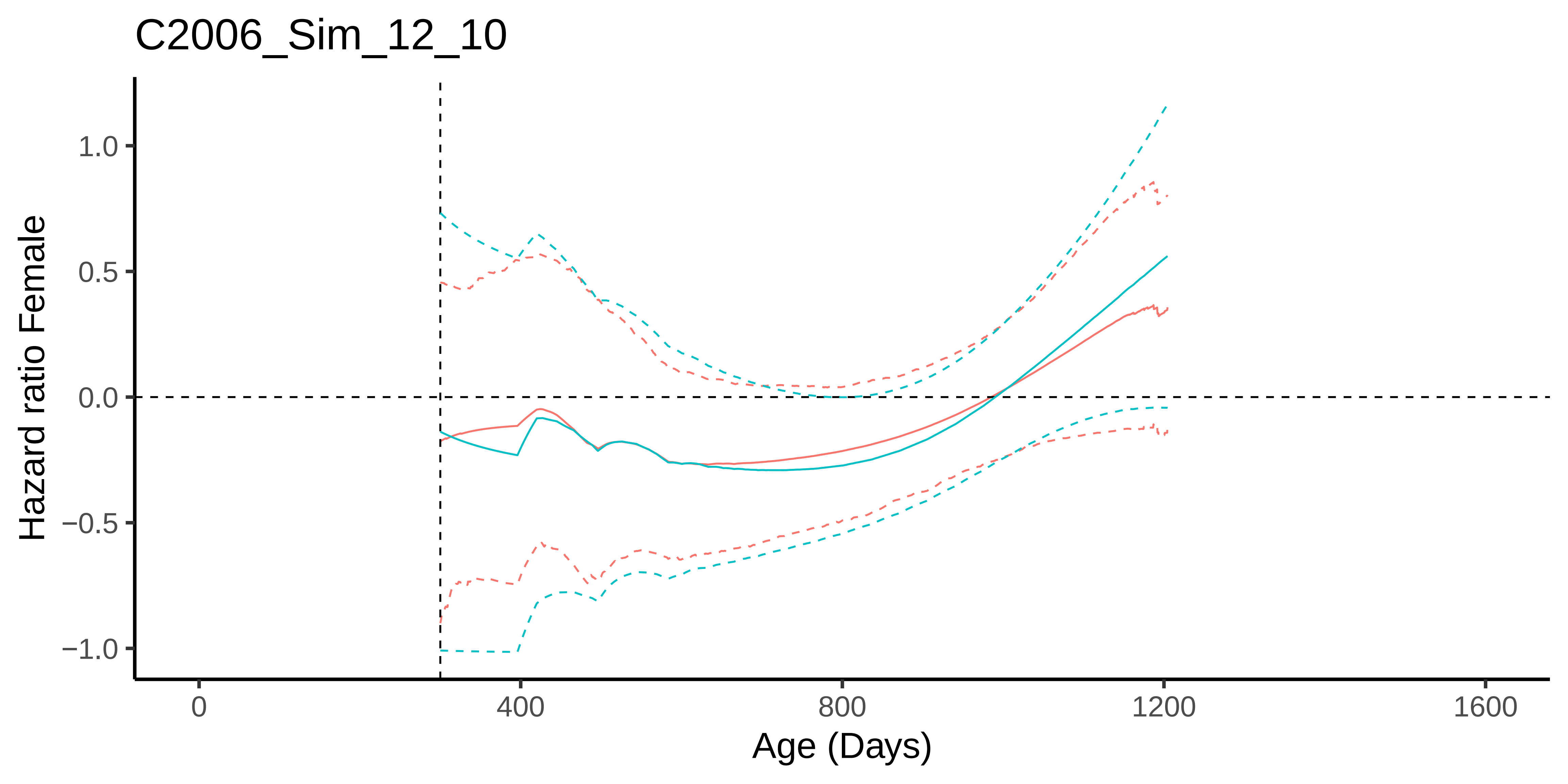

C2006_Sim_12_10
Hazard ratio Female
1.0
0.5
0.0
−0.5
−1.0
0
400
800
1200
1600
Age (Days)

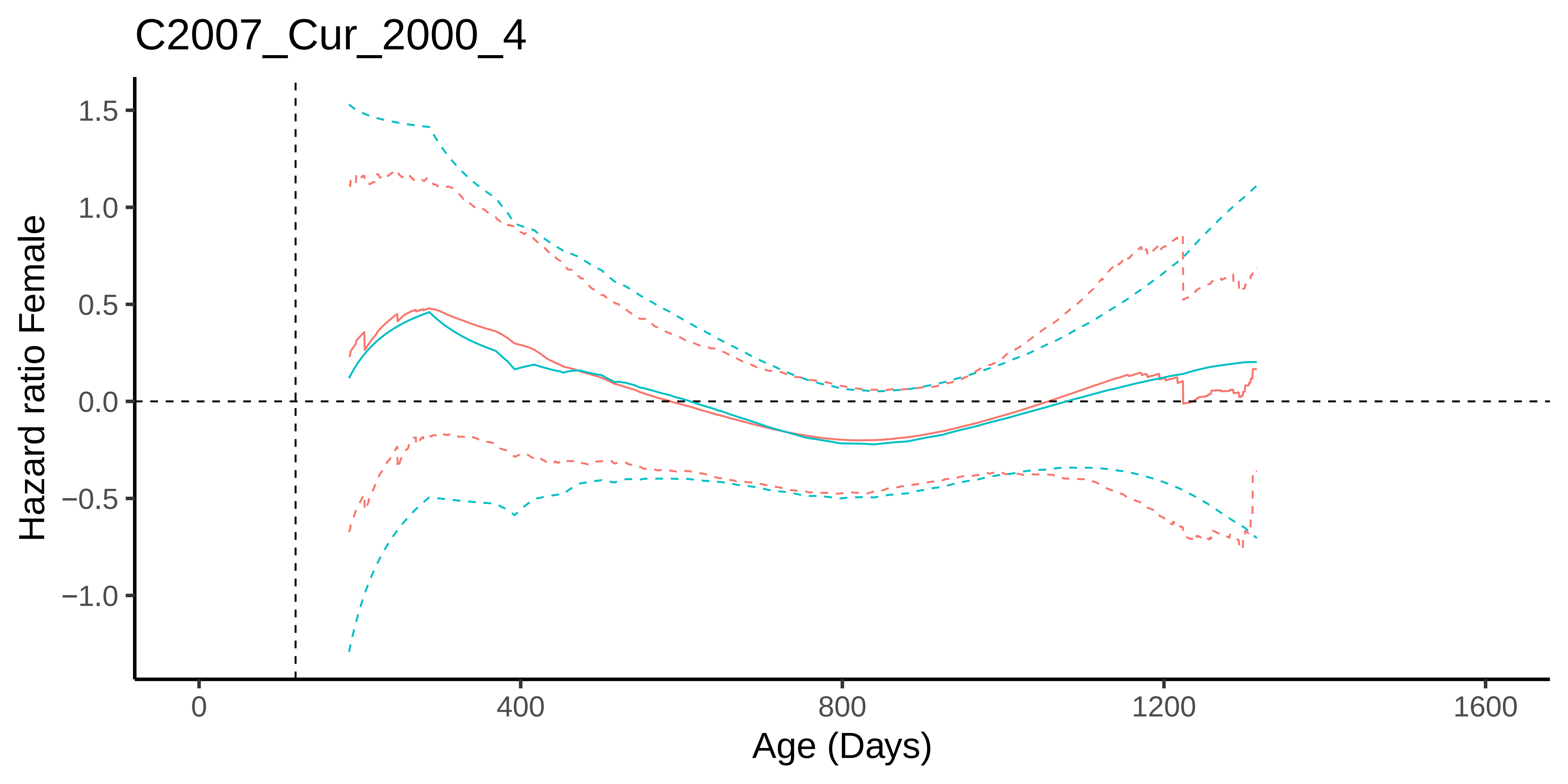
C2007_Cur_2000_4
Hazard ratio Female
1.5
1.0
0.5
0.0
−0.5
−1.0
0
400
800
1200
1600
Age (Days)

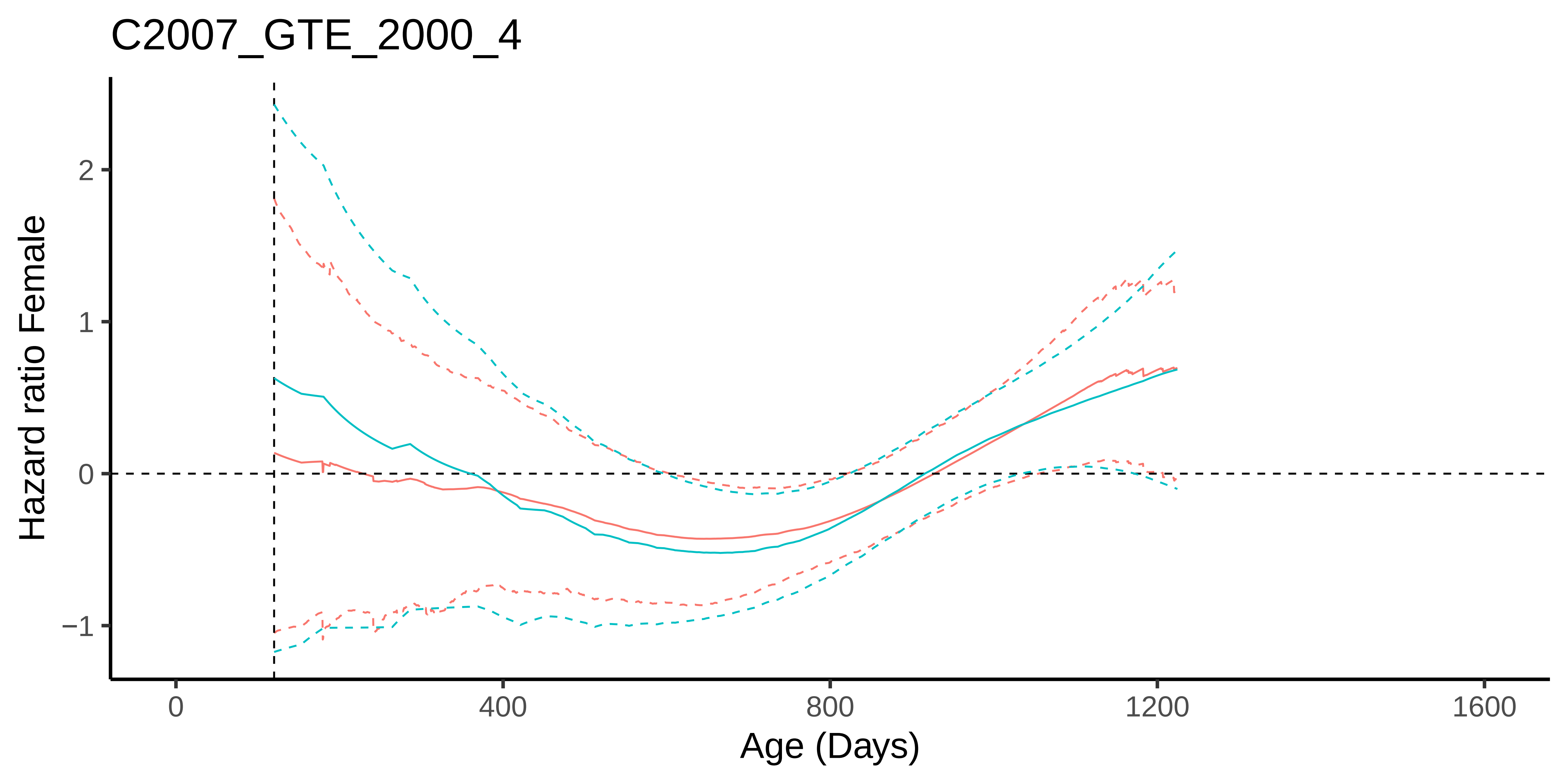
C2007_GTE_2000_4
Hazard ratio Female
2
1
0
−1
0
400
800
1200
1600
Age (Days)

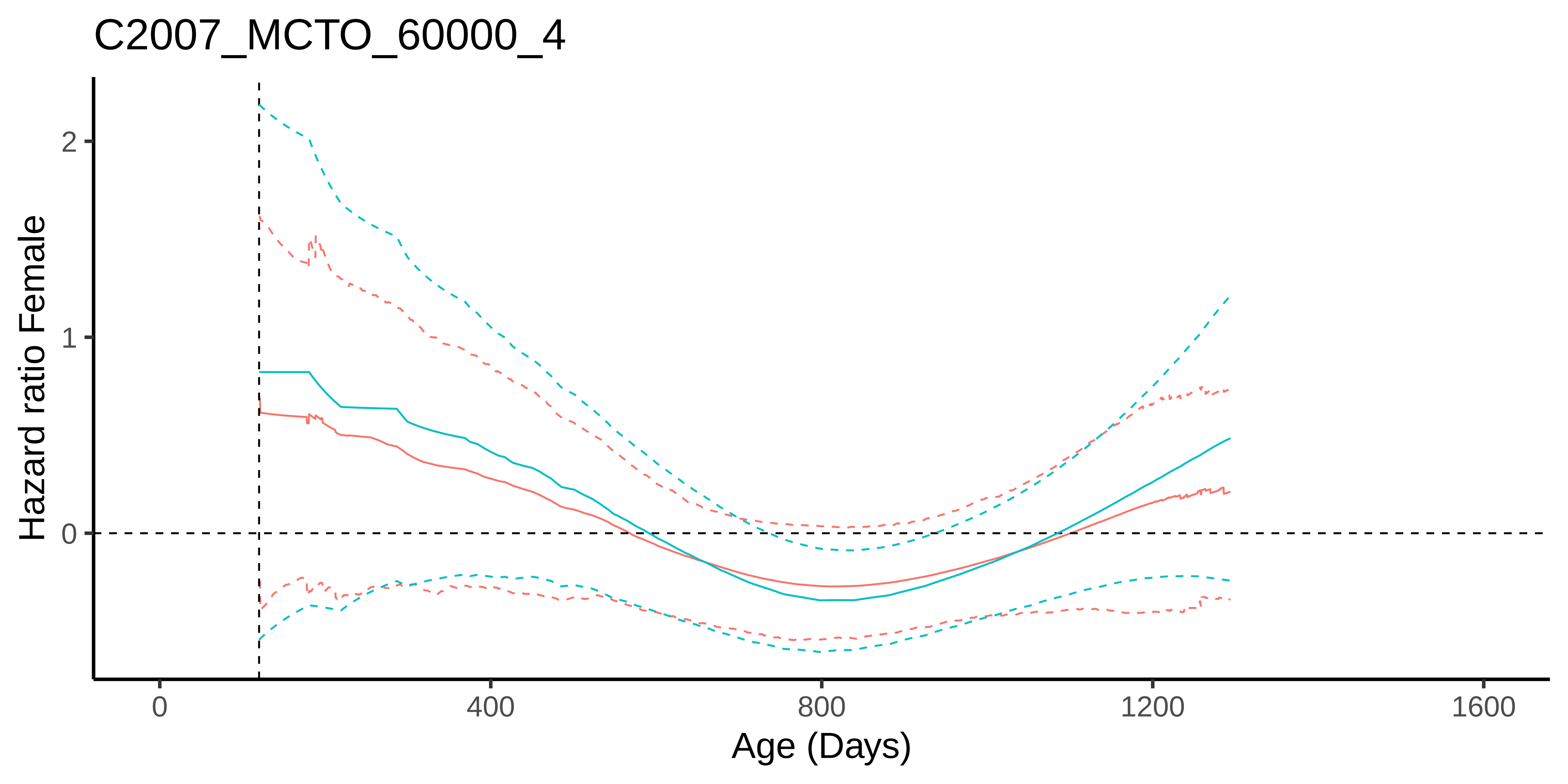
C2007_MCTO_60000_4
Hazard ratio Female
2
1
0
0
400
800
1200
1600
Age (Days)

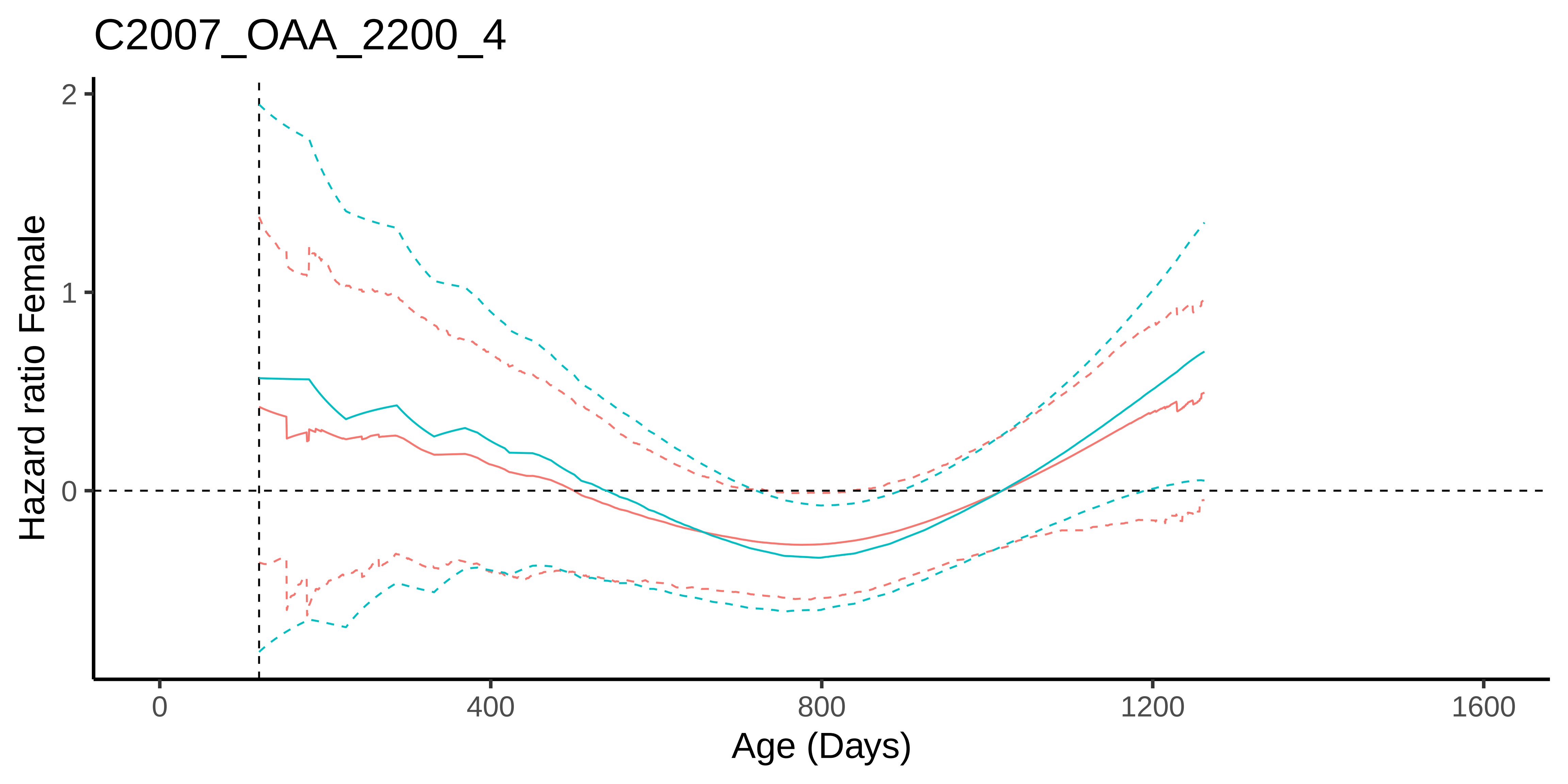
C2007_OAA_2200_4
Hazard ratio Female
2
1
0
0
400
800
1200
1600
Age (Days)

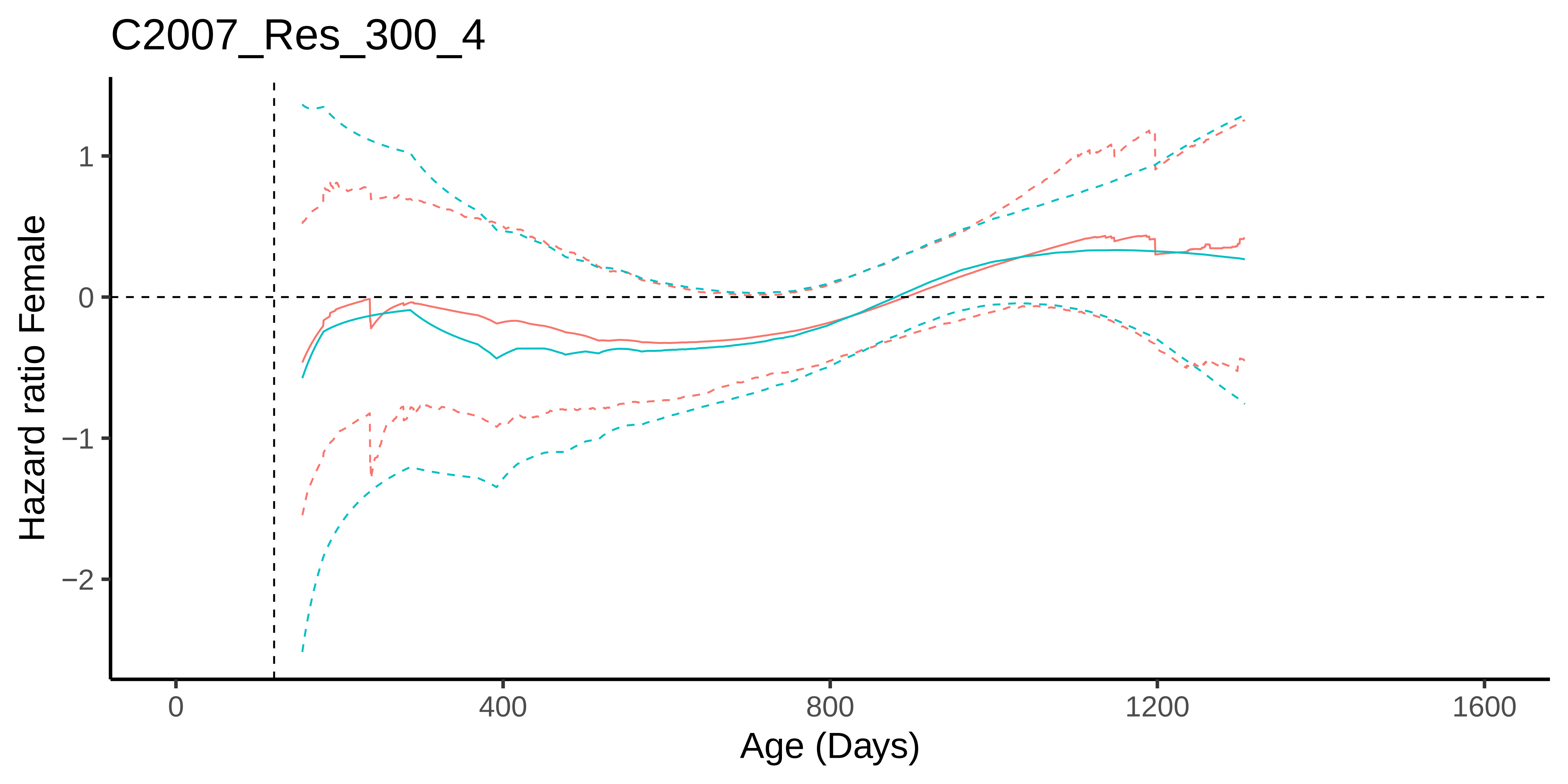
C2007_Res_300_4
Hazard ratio Female
1
0
−1
−2
0
400
800
1200
1600
Age (Days)

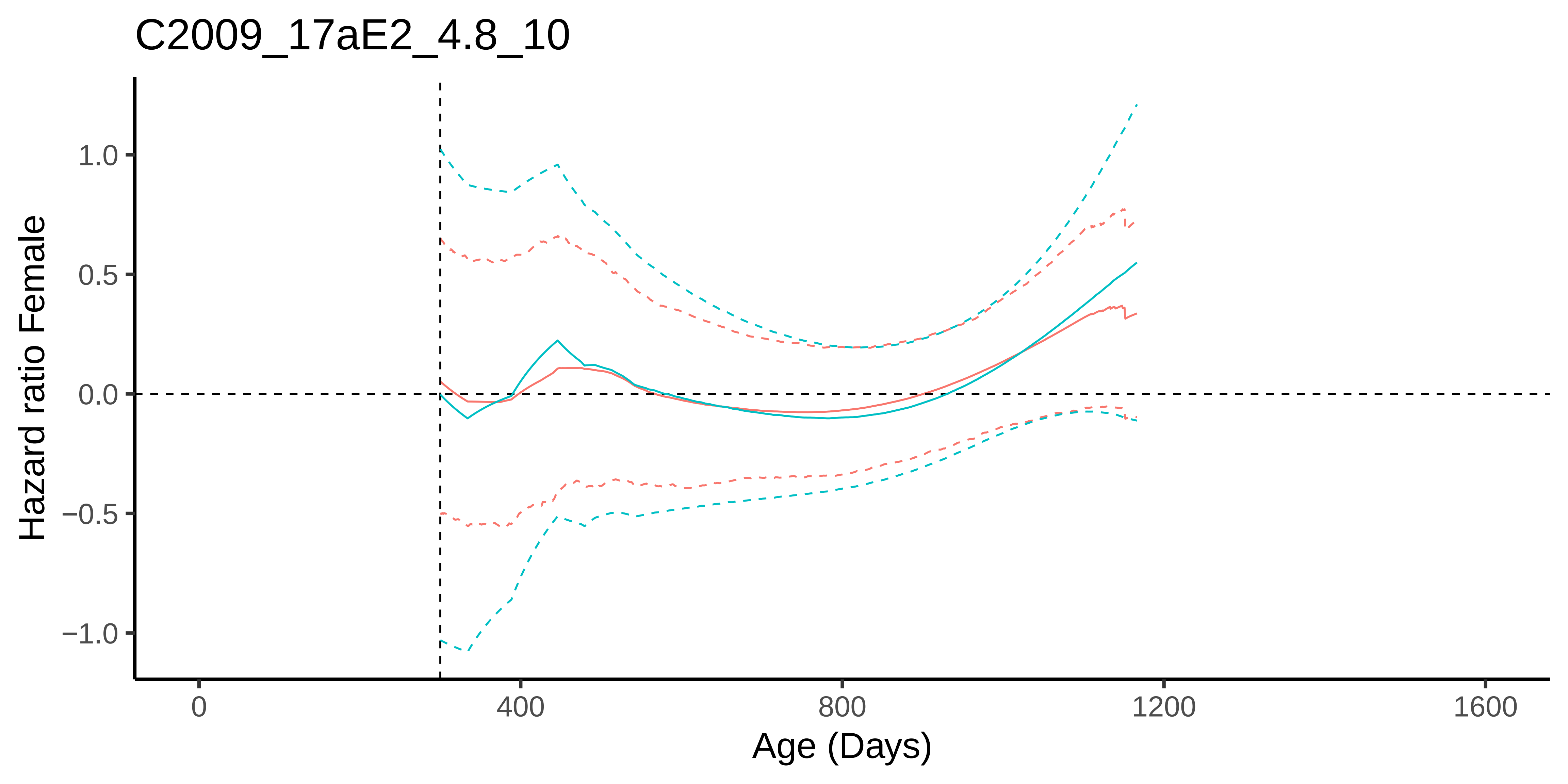

C2009_17aE2_4.8_10
Hazard ratio Female
1.0
0.5
0.0
−0.5
−1.0
0
400
800
1200
1600
Age (Days)

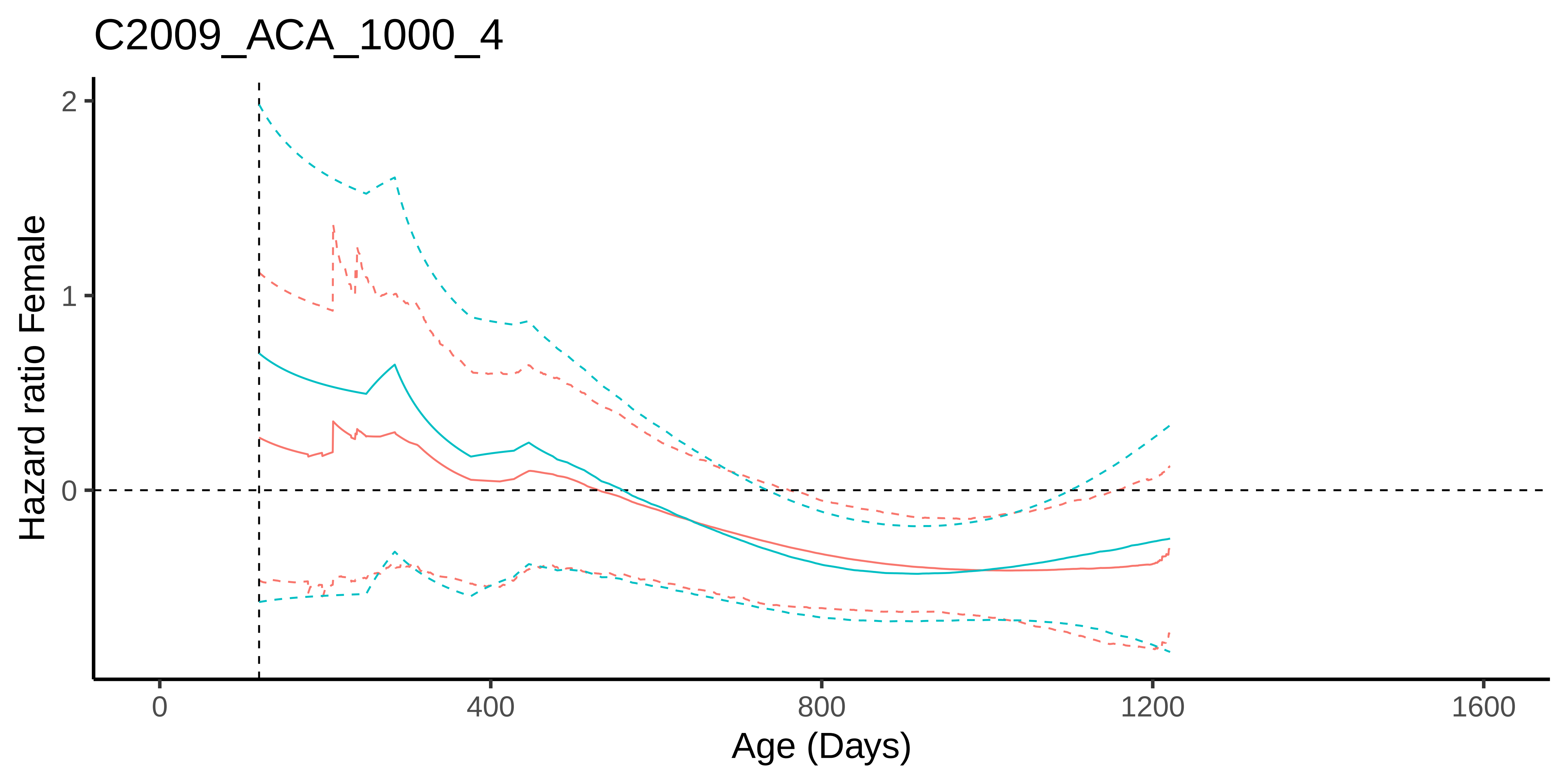
C2009_ACA_1000_4
Hazard ratio Female
2
1
0
0
400
800
1200
1600
Age (Days)

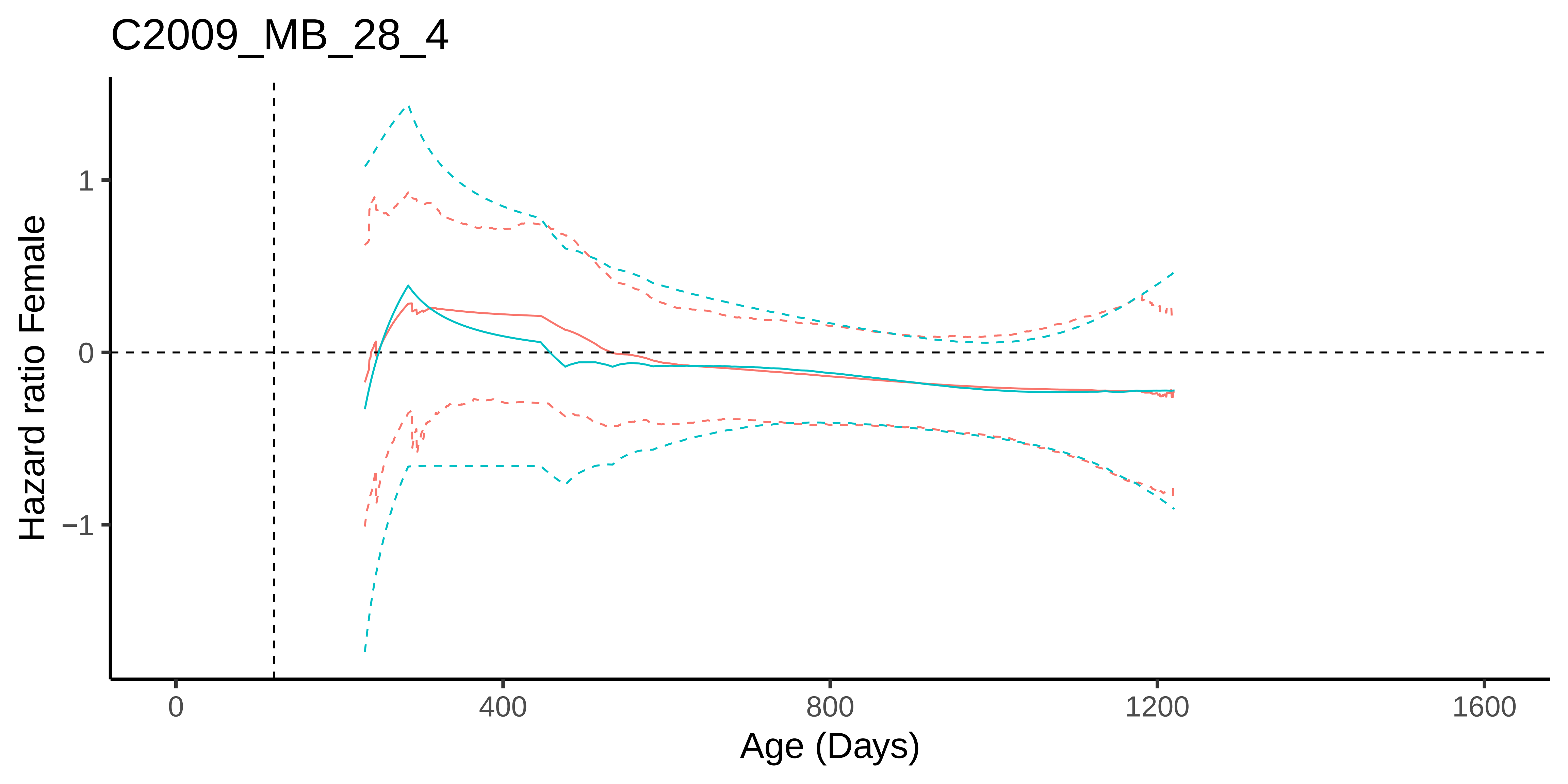
C2009_MB_28_4
Hazard ratio Female
1
0
−1
0
400
800
1200
1600
Age (Days)

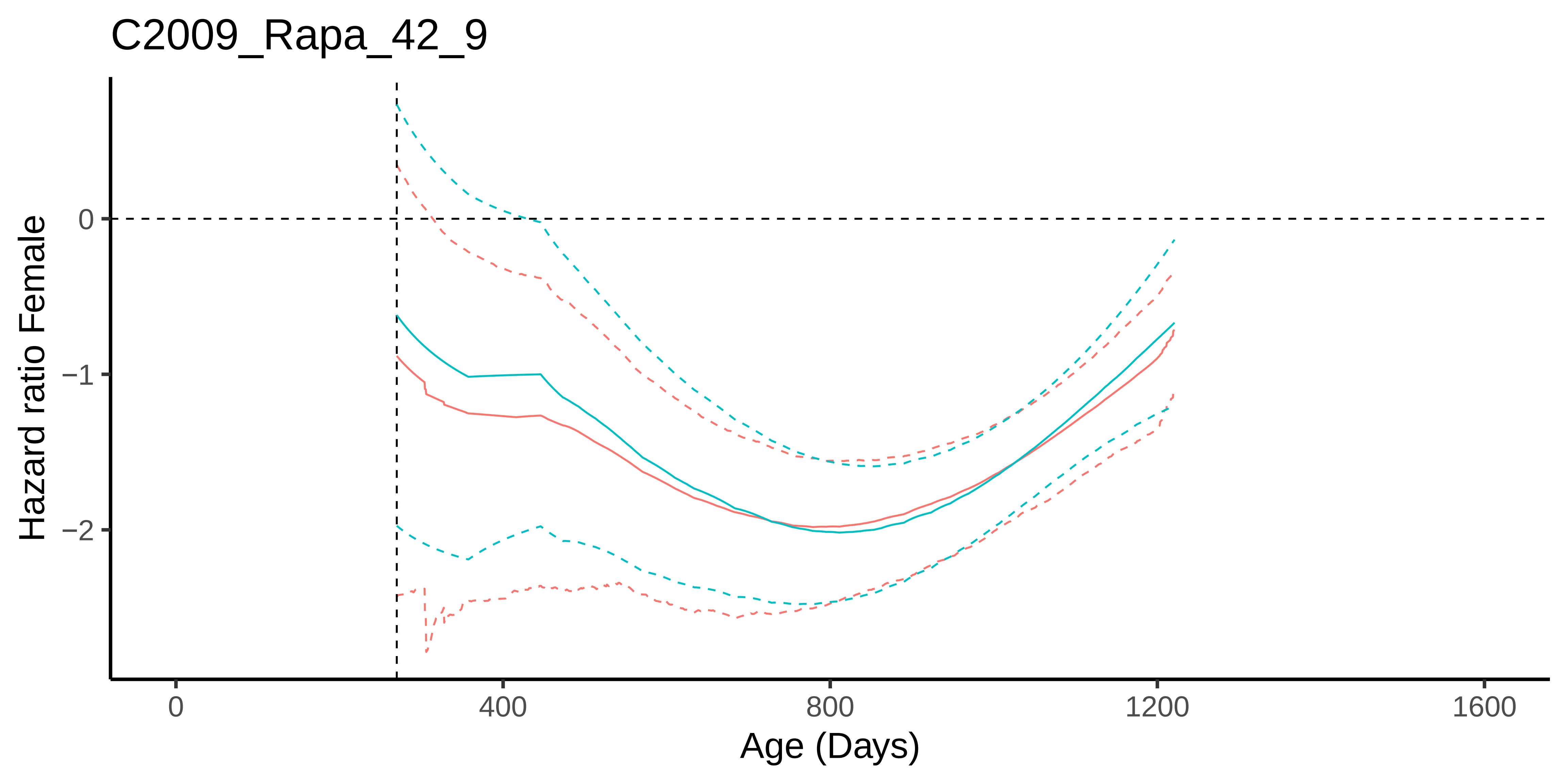
C2009_Rapa_42_9
Hazard ratio Female
0
−1
−2
0
400
800
1200
1600
Age (Days)

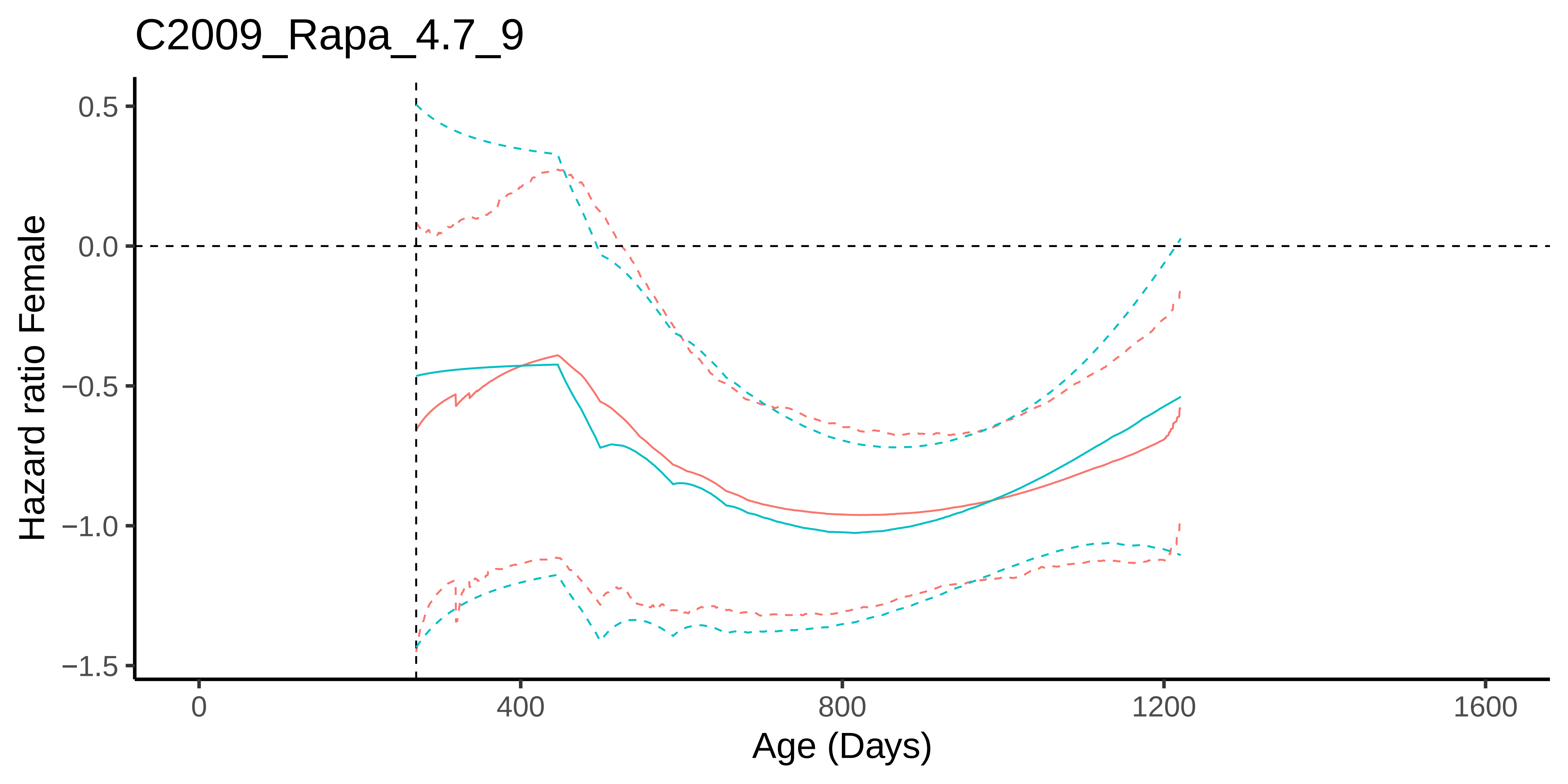
C2009_Rapa_4.7_9
Hazard ratio Female
0.5
0.0
−0.5
−1.0
−1.5
0
400
800
1200
1600
Age (Days)

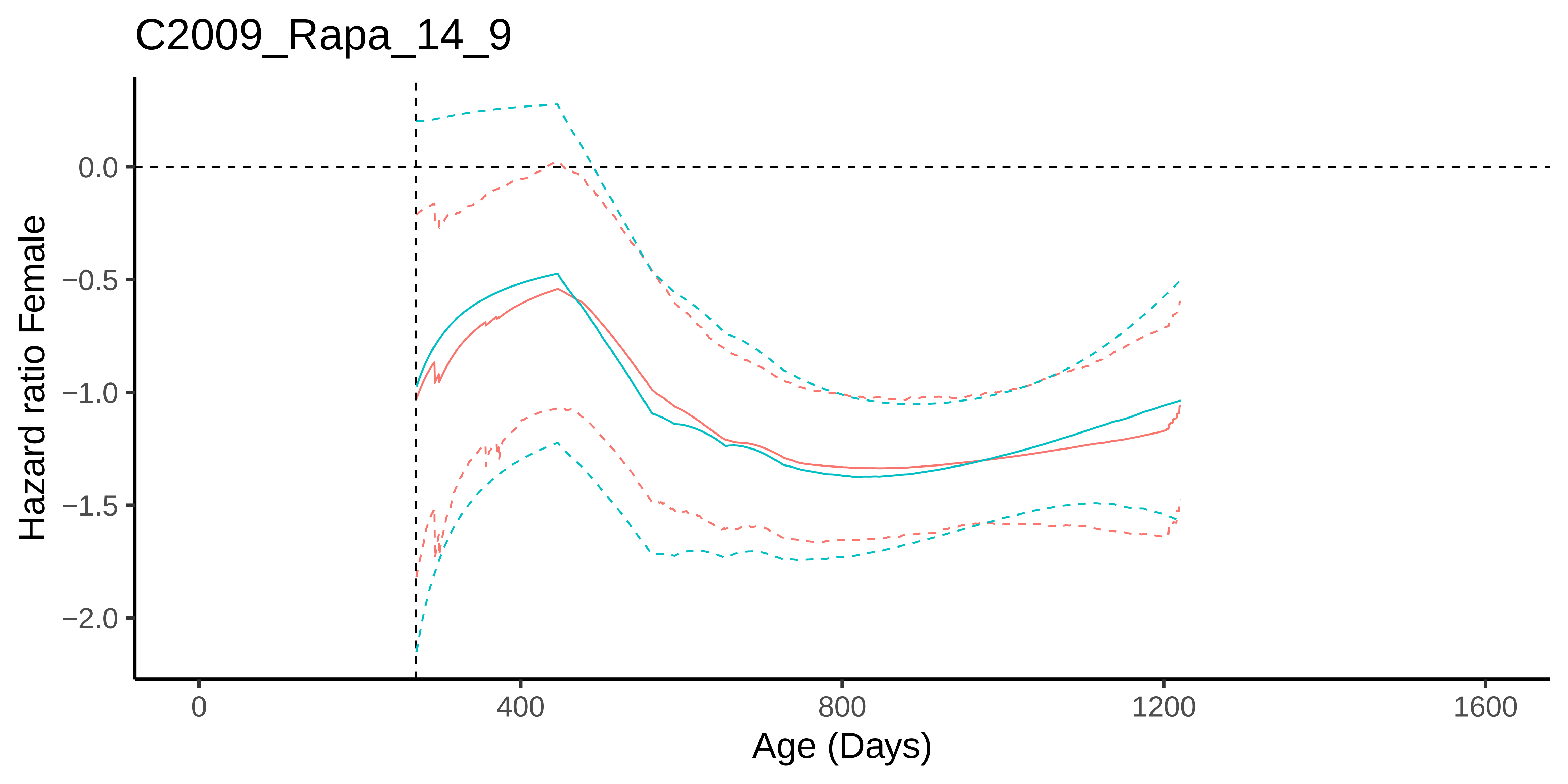
C2009_Rapa_14_9
Hazard ratio Female
0.0
−0.5
−1.0
−1.5
−2.0
0
400
800
1200
1600
Age (Days)

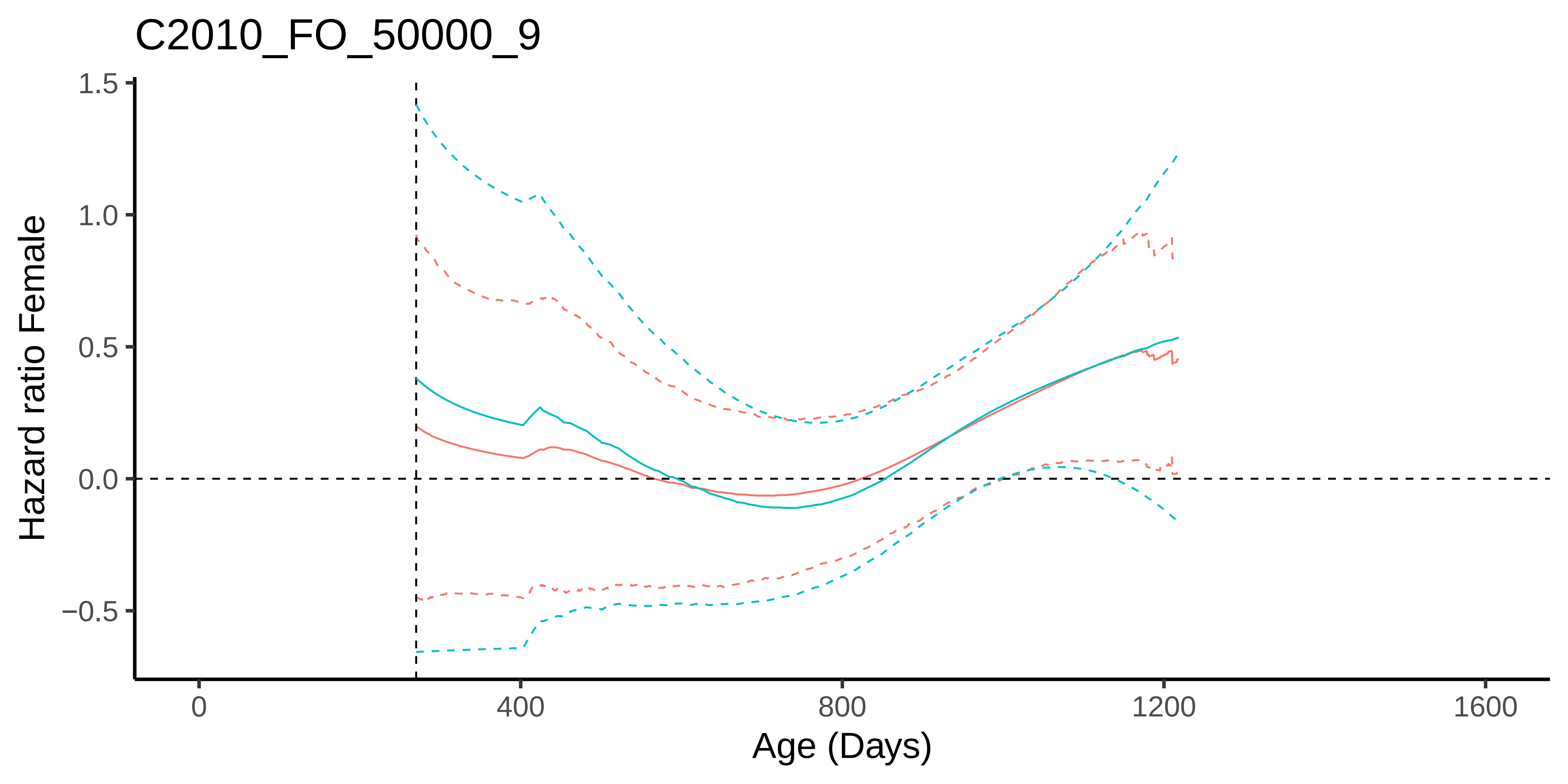

C2010_FO_50000_9
Hazard ratio Female
1.5
1.0
0.5
0.0
−0.5
0
400
800
1200
1600
Age (Days)

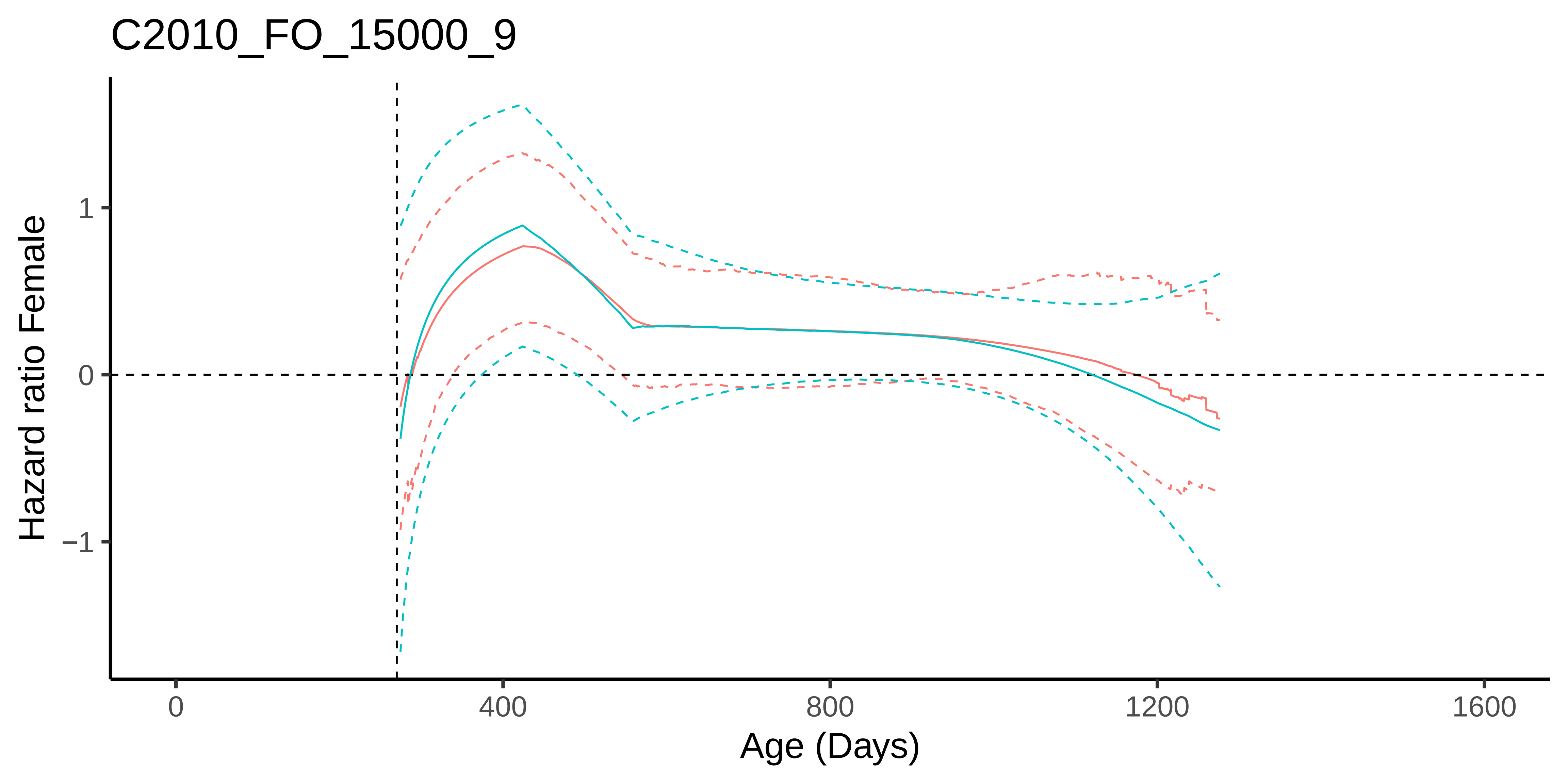
C2010_FO_15000_9
Hazard ratio Female
1
0
−1
0
400
800
1200
1600
Age (Days)

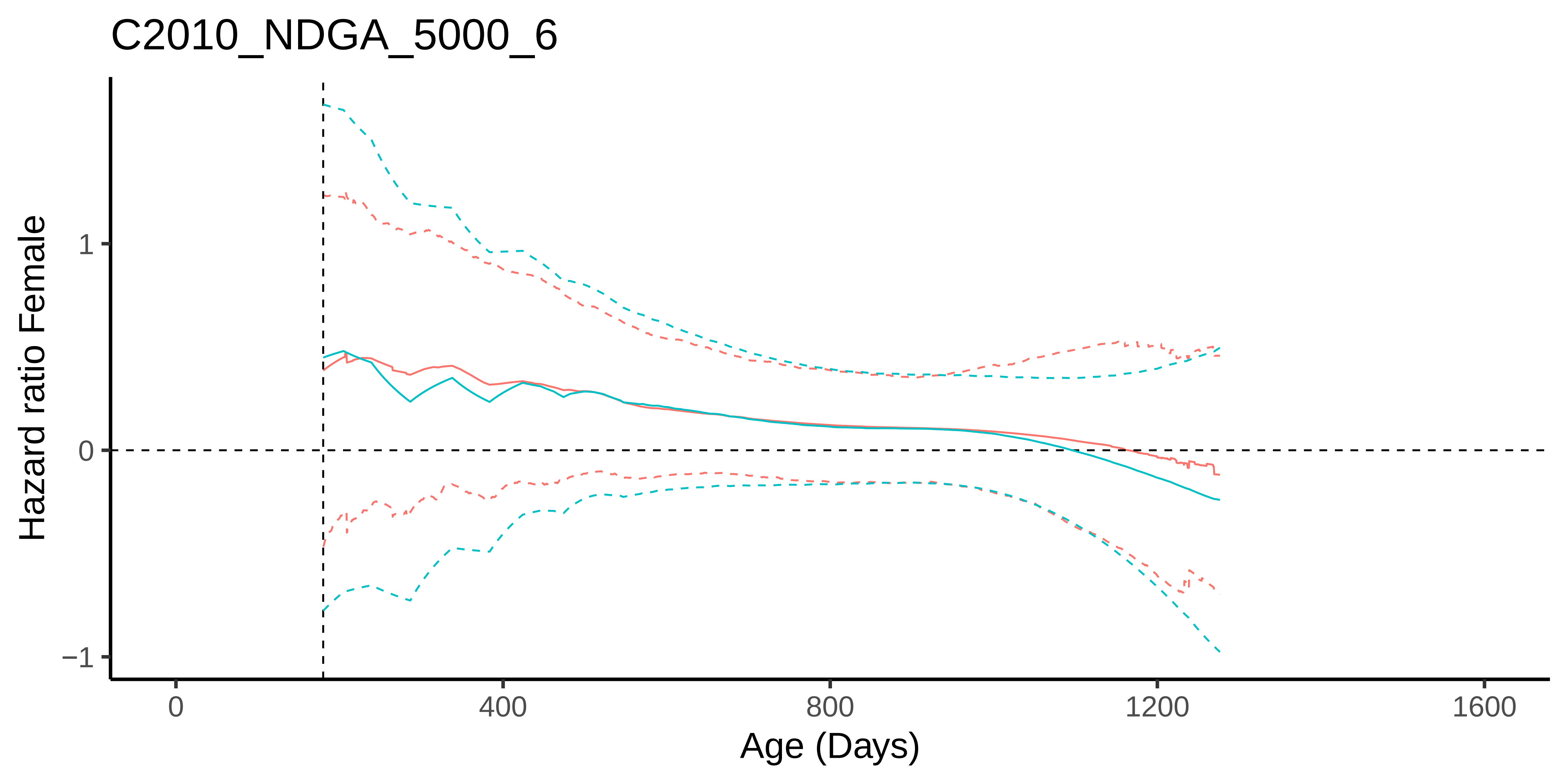
C2010_NDGA_5000_6
Hazard ratio Female
1
0
−1
0
400
800
1200
1600
Age (Days)

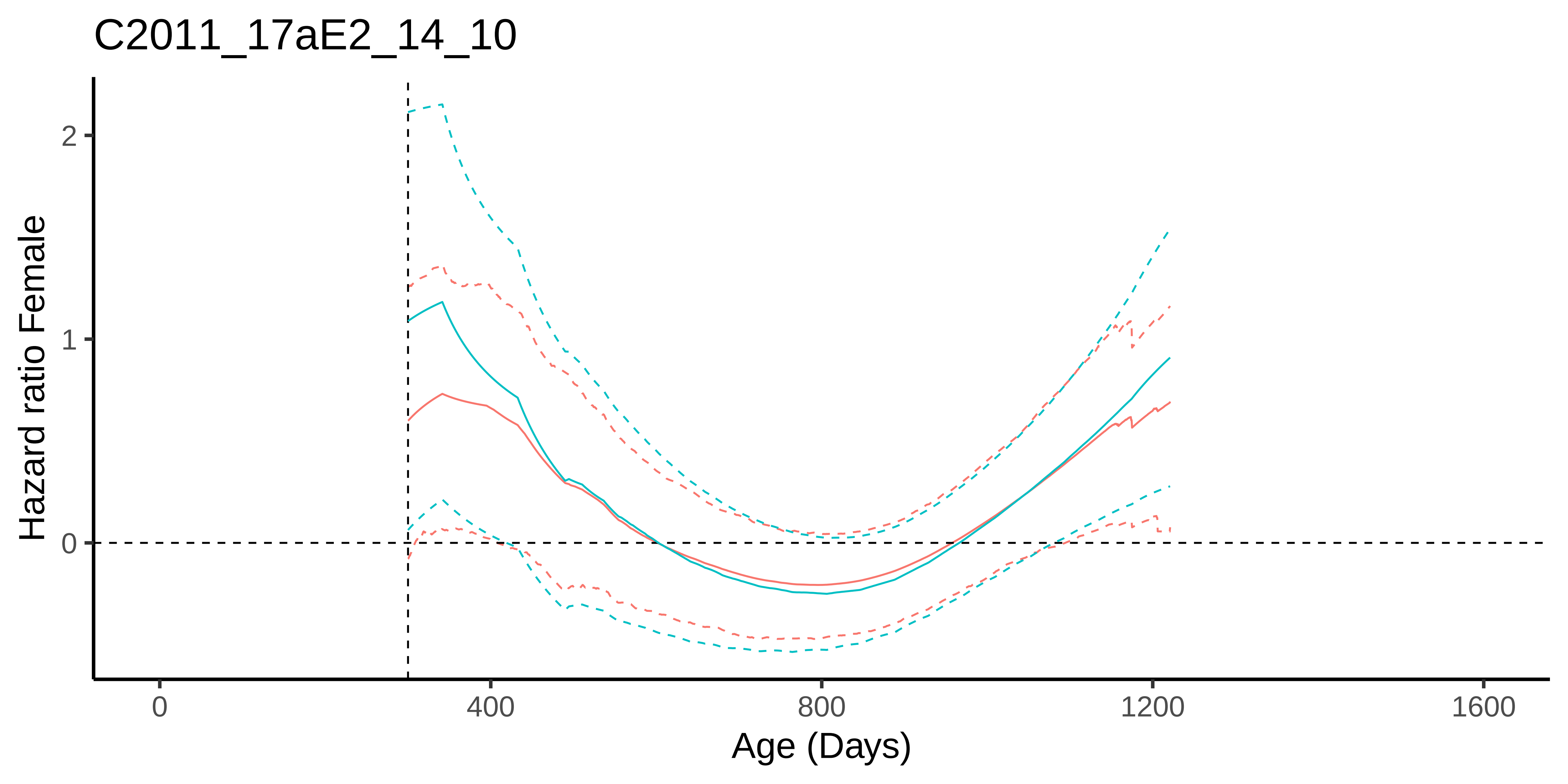

C2011_17aE2_14_10
Hazard ratio Female
2
1
0
0
400
800
1200
1600
Age (Days)

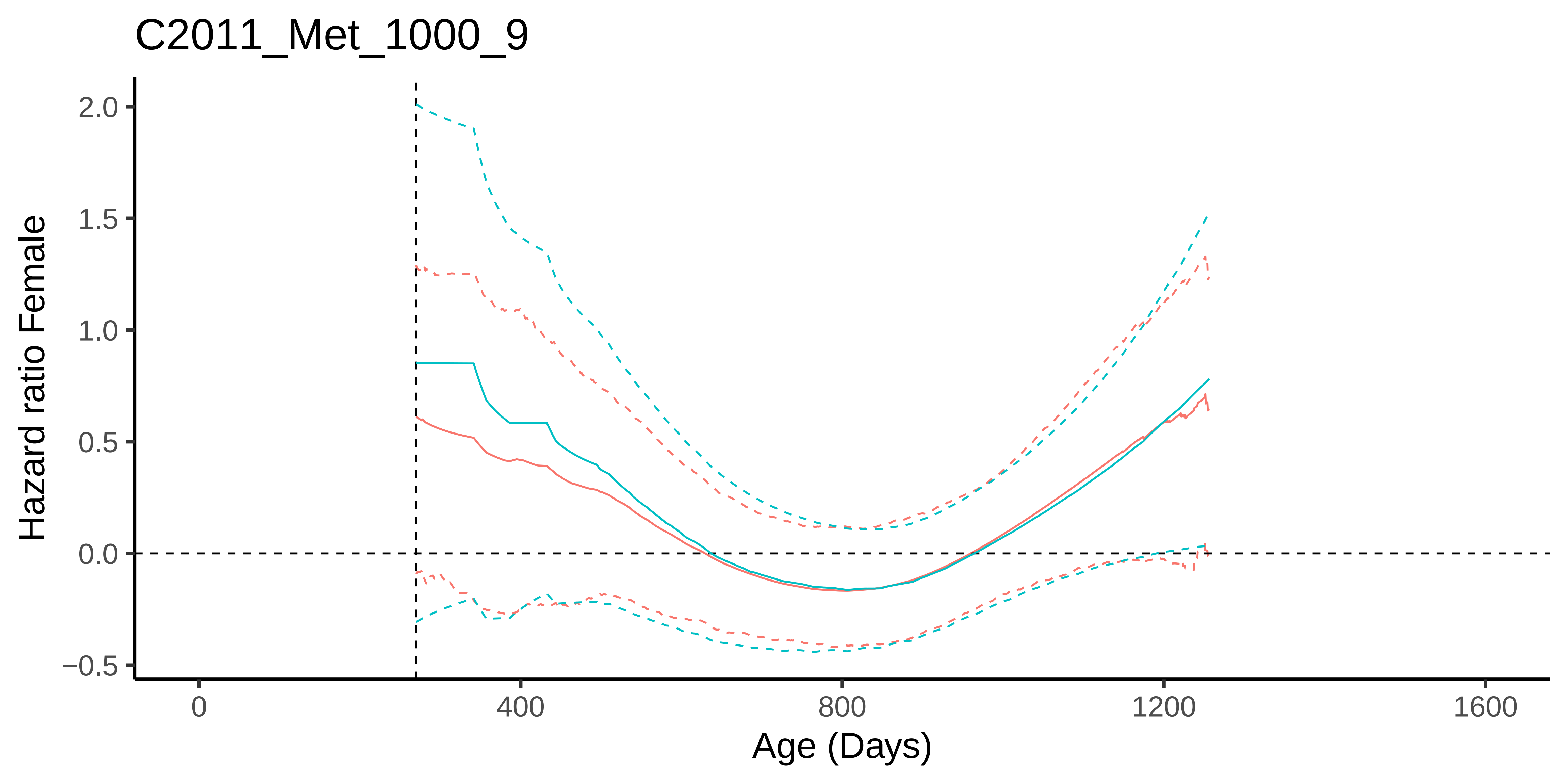

C2011_Met_1000_9
Hazard ratio Female
2.0
1.5
1.0
0.5
0.0
−0.5
0
400
800
1200
1600
Age (Days)

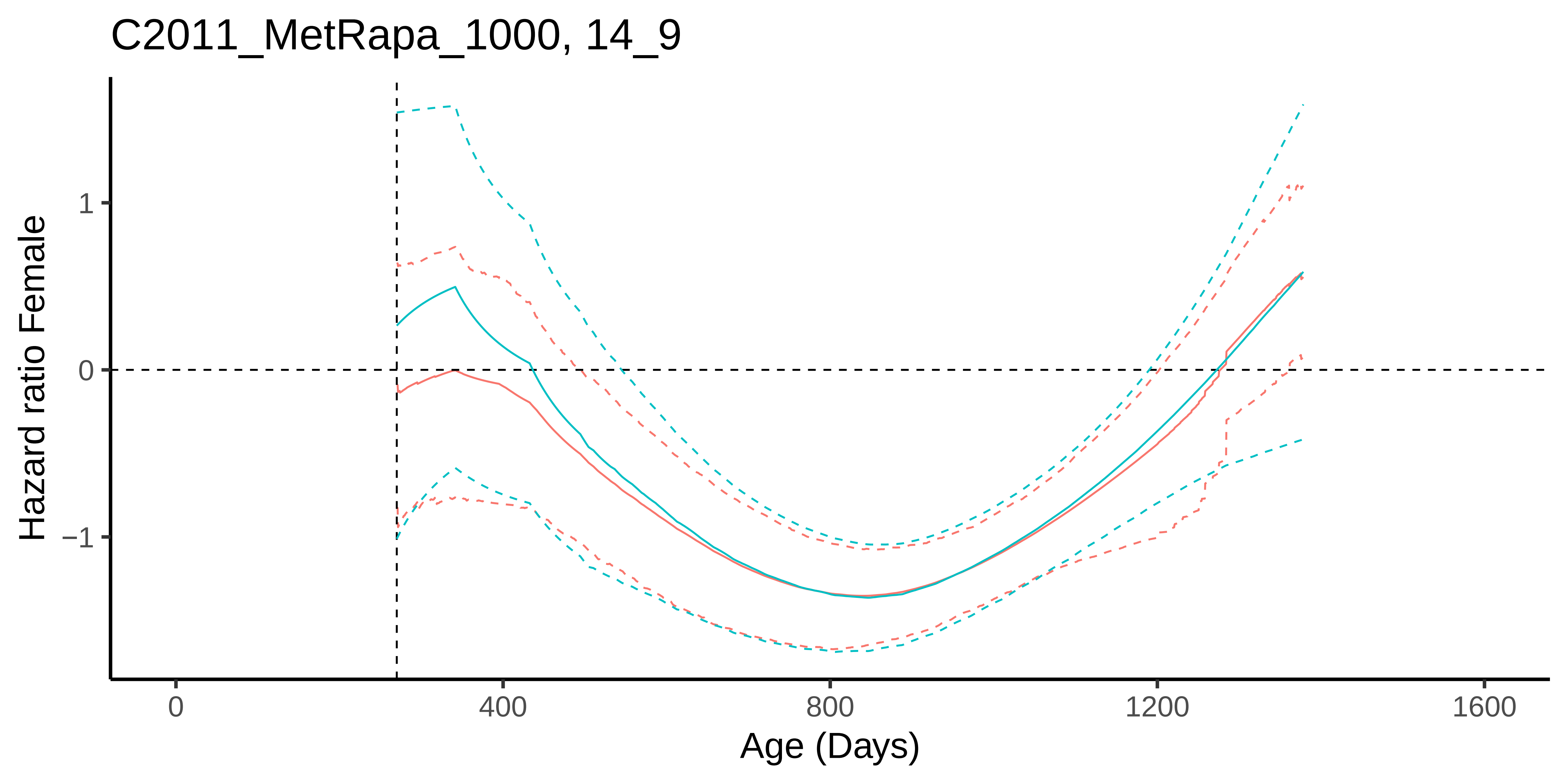
C2011_MetRapa_1000, 14_9
Hazard ratio Female
1
0
−1
0
400
800
1200
1600
Age (Days)

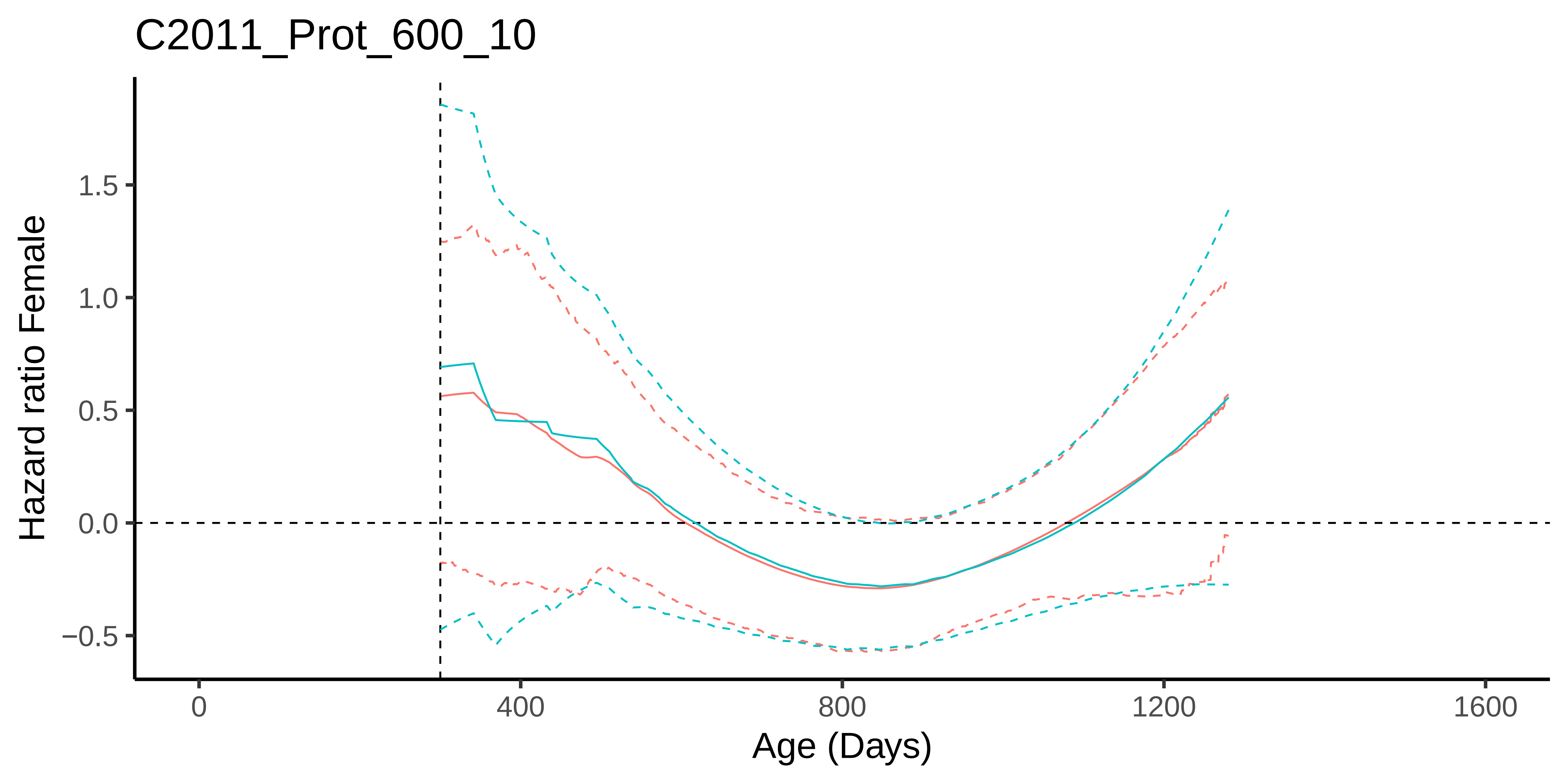
C2011_Prot_600_10
Hazard ratio Female
1.5
1.0
0.5
0.0
−0.5
0
400
800
1200
1600
Age (Days)

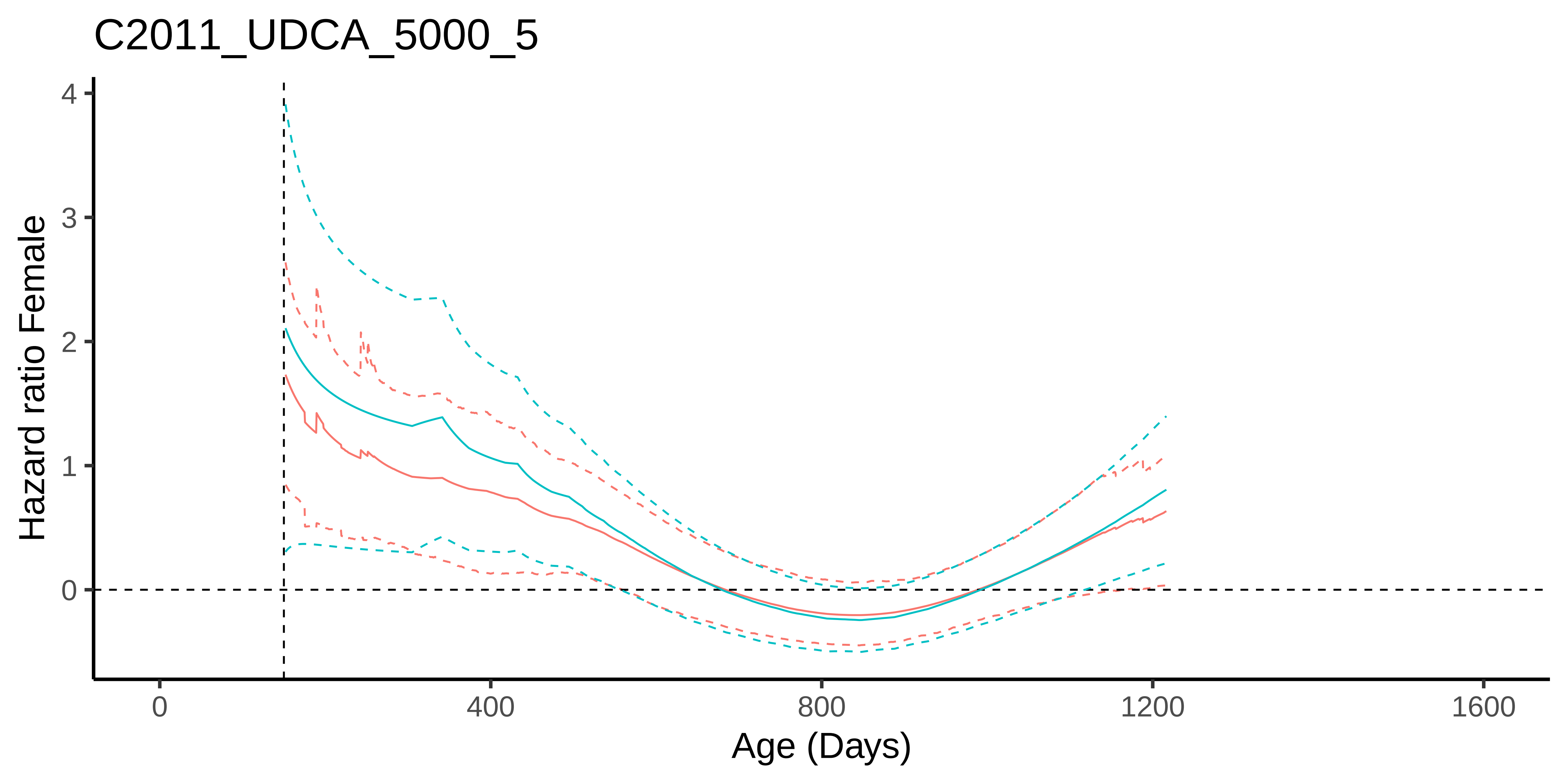

C2011_UDCA_5000_5
Hazard ratio Female
Age (Days)
0
1
2
3
4
0
400
800
1200
1600

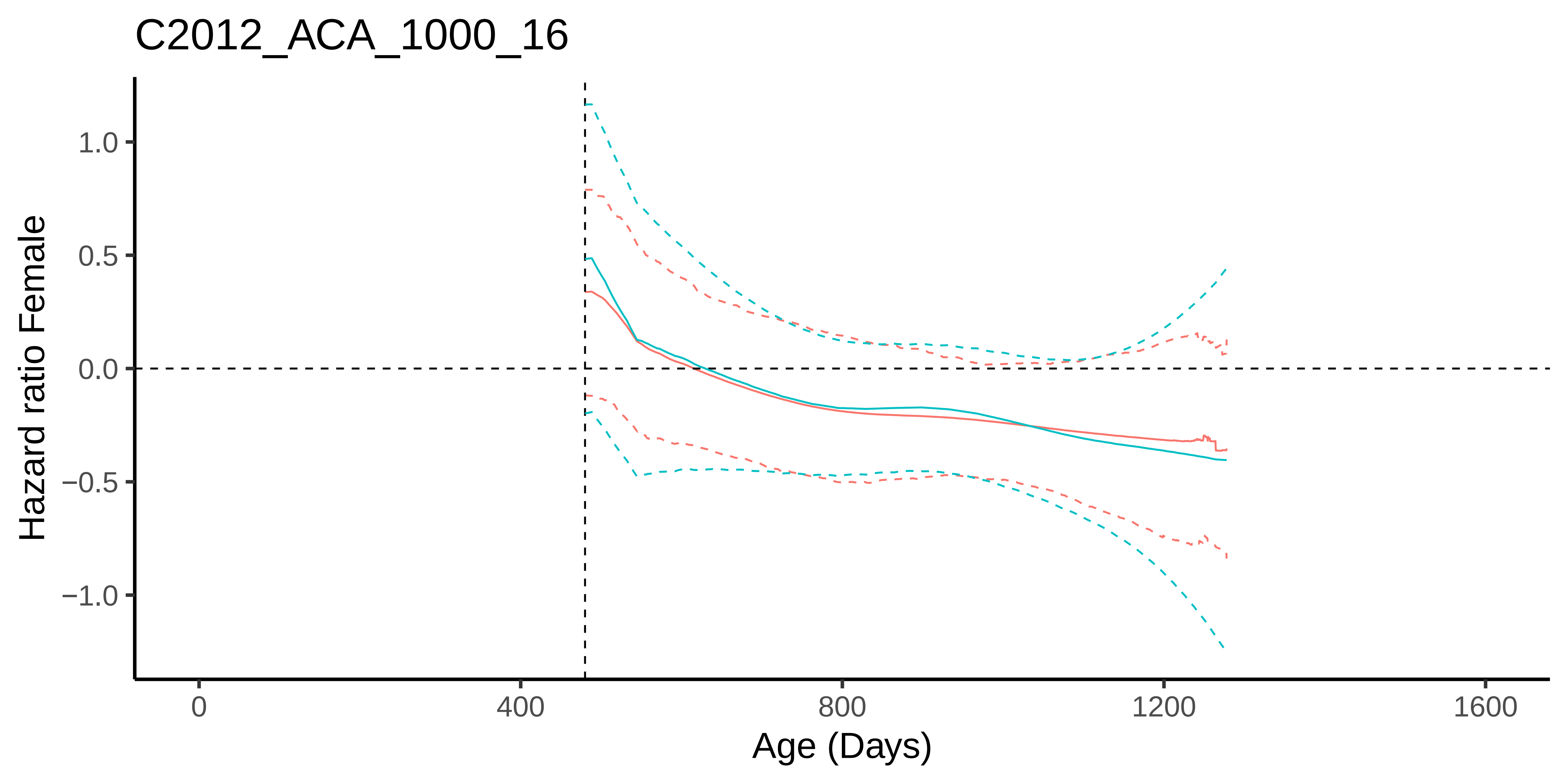

C2012_ACA_1000_16
Hazard ratio Female
1.0
0.5
0.0
−0.5
−1.0
0
400
800
1200
1600
Age (Days)

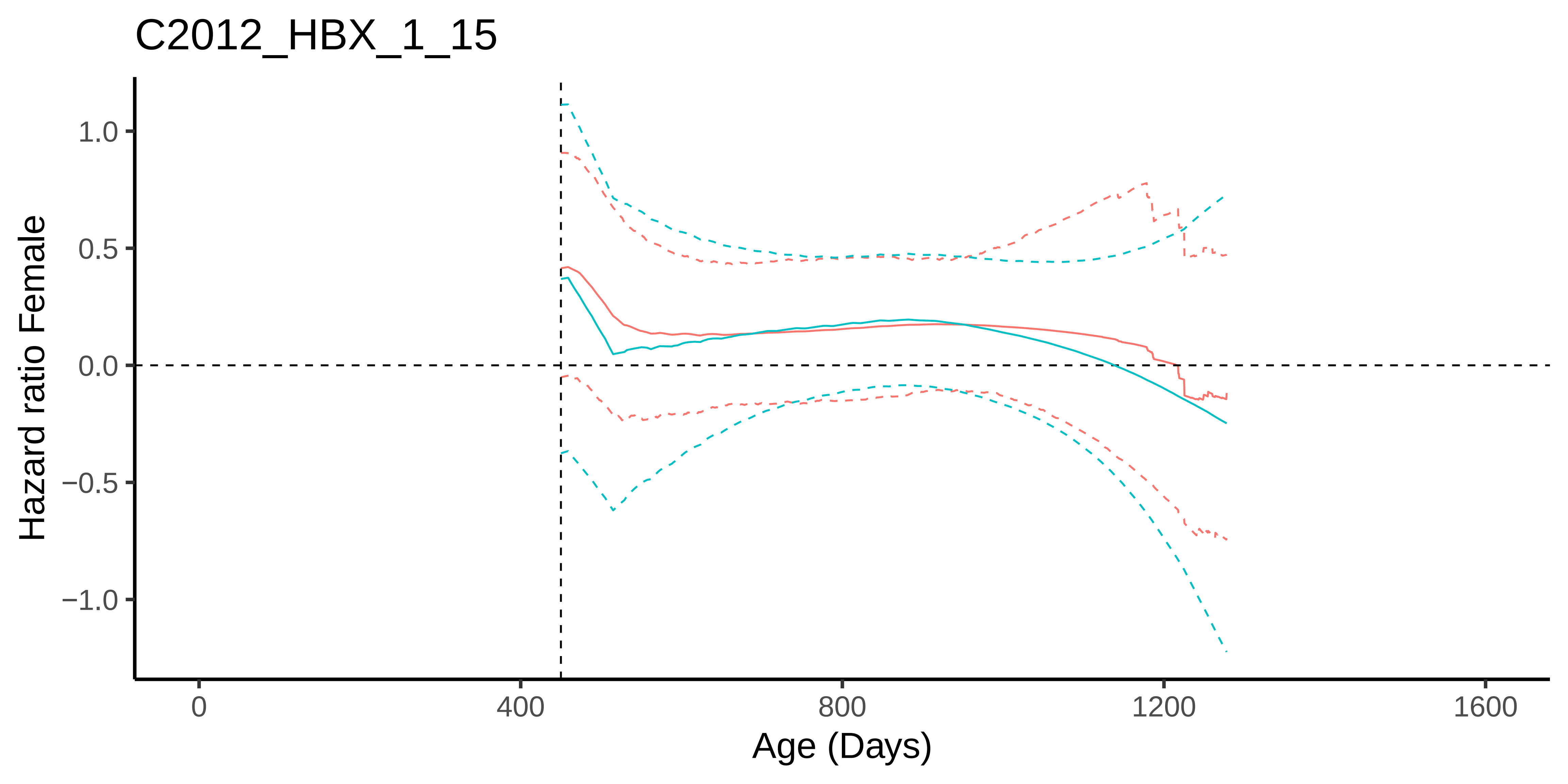
C2012_HBX_1_15
Hazard ratio Female
1.0
0.5
0.0
−0.5
−1.0
0
400
800
1200
1600
Age (Days)

### C2012_INT−767_180_10

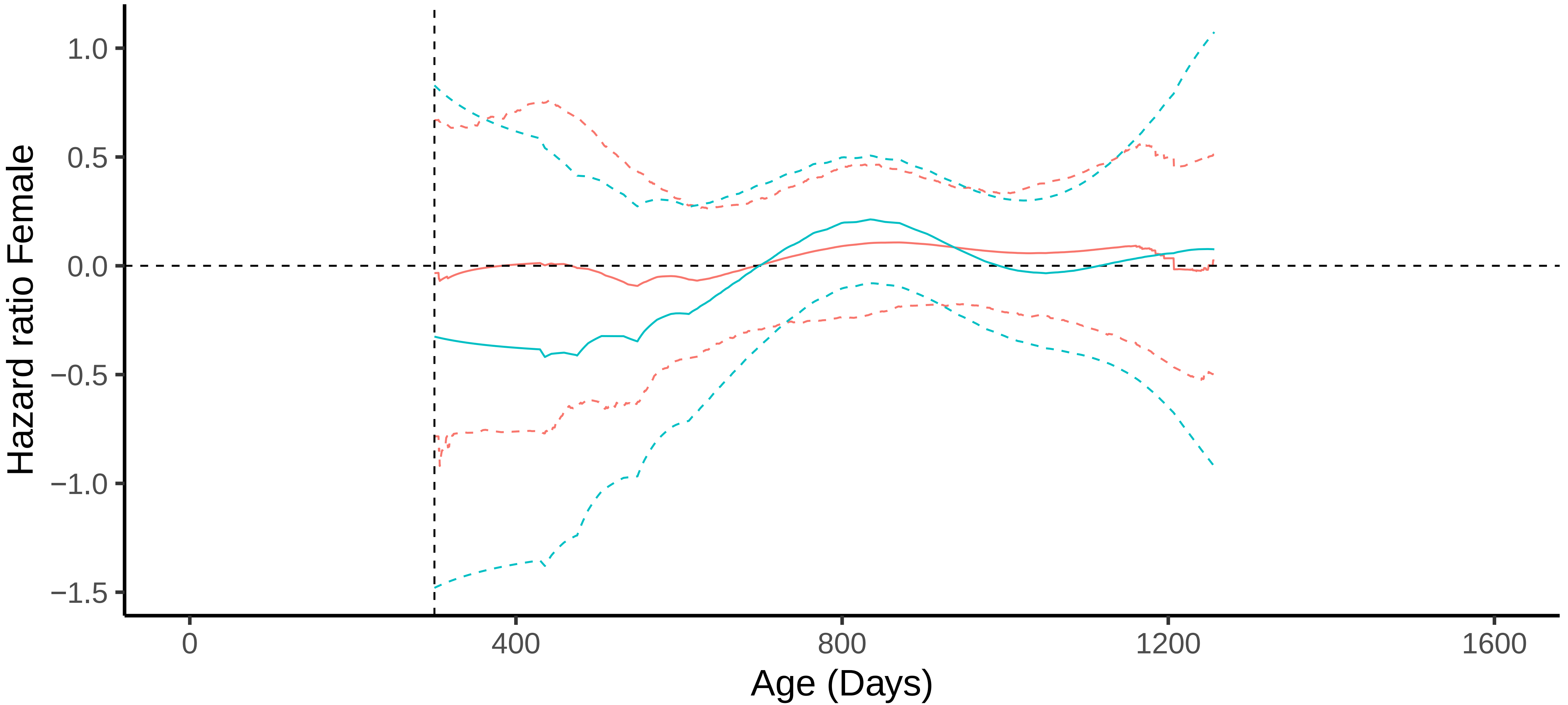

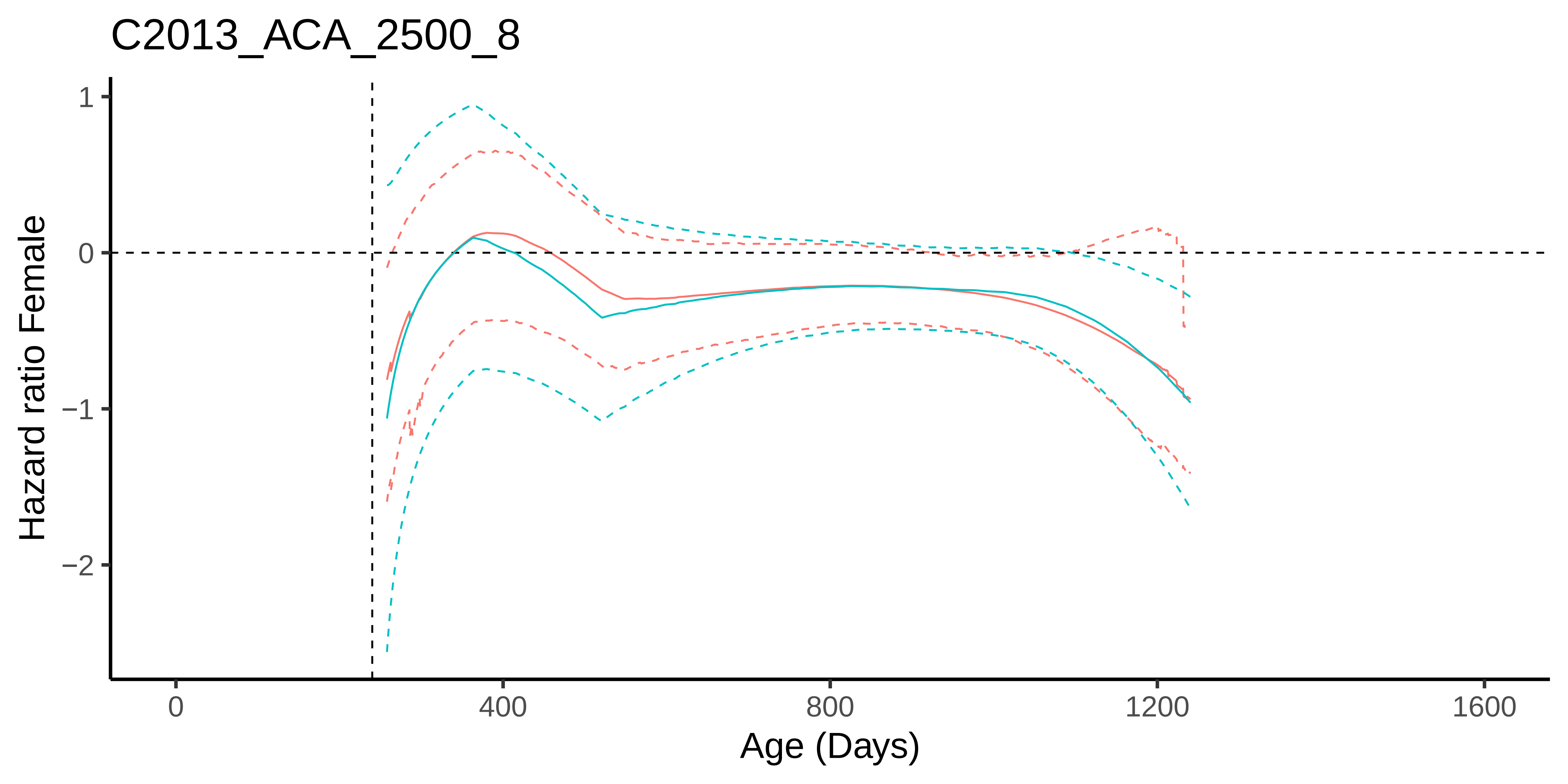
C2013_ACA_2500_8
Hazard ratio Female
1
0
−1
−2
0
400
800
1200
1600
Age (Days)

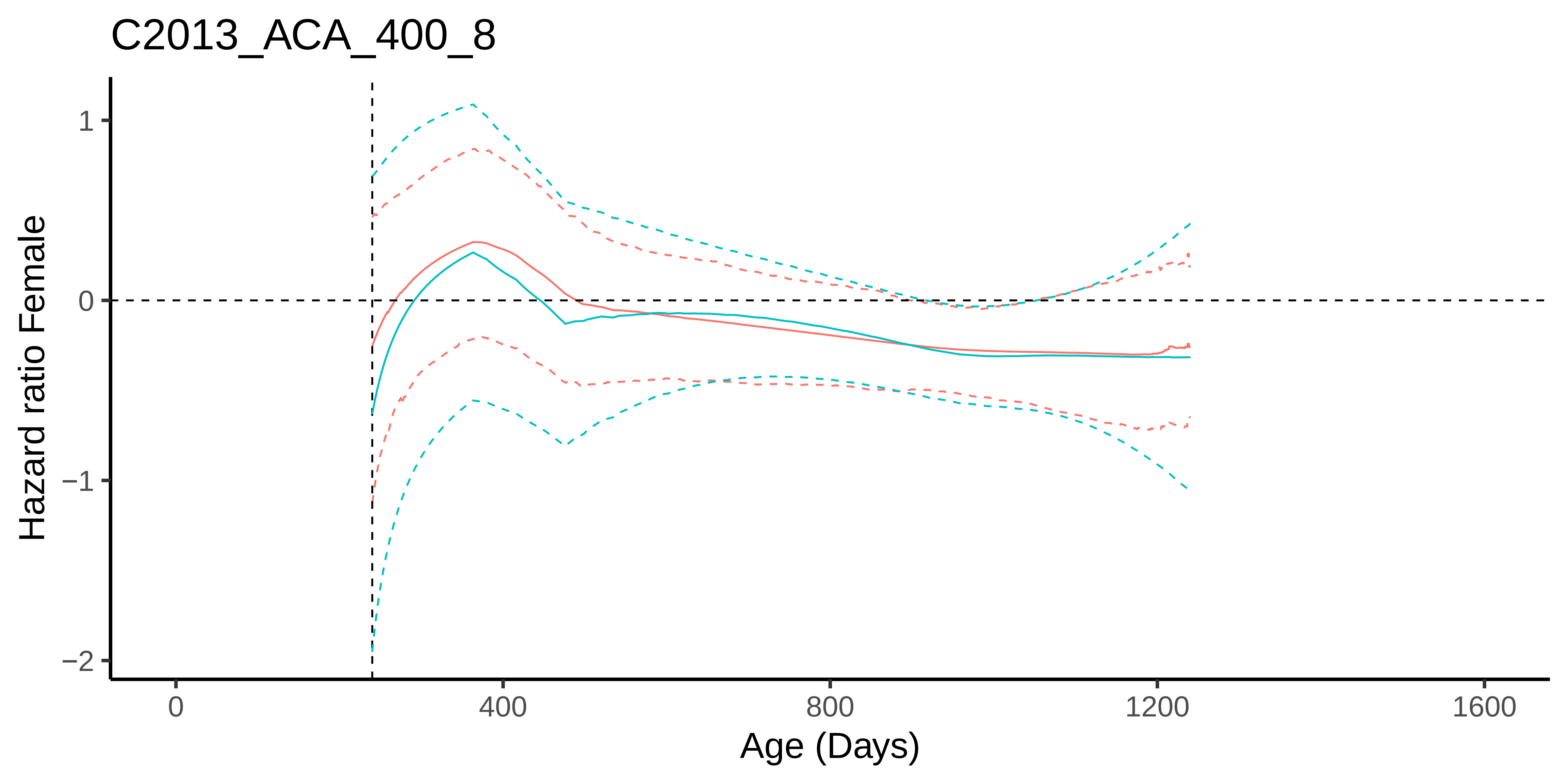
C2013_ACA_400_8
Hazard ratio Female
1
0
−1
−2
0
400
800
1200
1600
Age (Days)

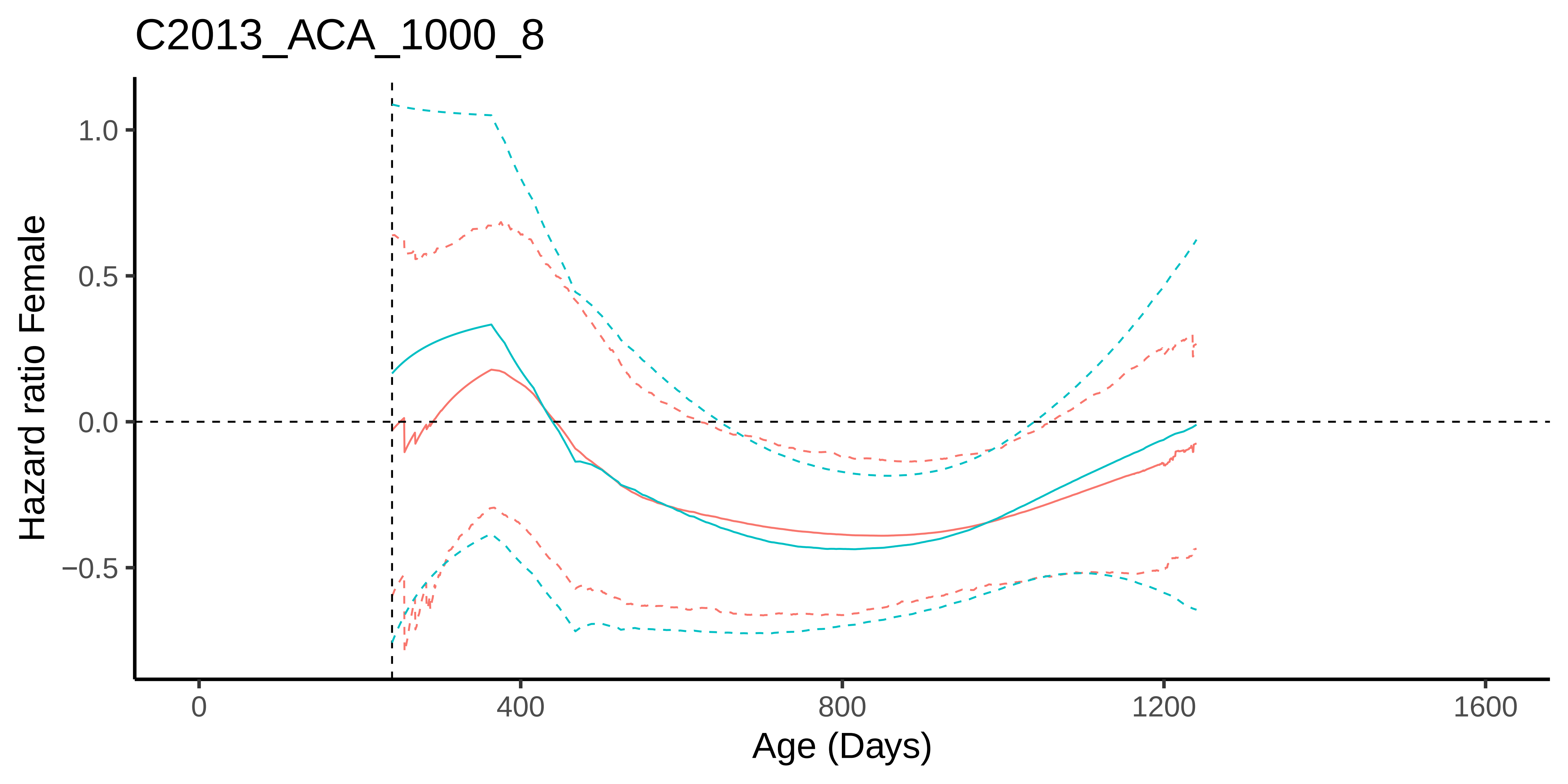

C2013_ACA_1000_8
Hazard ratio Female
1.0
0.5
0.0
−0.5
0
400
800
1200
1600
Age (Days)

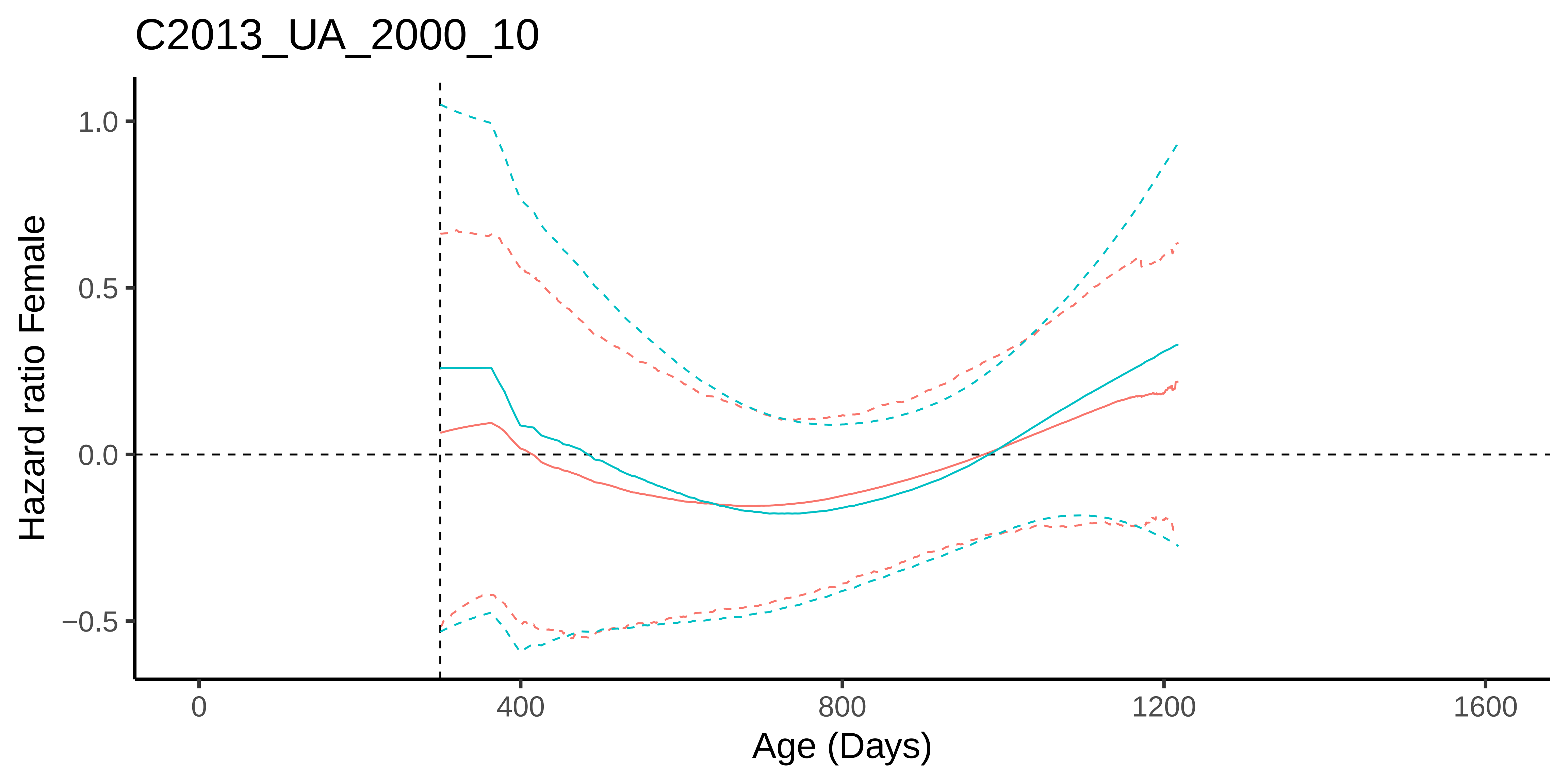
C2013_UA_2000_10
Hazard ratio Female
1.0
0.5
0.0
−0.5
0
400
800
1200
1600
Age (Days)

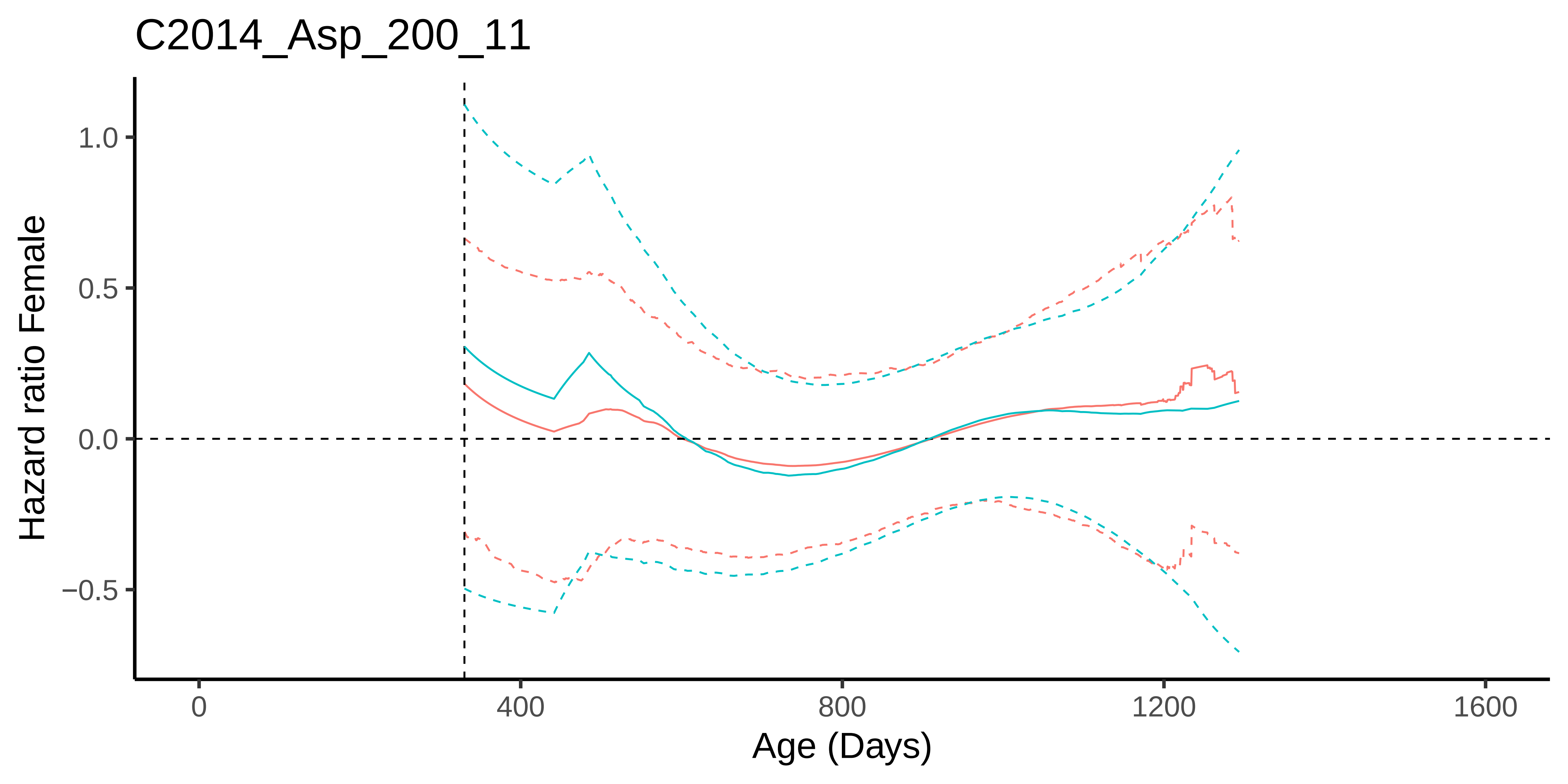

C2014_Asp_200_11
Hazard ratio Female
1.0
0.5
0.0
−0.5
0
400
800
1200
1600
Age (Days)

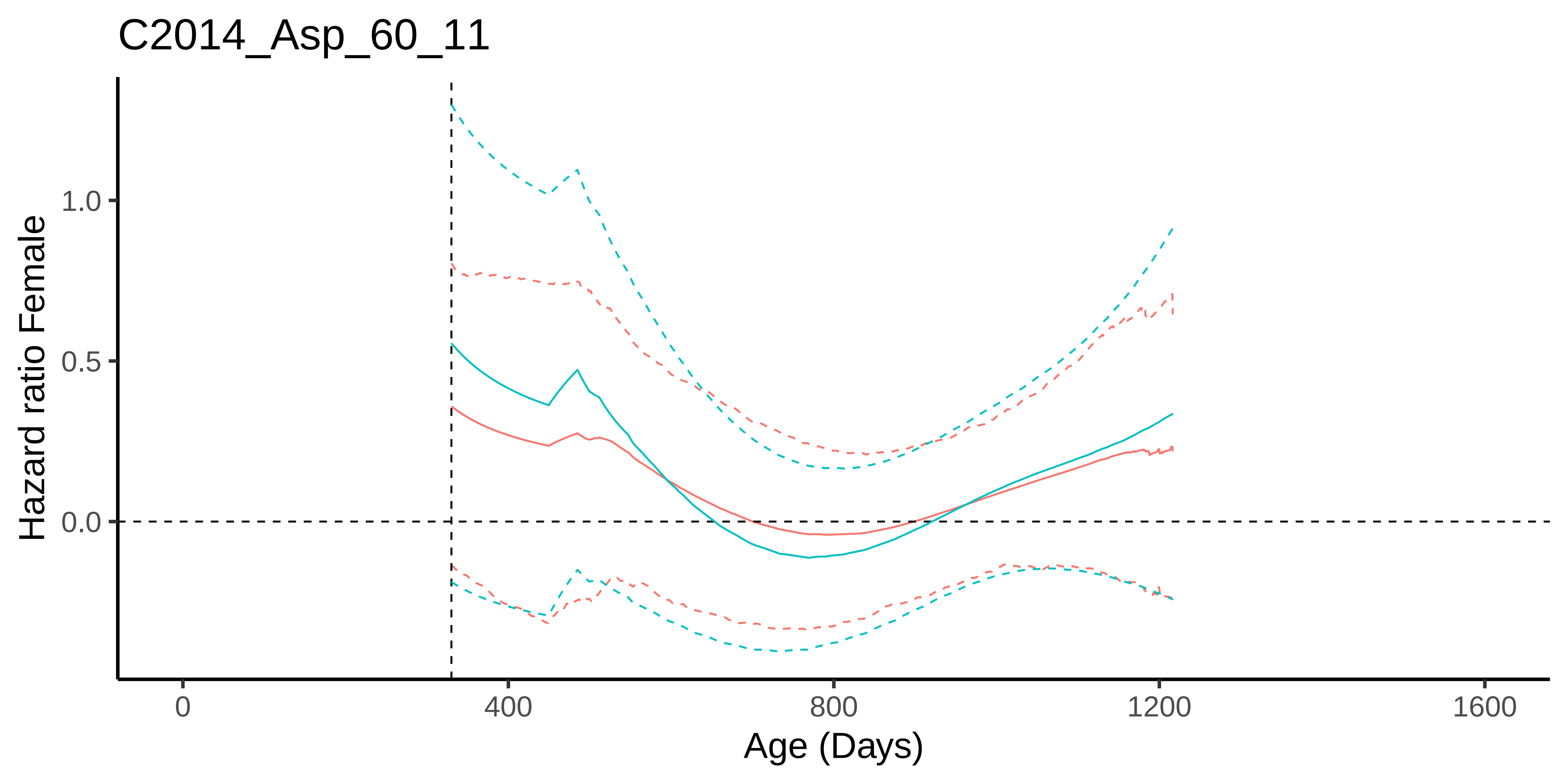

C2014_Asp_60_11
Hazard ratio Female
1.0
0.5
0.0
0
400
800
1200
1600
Age (Days)

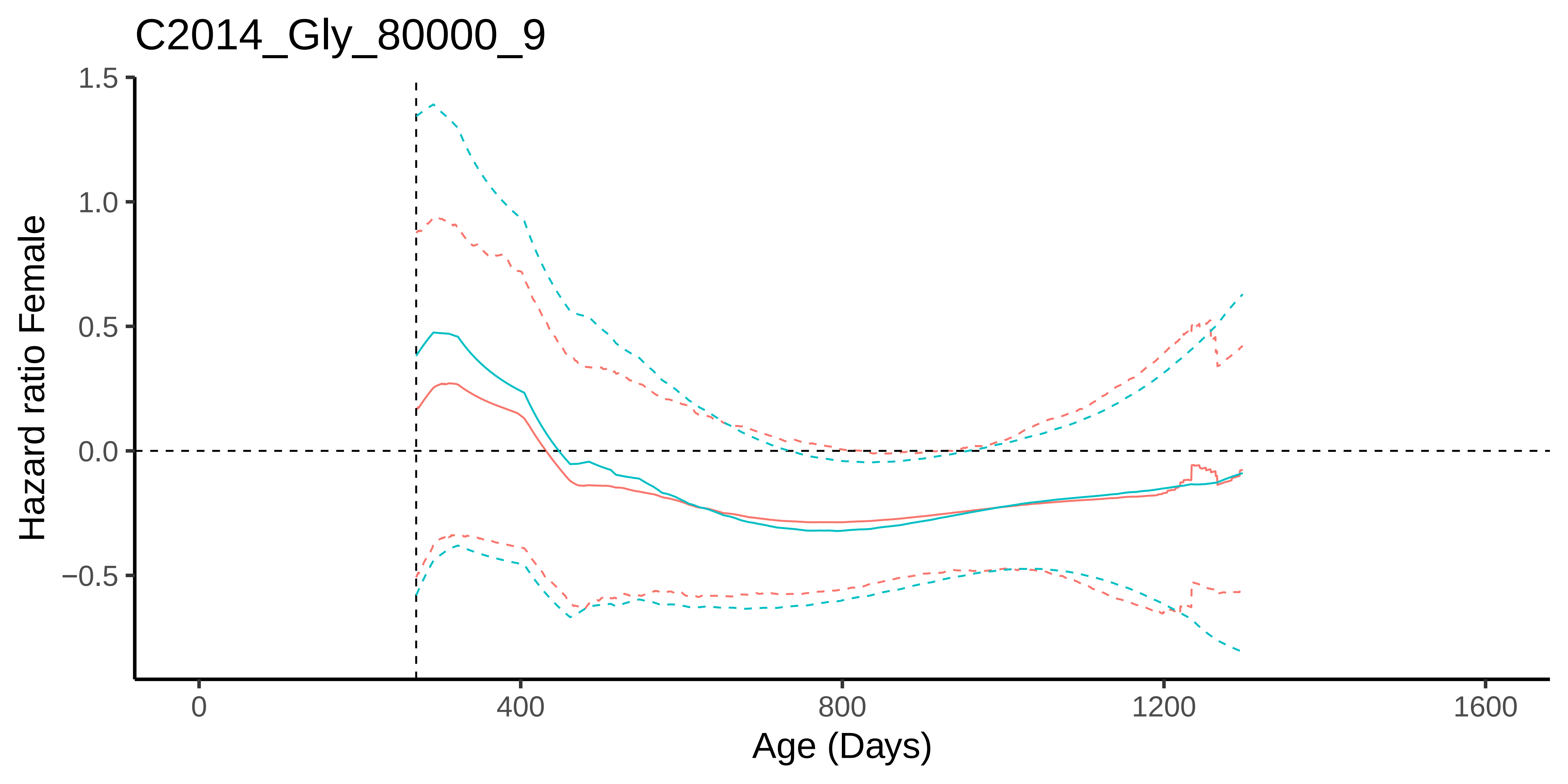
C2014_Gly_80000_9
Hazard ratio Female
1.5
1.0
0.5
0.0
−0.5
0
400
800
1200
1600
Age (Days)

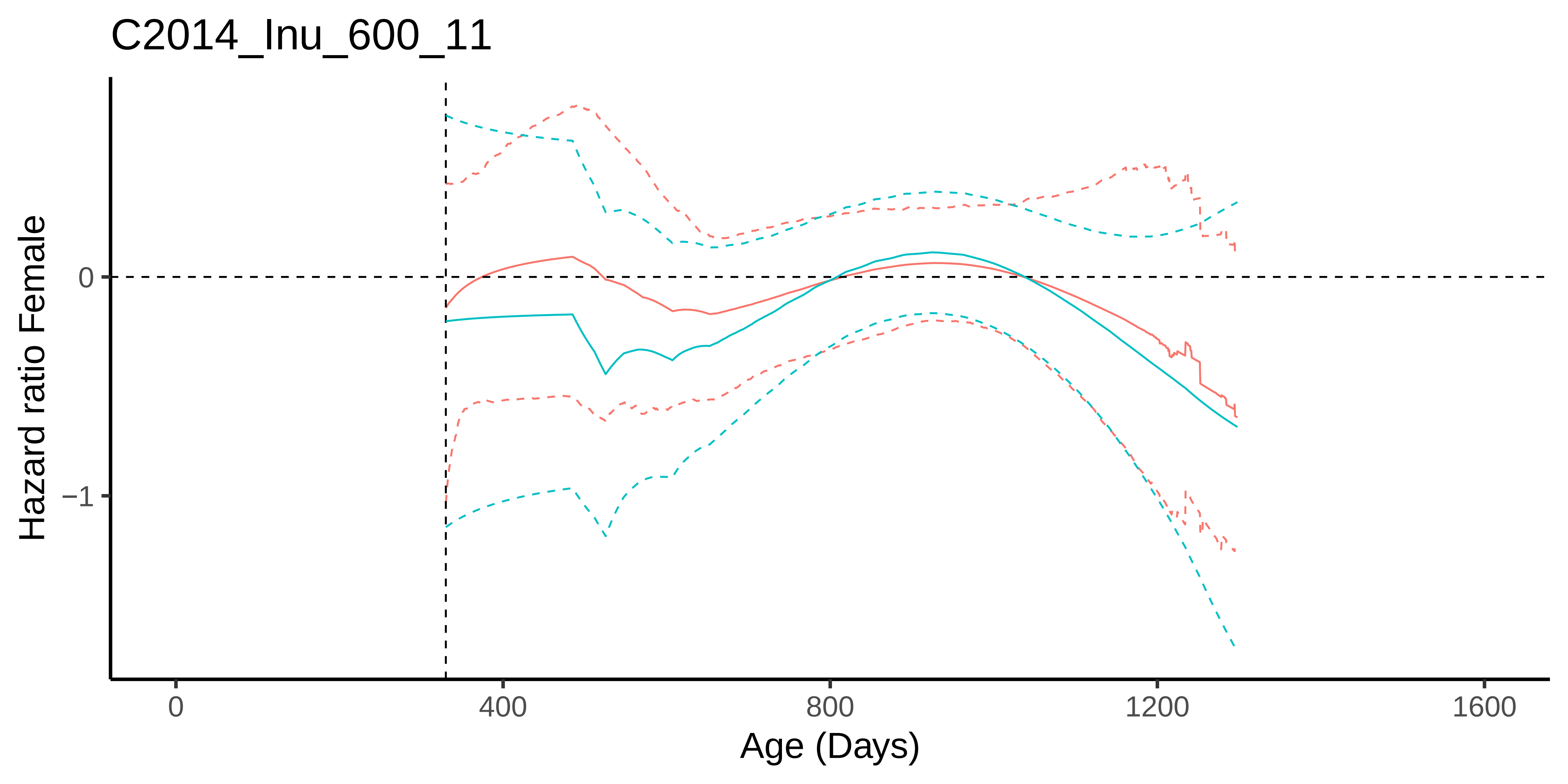
C2014_Inu_600_11
Hazard ratio Female
0
−1
0
400
800
1200
1600
Age (Days)

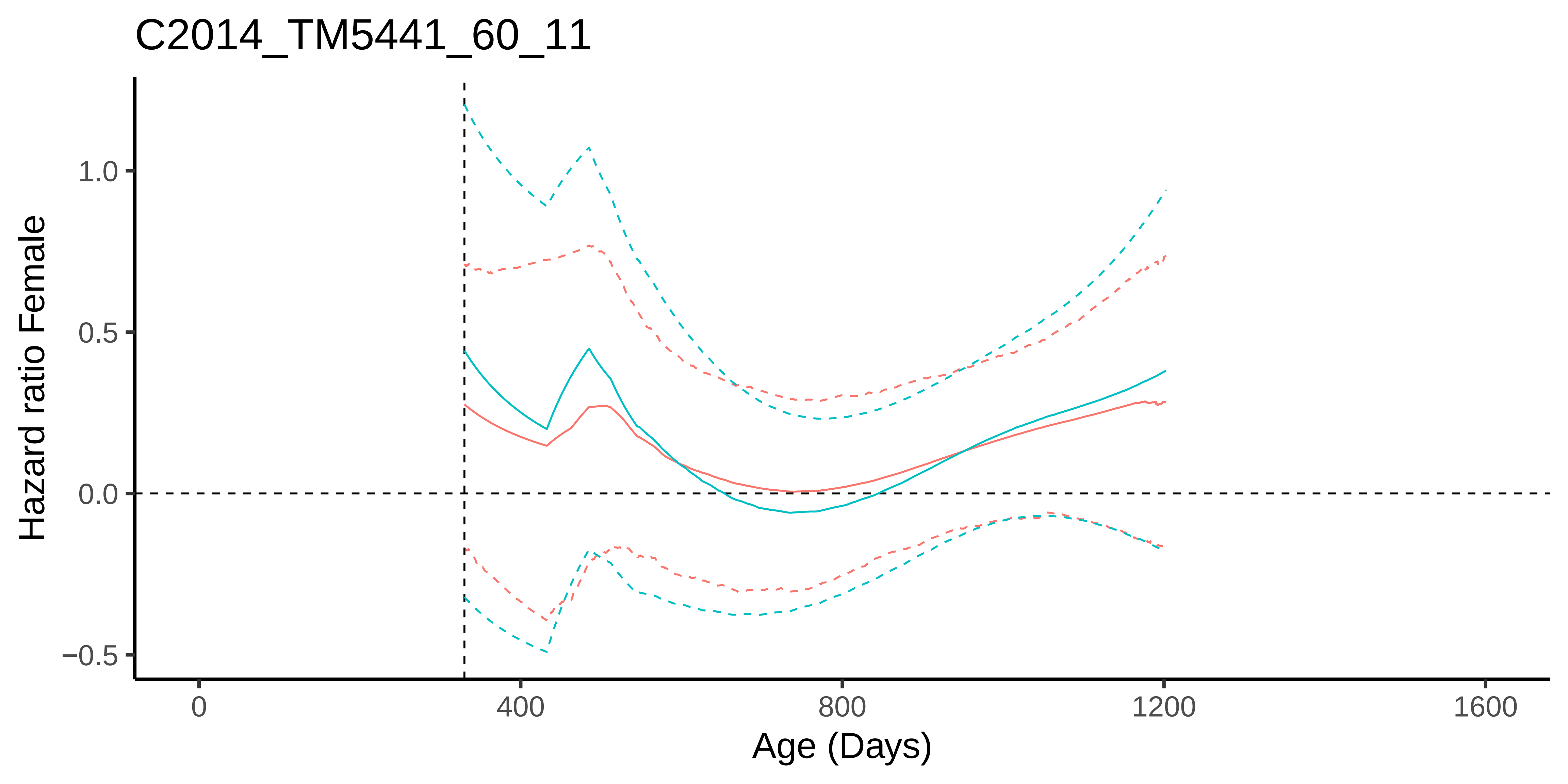

C2014_TM5441_60_11
Hazard ratio Female
1.0
0.5
0.0
−0.5
0
400
800
1200
1600
Age (Days)

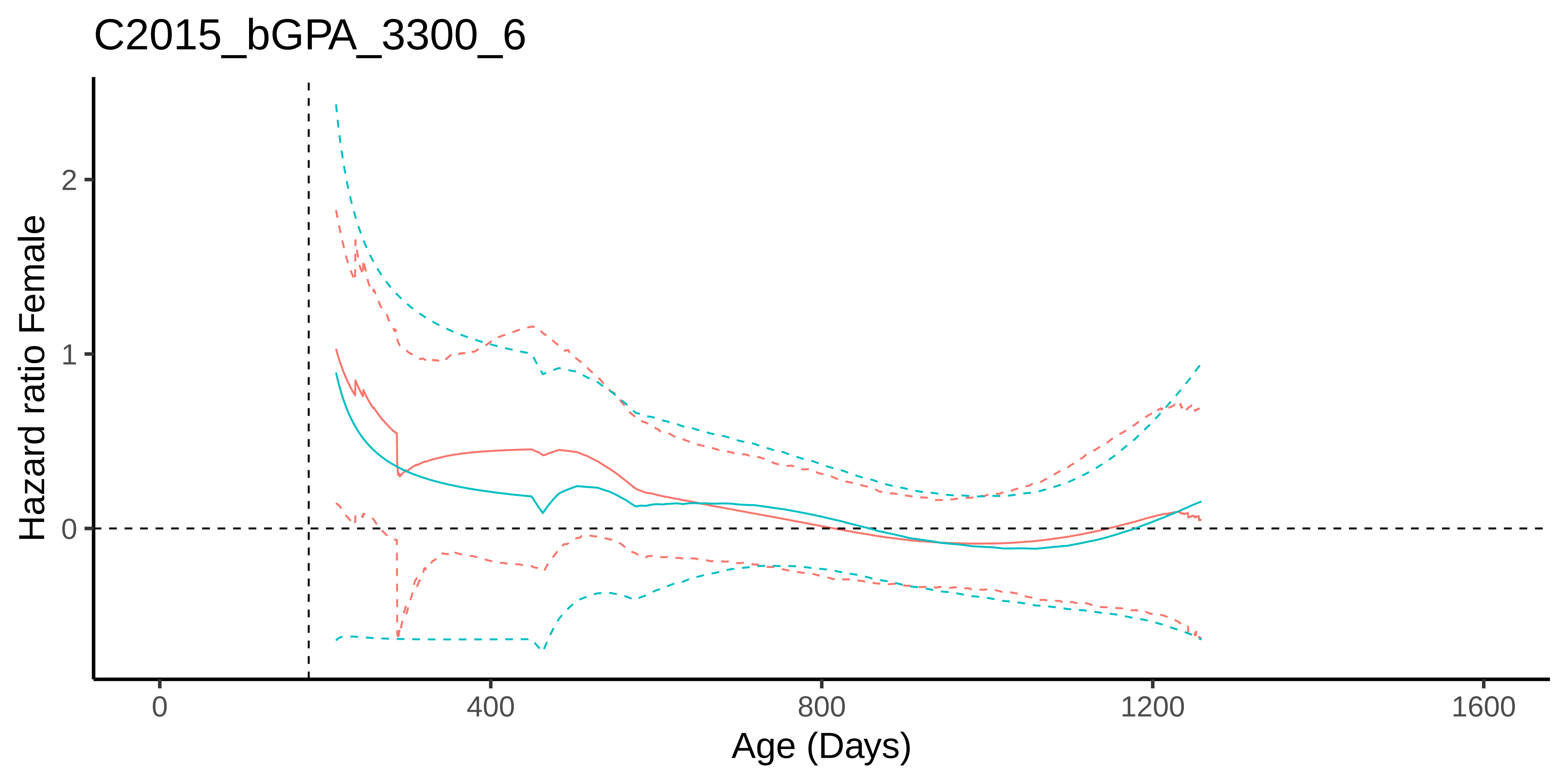

C2015_bGPA_3300_6
Hazard ratio Female
2
1
0
0
400
800
1200
1600
Age (Days)

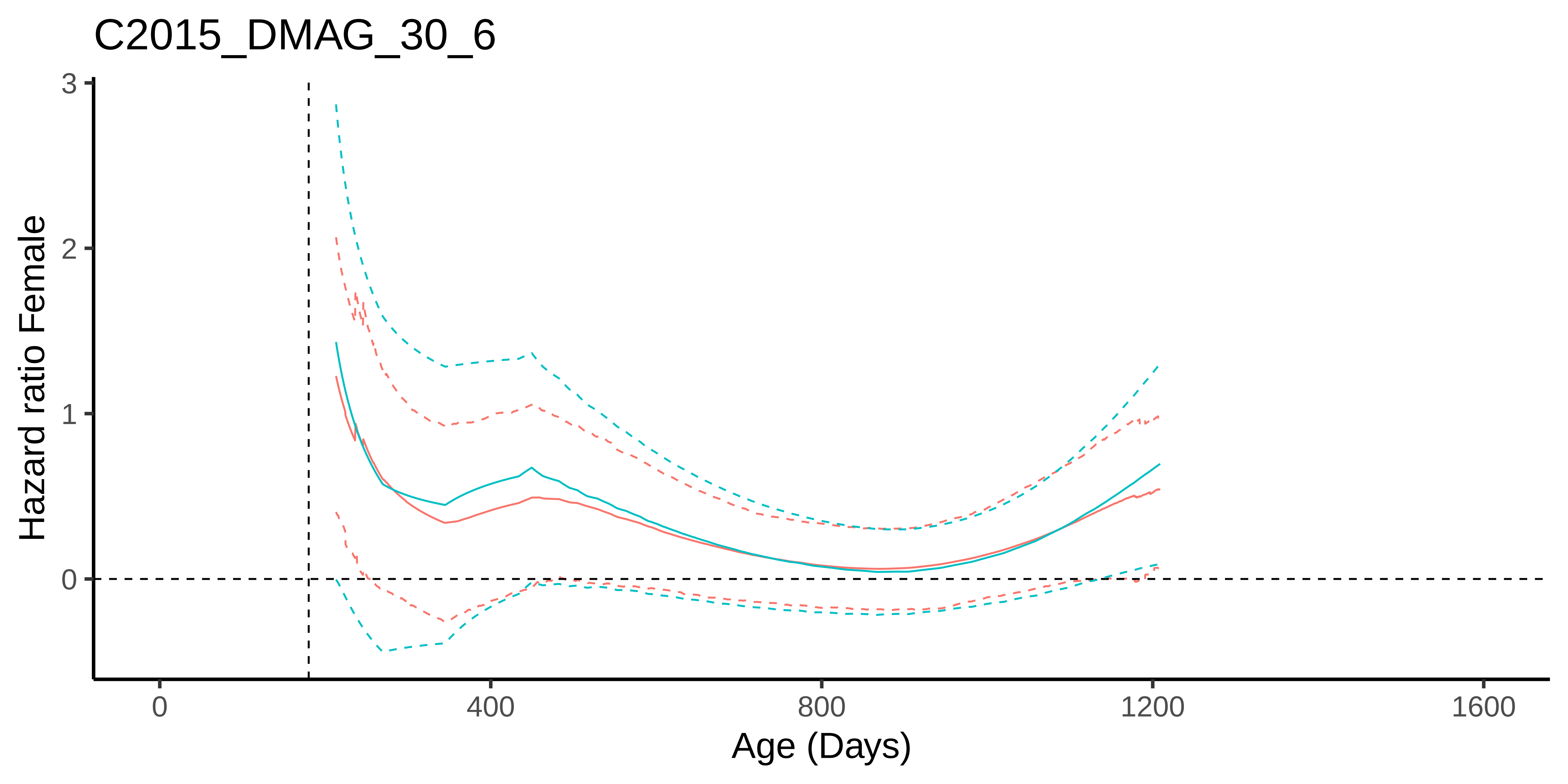

C2015_DMAG_30_6
Hazard ratio Female
3
2
1
0
0
400
800
1200
1600
Age (Days)

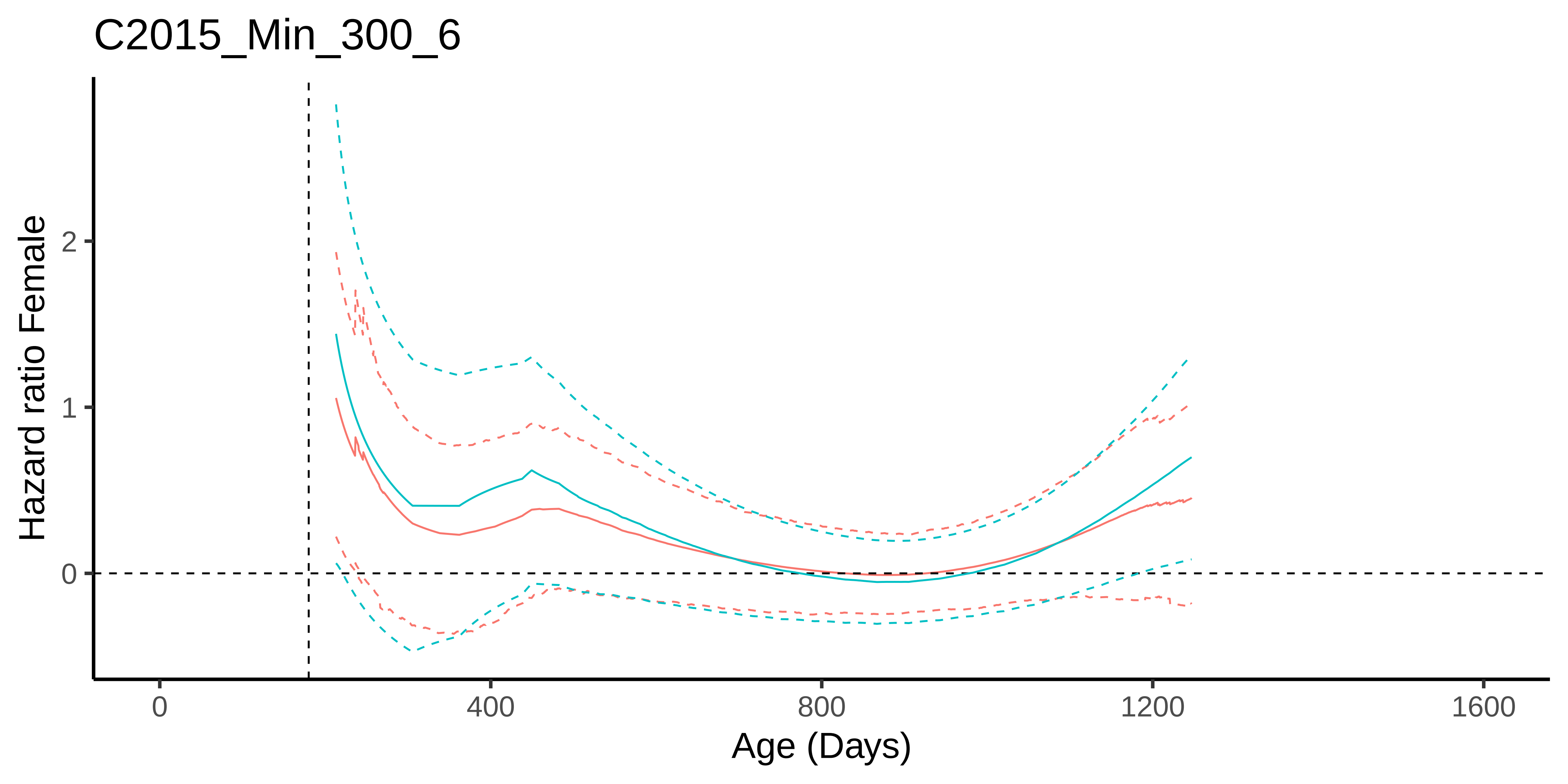
C2015_Min_300_6
Hazard ratio Female
2
1
0
0
400
800
1200
1600
Age (Days)

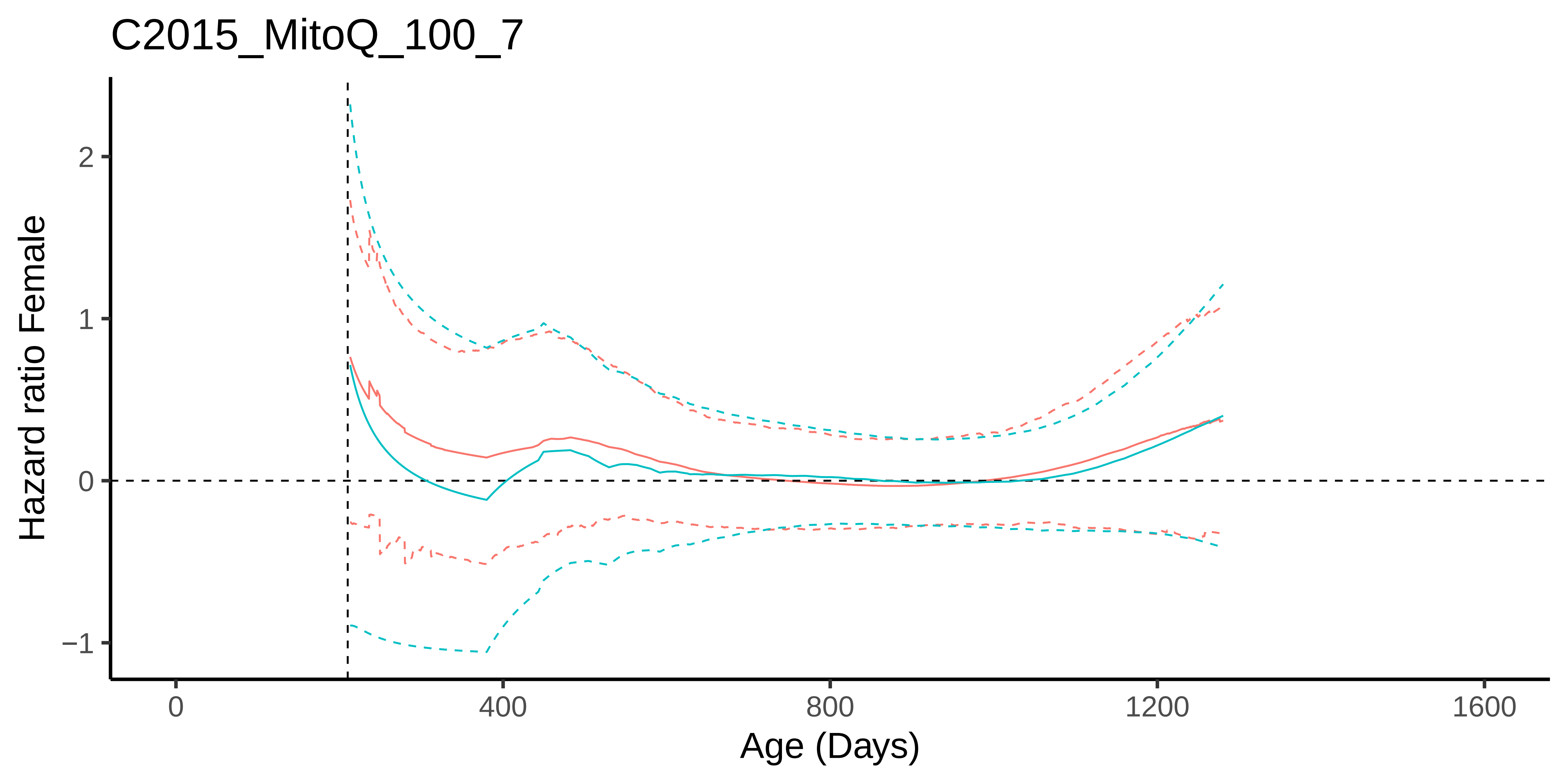

C2015_MitoQ_100_7
Hazard ratio Female
2
1
0
−1
0
400
800
1200
1600
Age (Days)

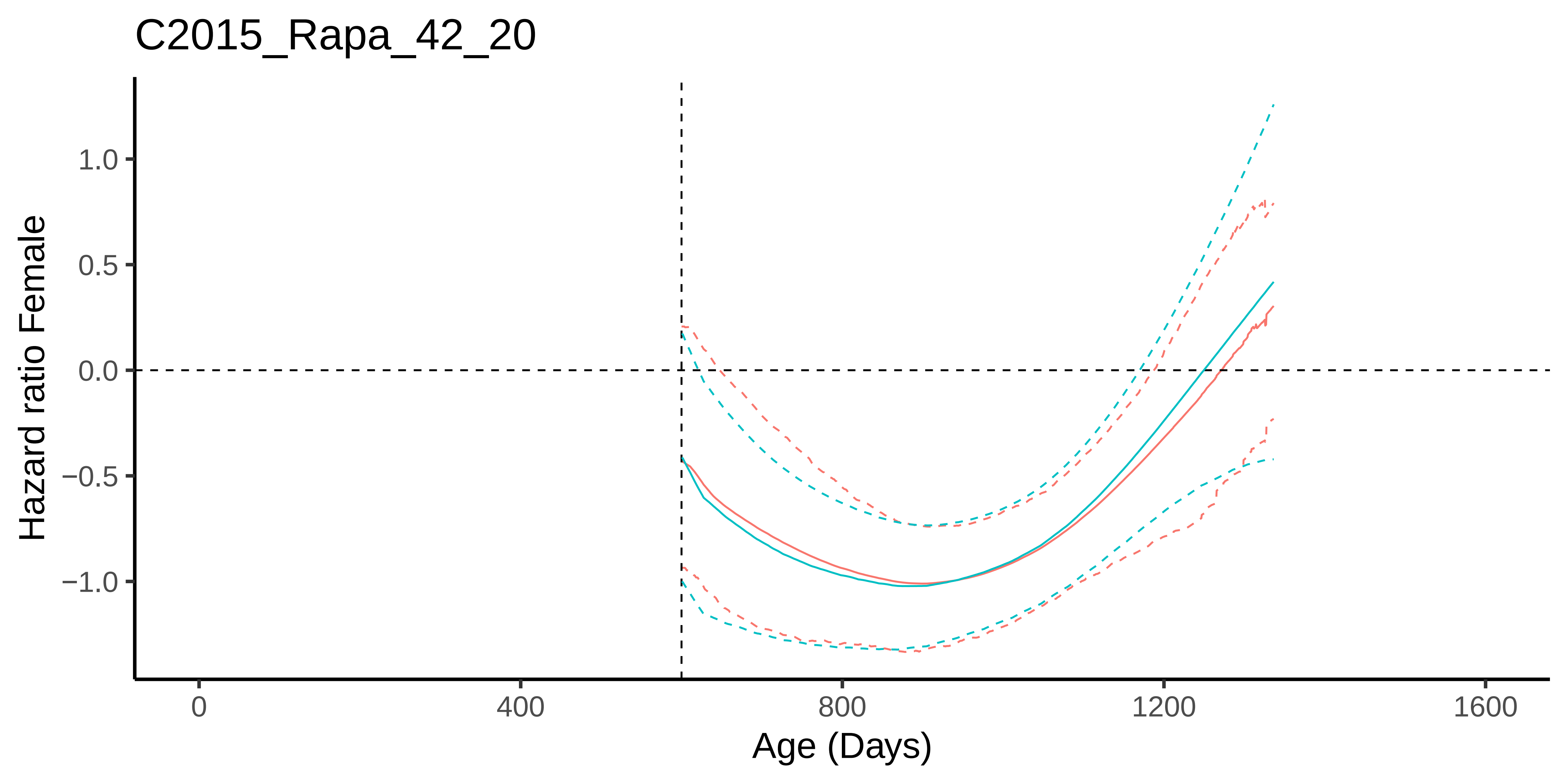

C2015_Rapa_42_20
Hazard ratio Female
1.0
0.5
0.0
−0.5
−1.0
0
400
800
1200
1600
Age (Days)

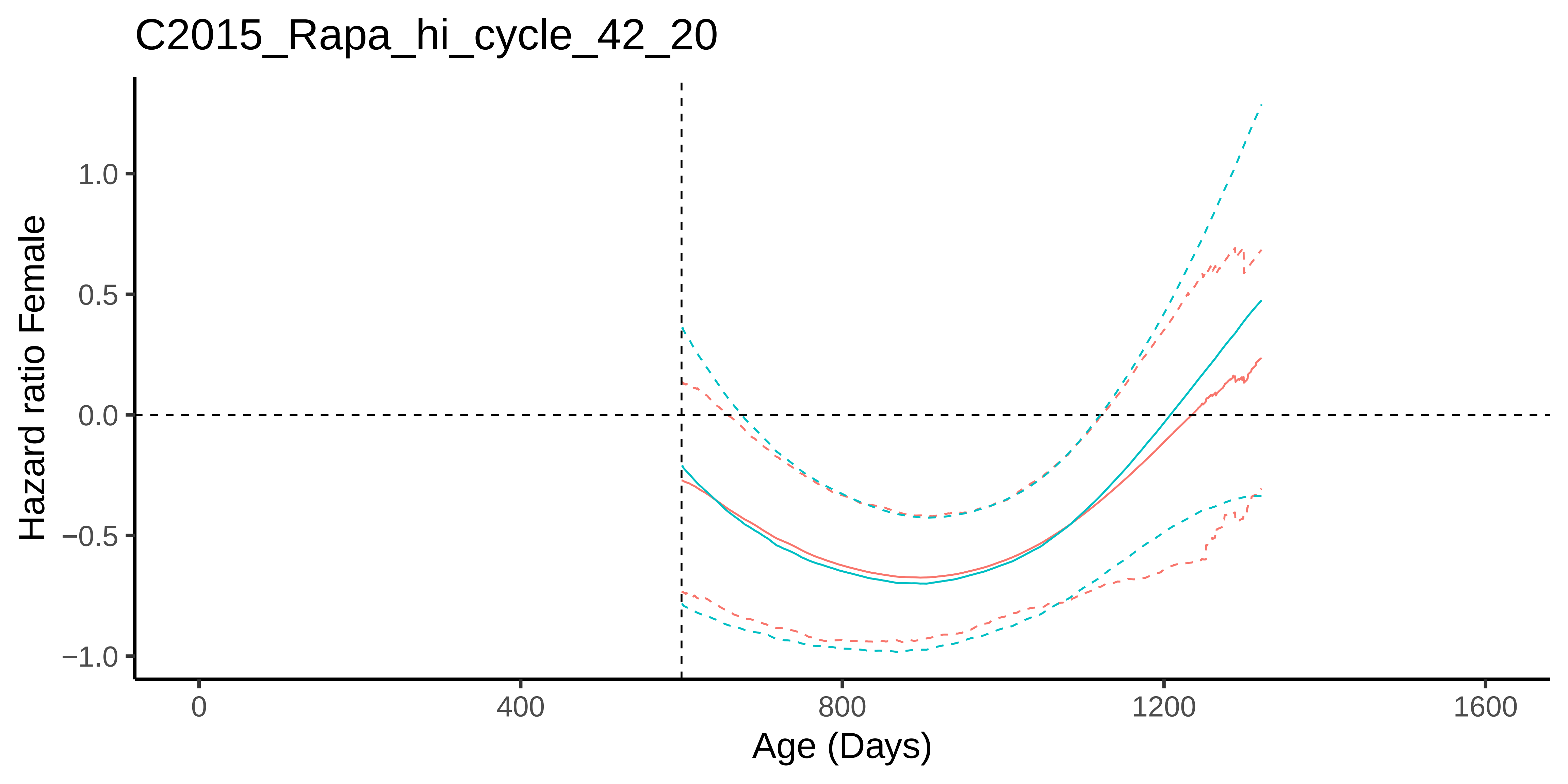
C2015_Rapa_hi_cycle_42_20
Hazard ratio Female
1.0
0.5
0.0
−0.5
−1.0
0
400
800
1200
1600
Age (Days)

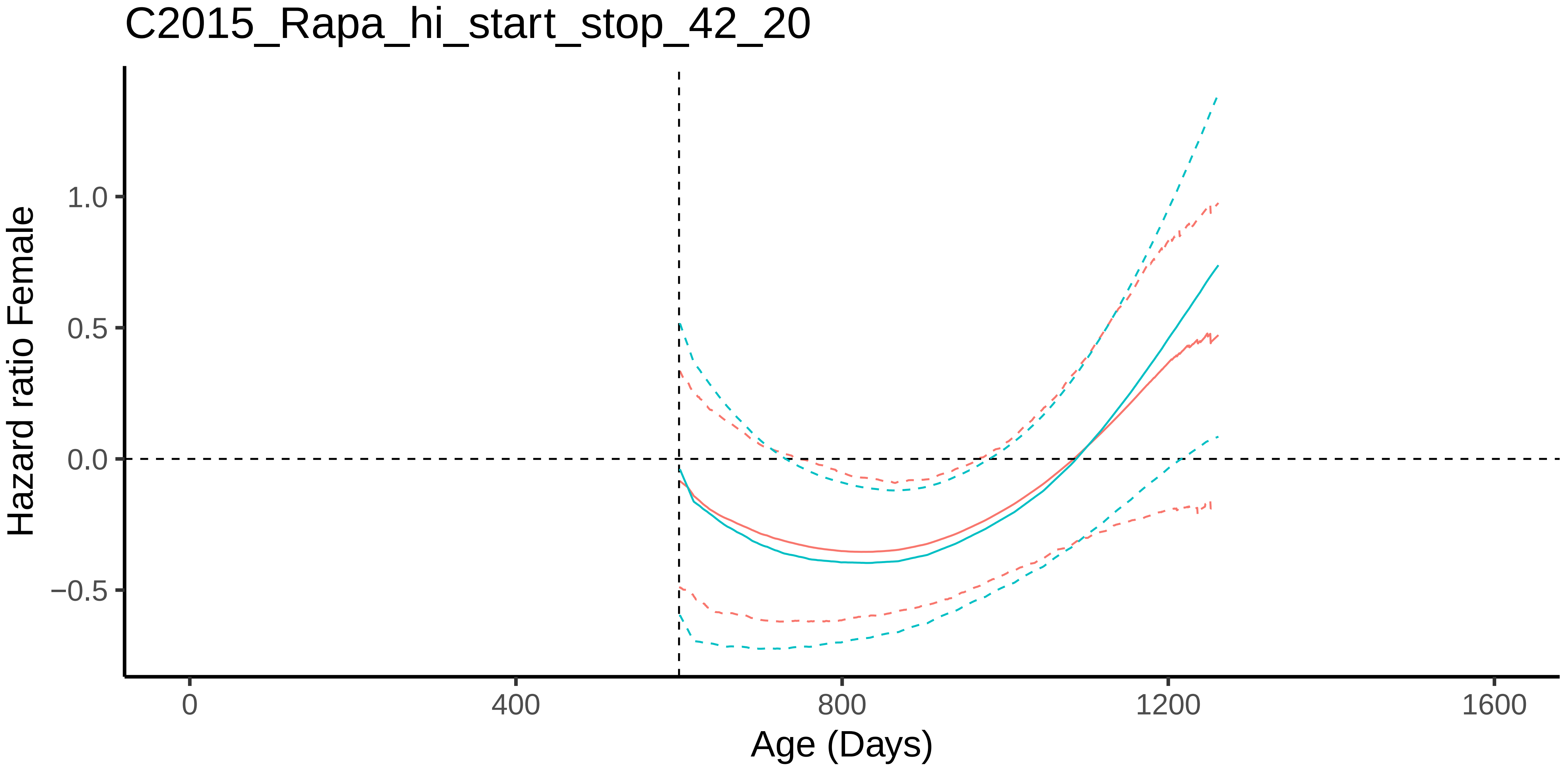

C2015_Rapa_hi_start_stop_42_20
Hazard ratio Female
1.0
0.5
0.0
−0.5
0
400
800
1200
1600
Age (Days)

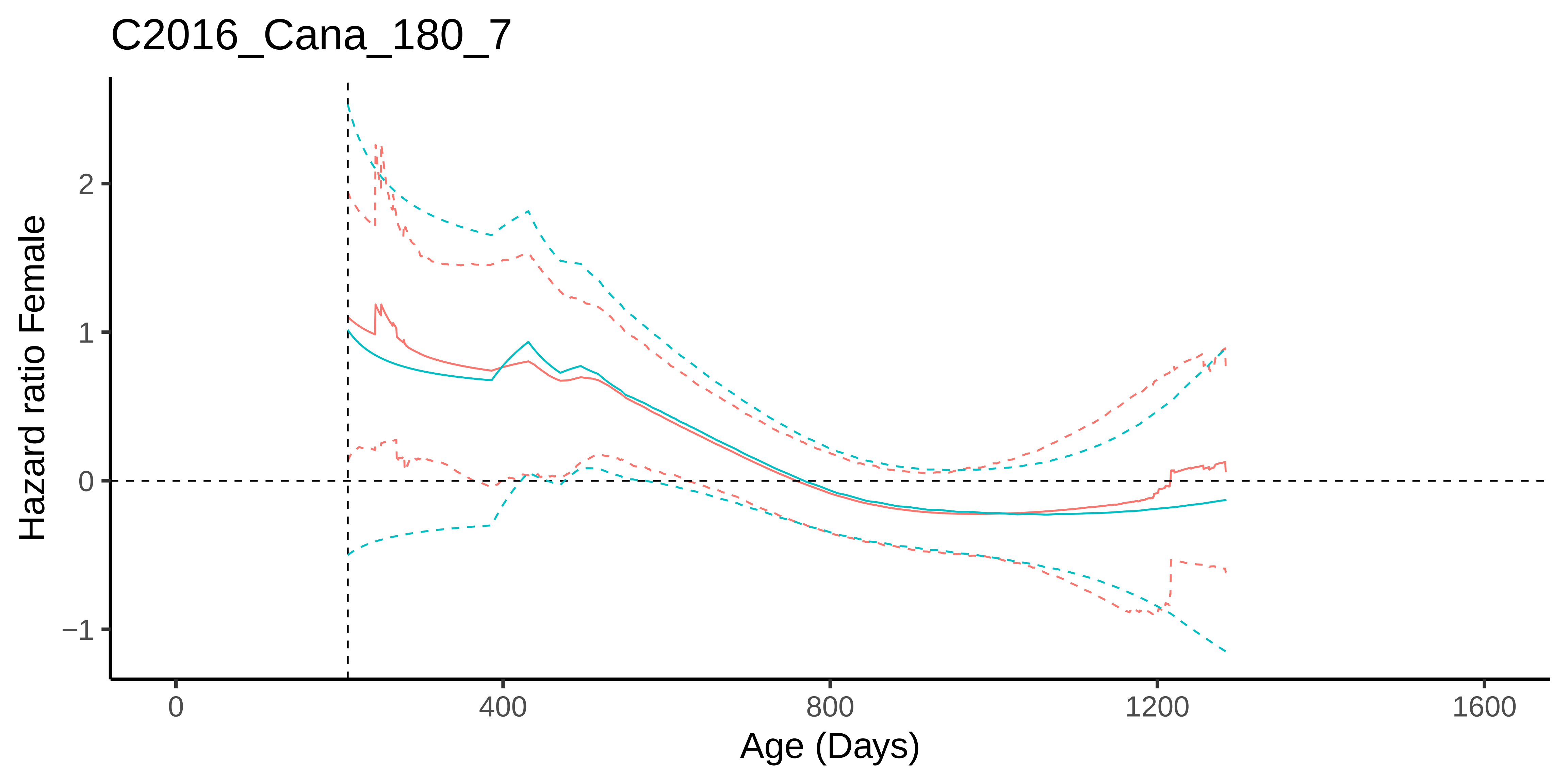
C2016_Cana_180_7
Hazard ratio Female
2
1
0
−1
0
400
800
1200
1600
Age (Days)

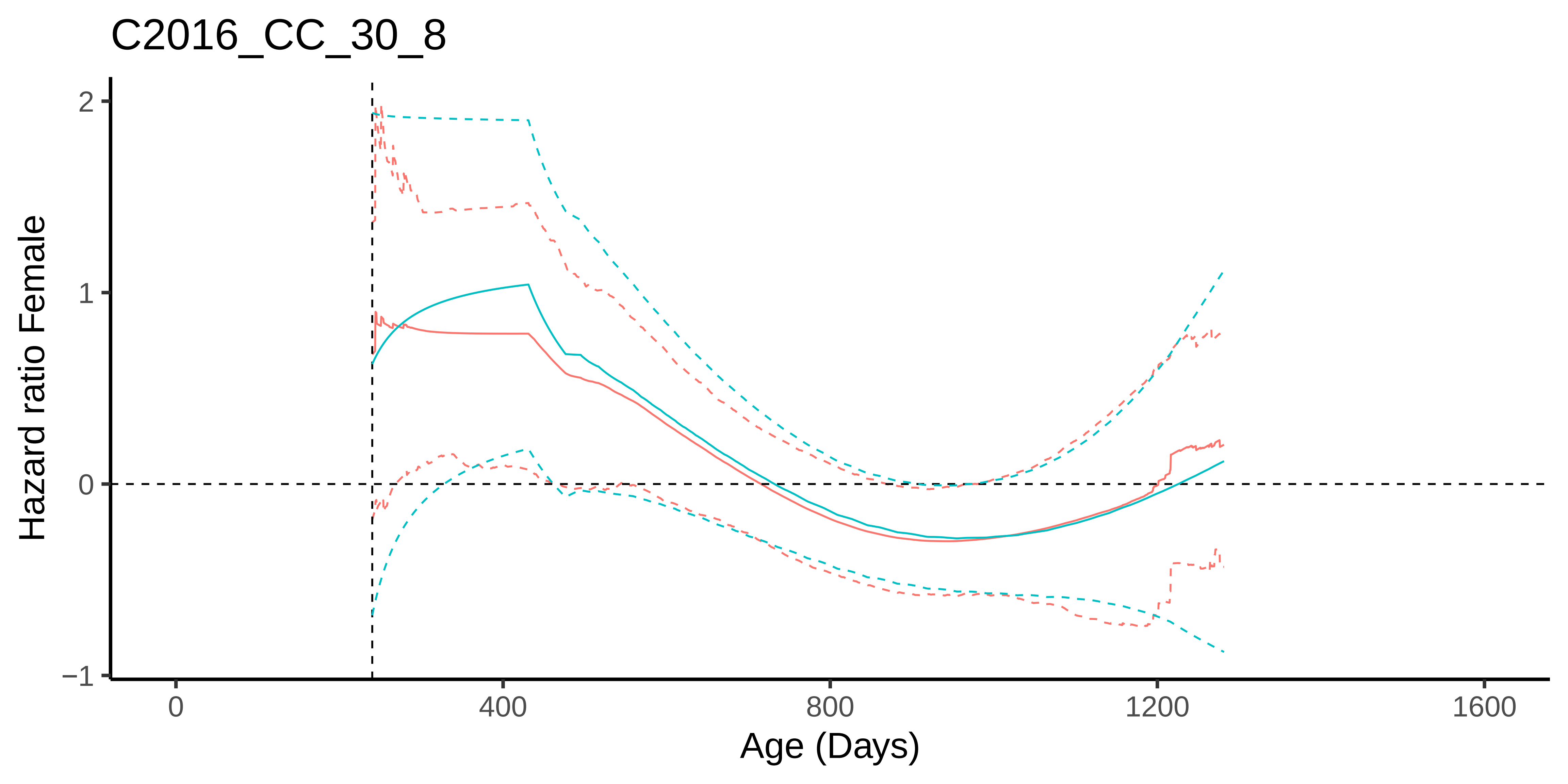

C2016_CC_30_8
Hazard ratio Female
2
1
0
−1
0
400
800
1200
1600
Age (Days)

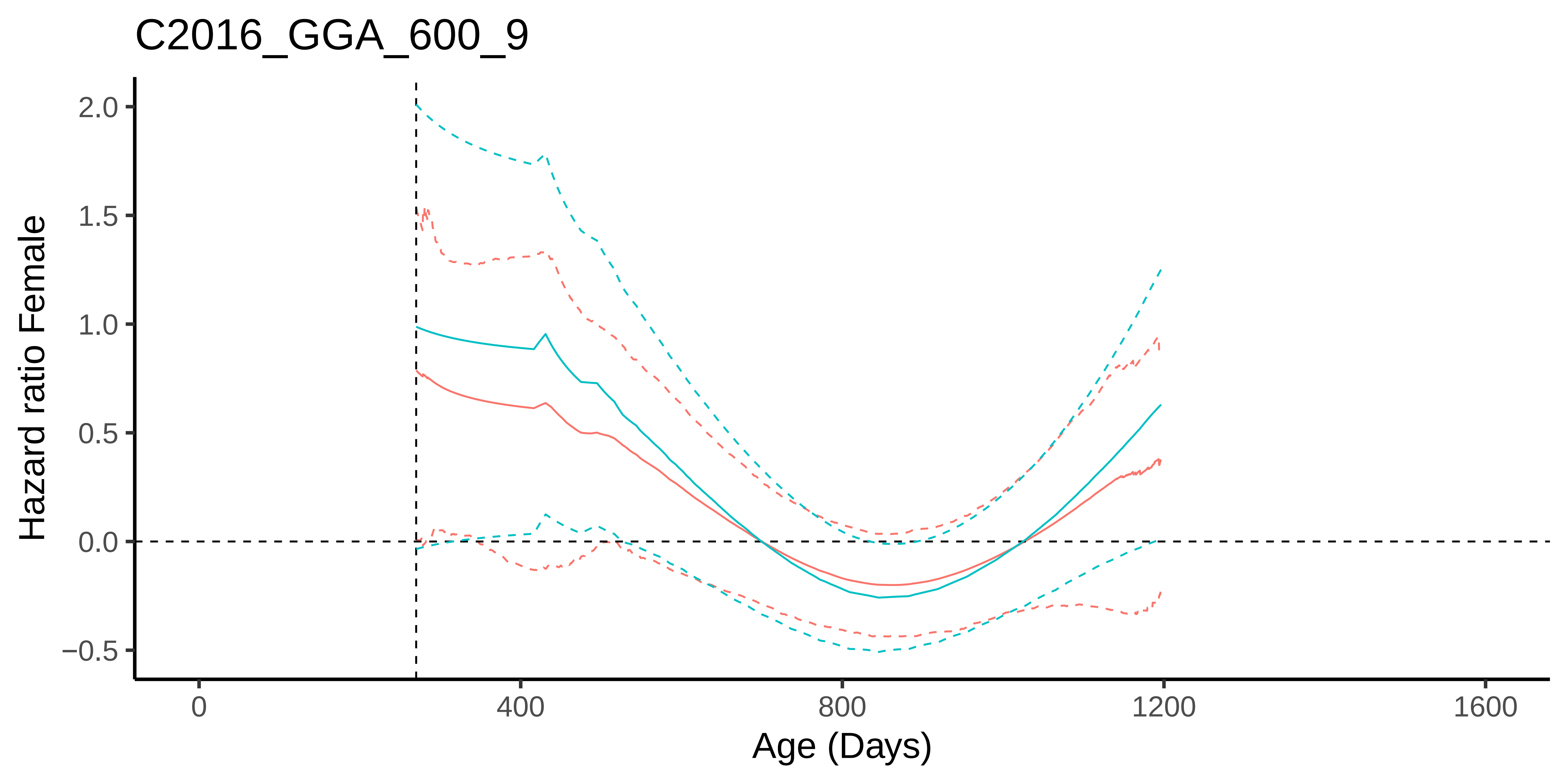
C2016_GGA_600_9
Hazard ratio Female
2.0
1.5
1.0
0.5
0.0
−0.5
0
400
800
1200
1600
Age (Days)

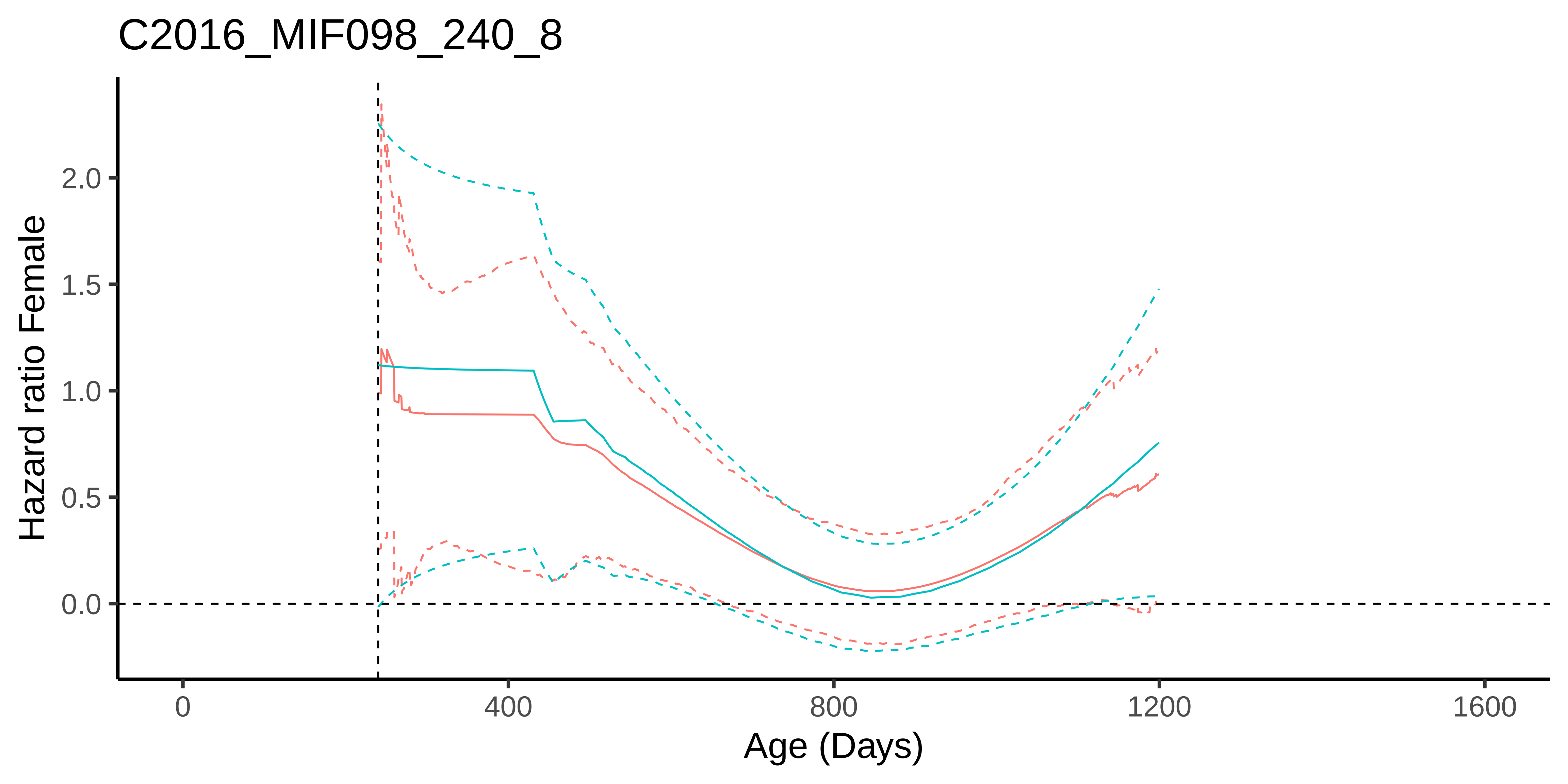
C2016_MIF098_240_8
Hazard ratio Female
2.0
1.5
1.0
0.5
0.0
0
400
800
1200
1600
Age (Days)

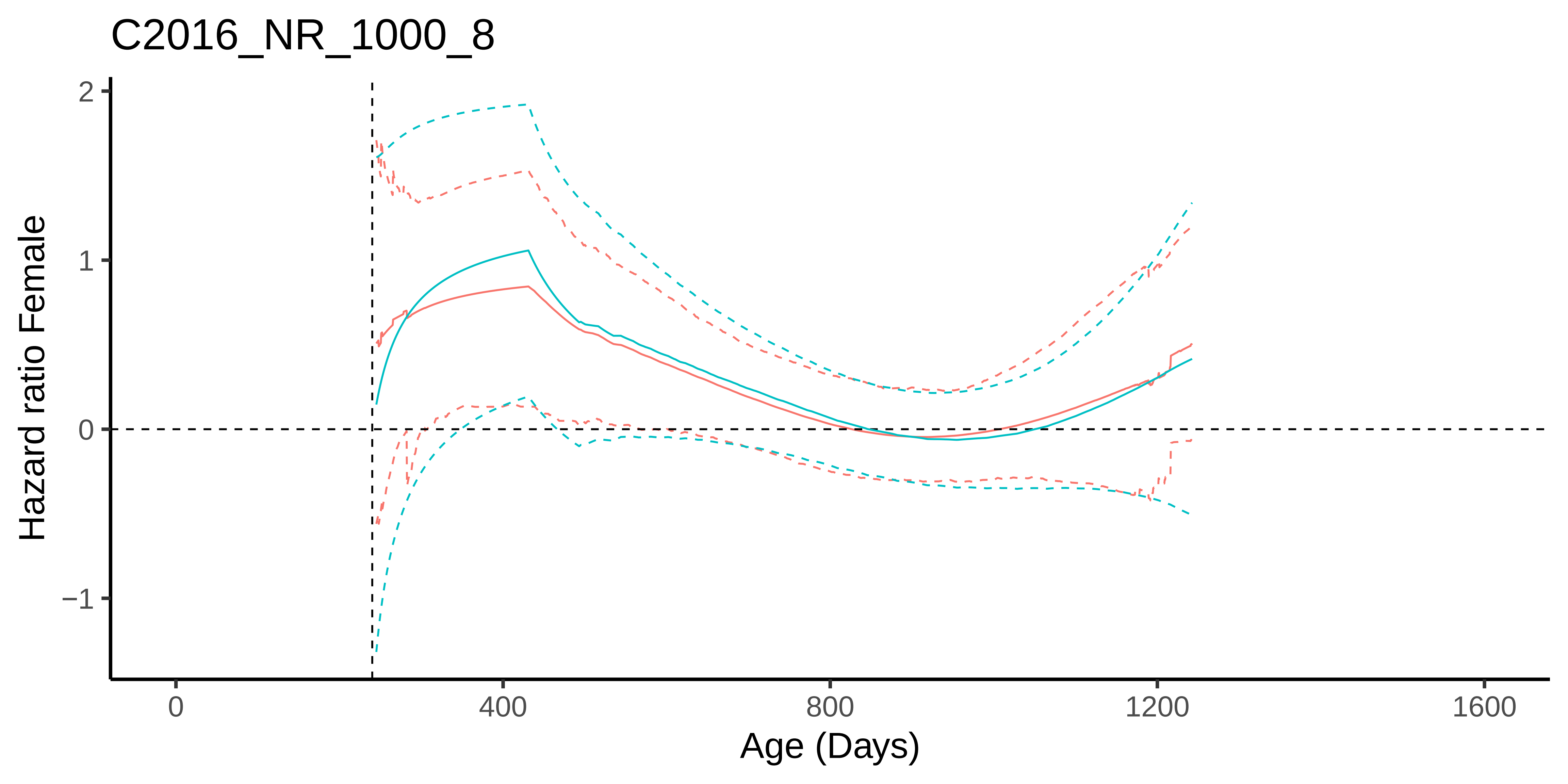
C2016_NR_1000_8
Hazard ratio Female
2
1
0
−1
0
400
800
1200
1600
Age (Days)

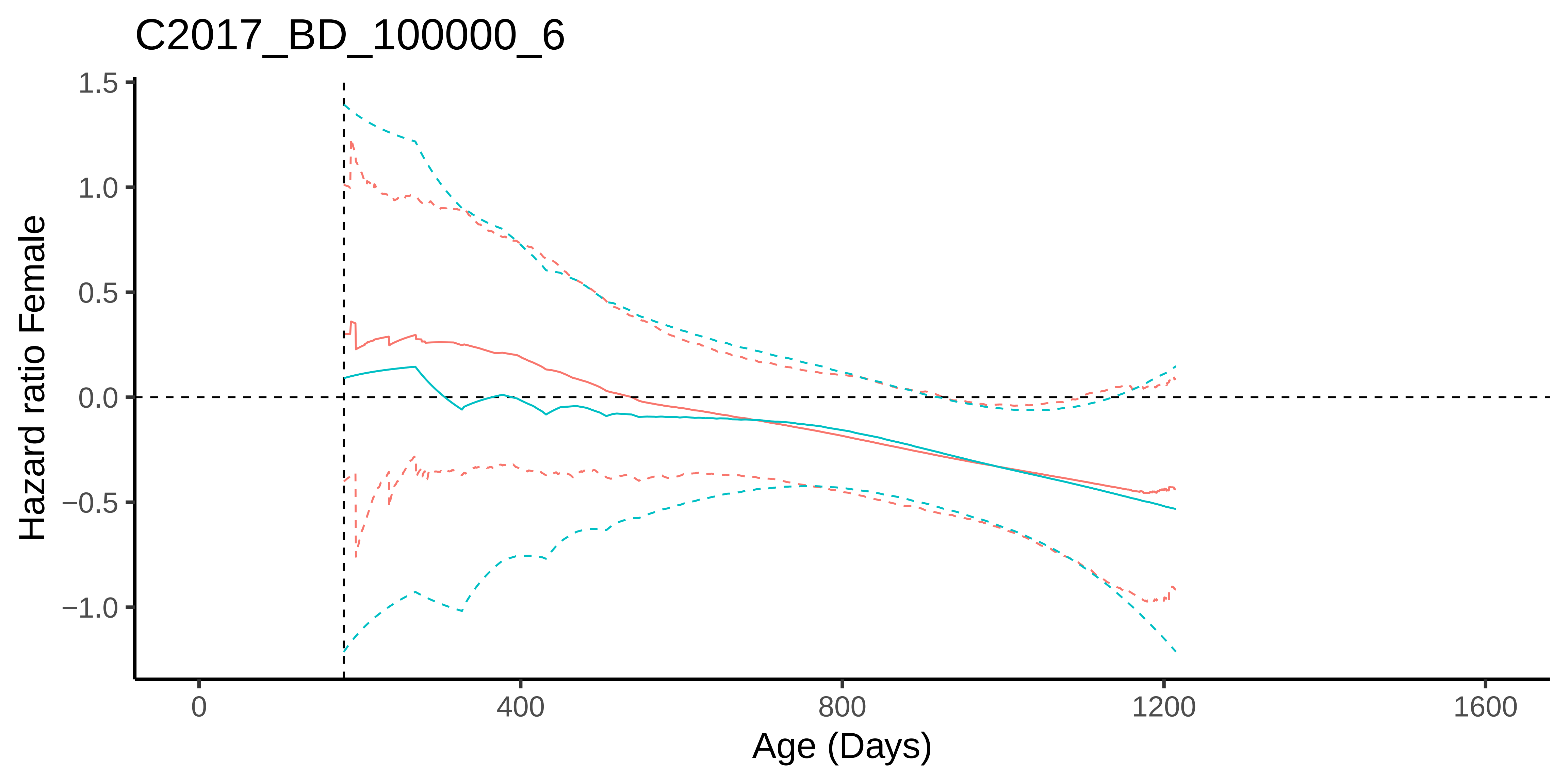
C2017_BD_100000_6
Hazard ratio Female
1.5
1.0
0.5
0.0
−0.5
−1.0
0
400
800
1200
1600
Age (Days)

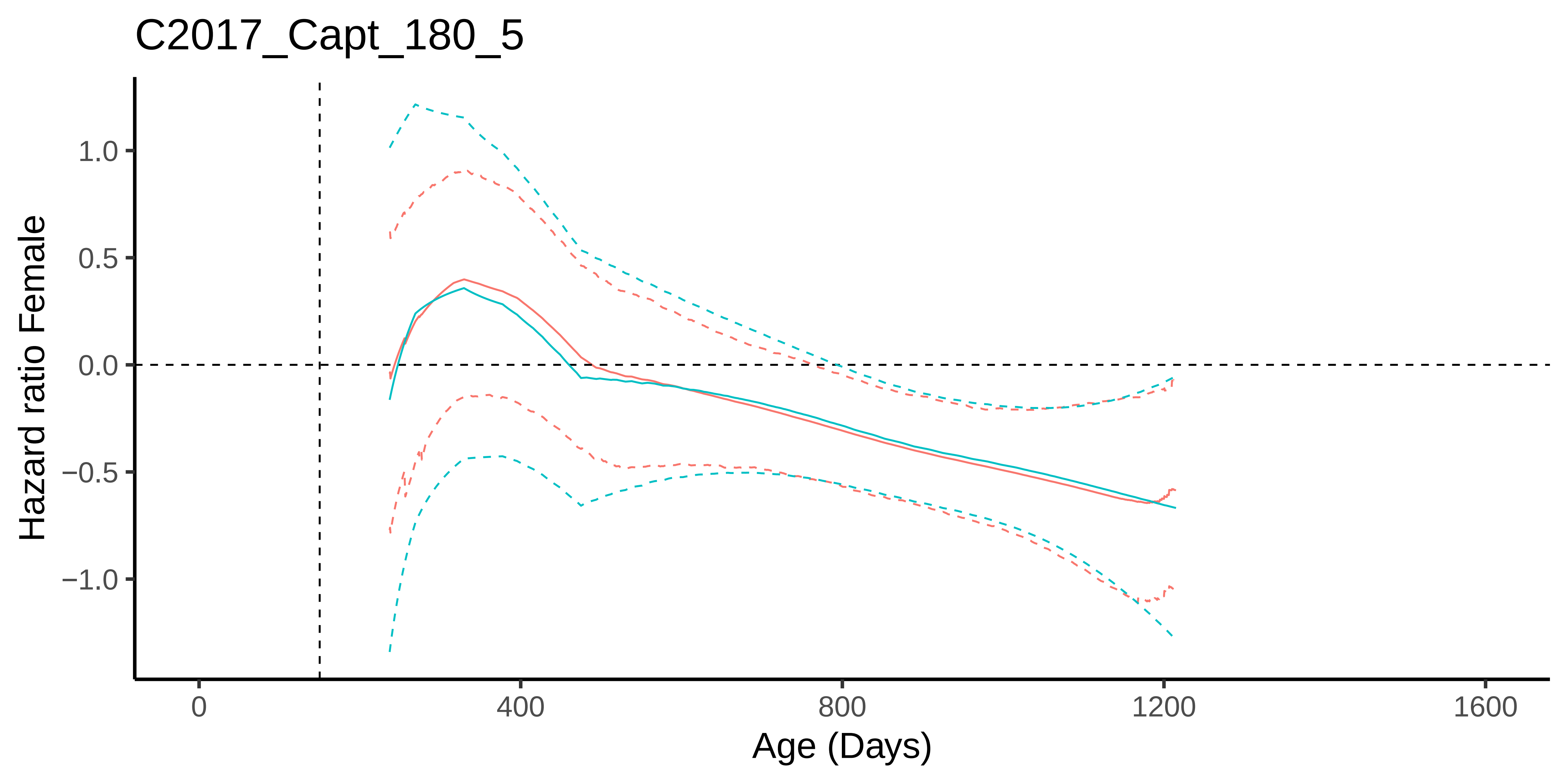

C2017_Capt_180_5
Hazard ratio Female
1.0
0.5
0.0
−0.5
−1.0
0
400
800
1200
1600
Age (Days)

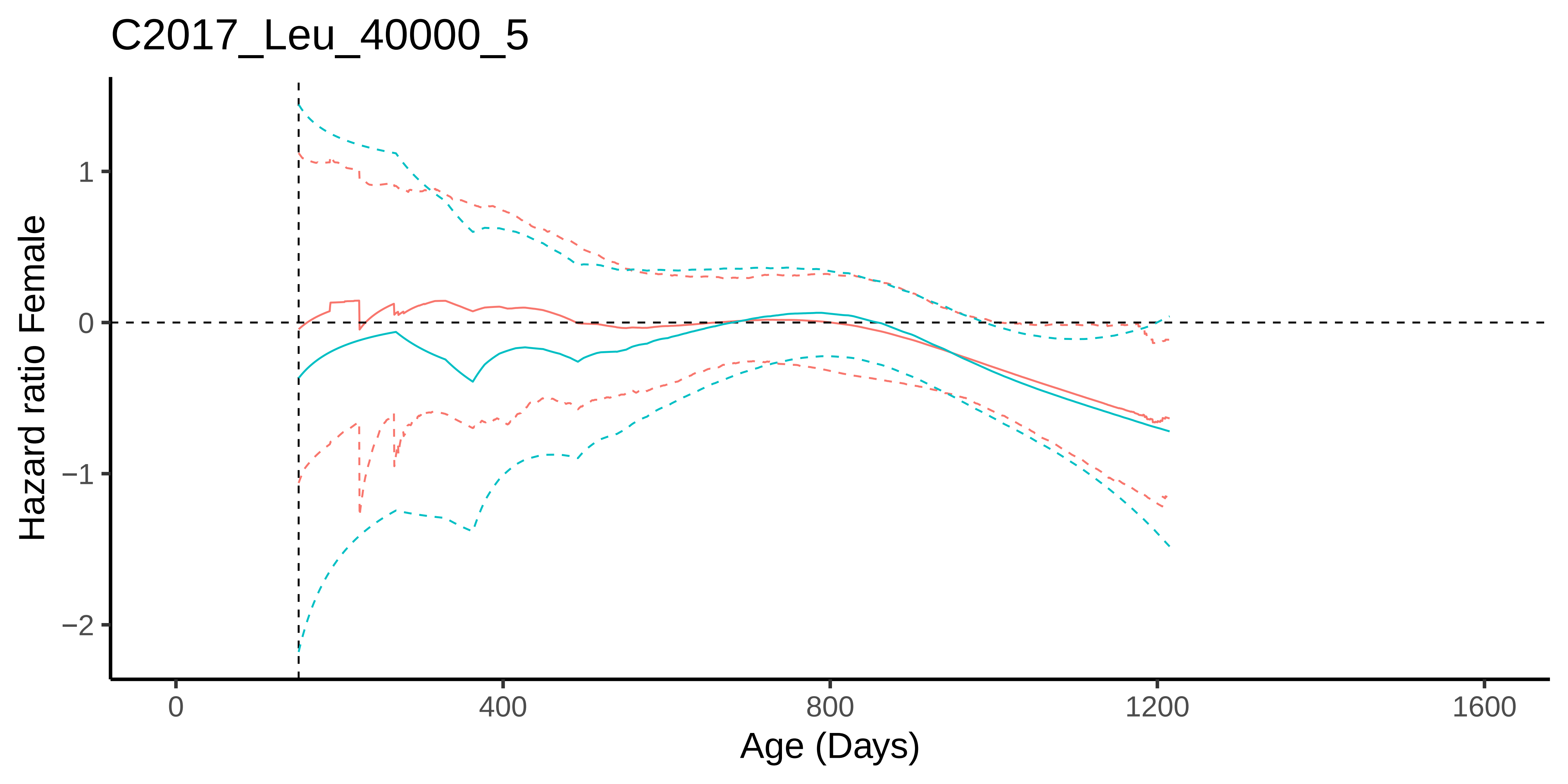

C2017_Leu_40000_5
Hazard ratio Female
1
0
−1
−2
0
400
800
1200
1600
Age (Days)

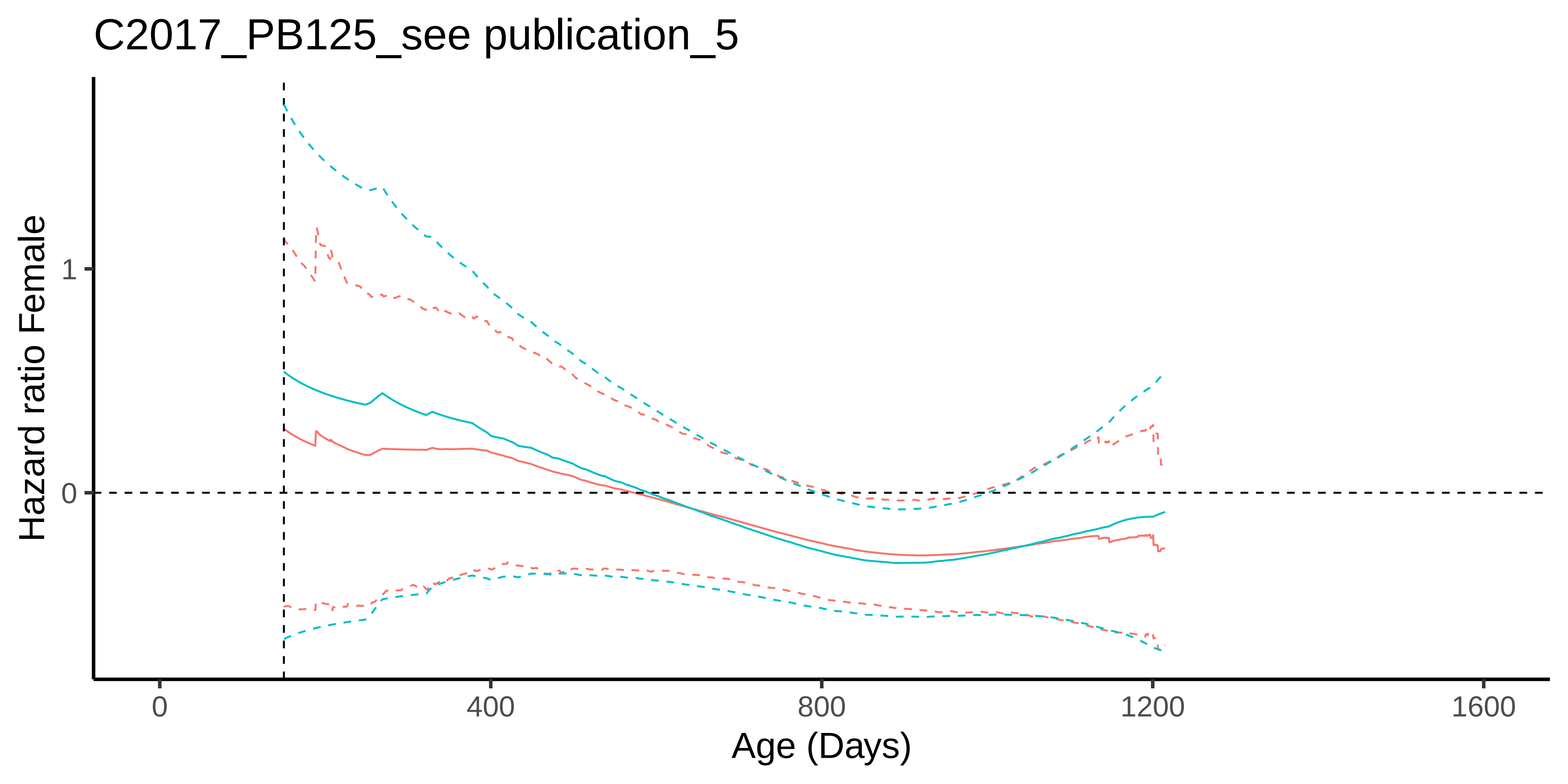
C2017_PB125_see publication_5
Hazard ratio Female
1
0
0
400
800
1200
1600
Age (Days)

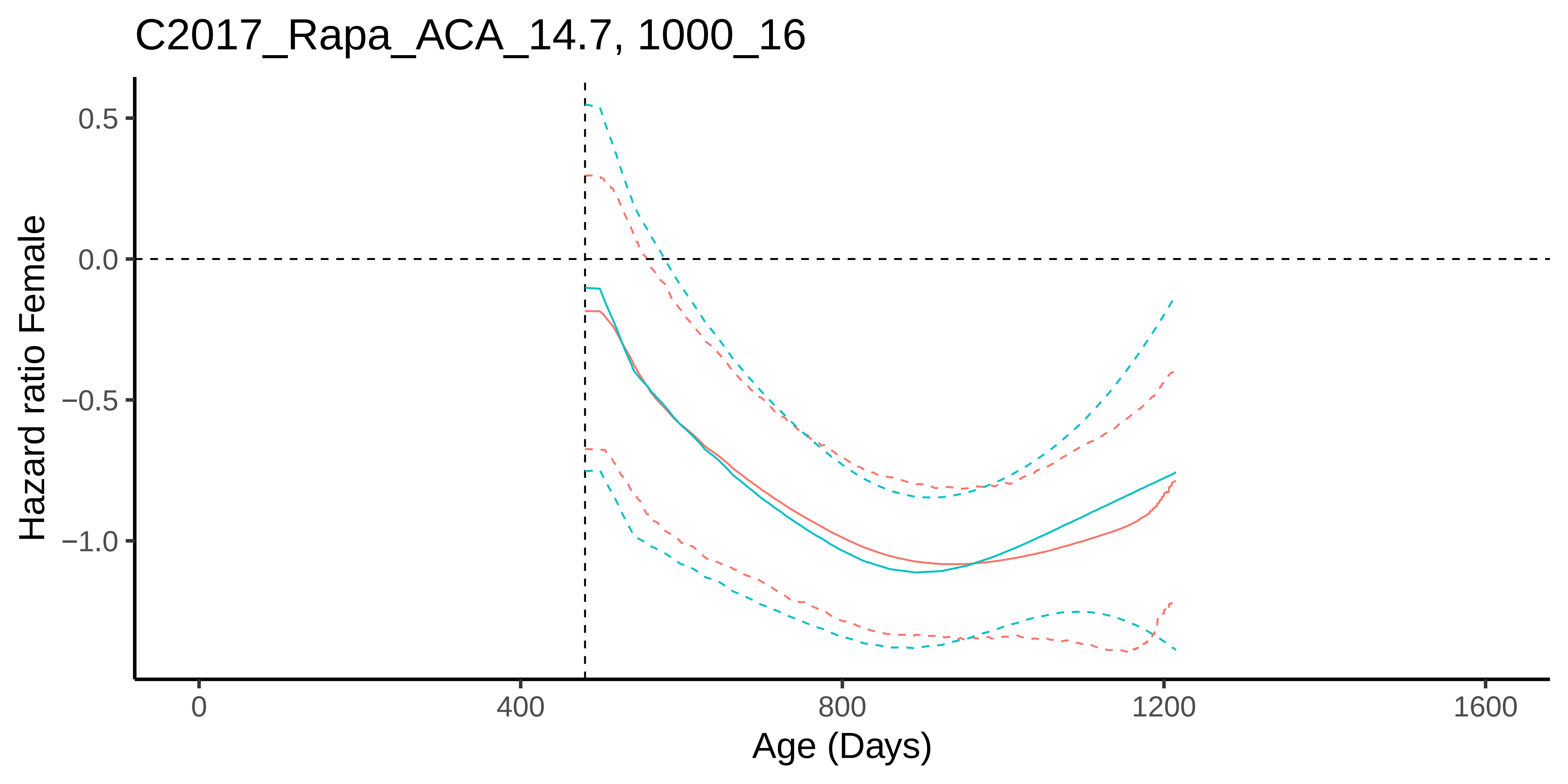
C2017_Rapa_ACA_14.7, 1000_16
0.5
0.0
−0.5
−1.0
Hazard ratio Female
0
400
800
1200
1600
Age (Days)

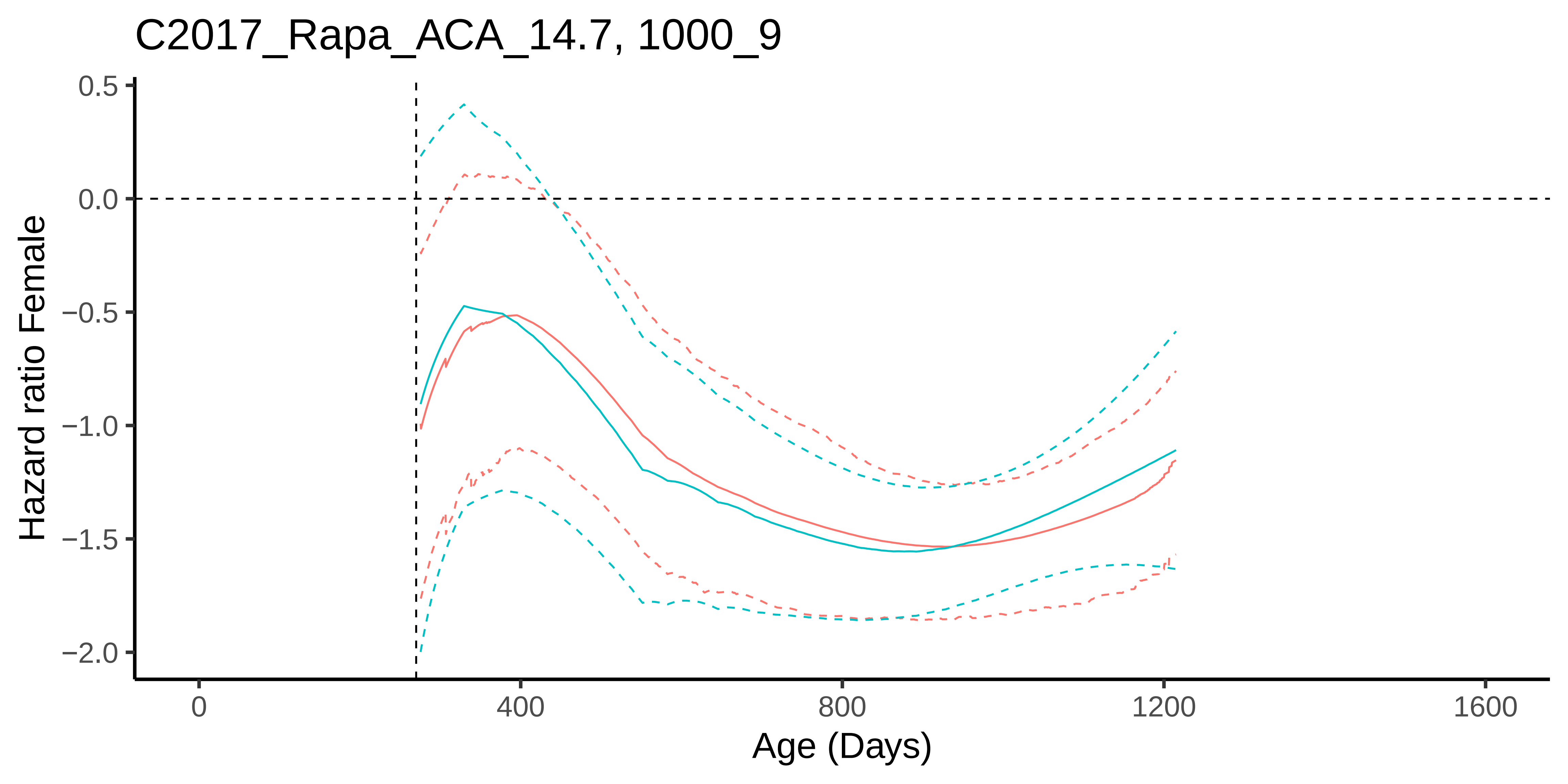
C2017_Rapa_ACA_14.7, 1000_9
Hazard ratio Female
0.5
0.0
−0.5
−1.0
−1.5
−2.0
0
400
800
1200
1600
Age (Days)

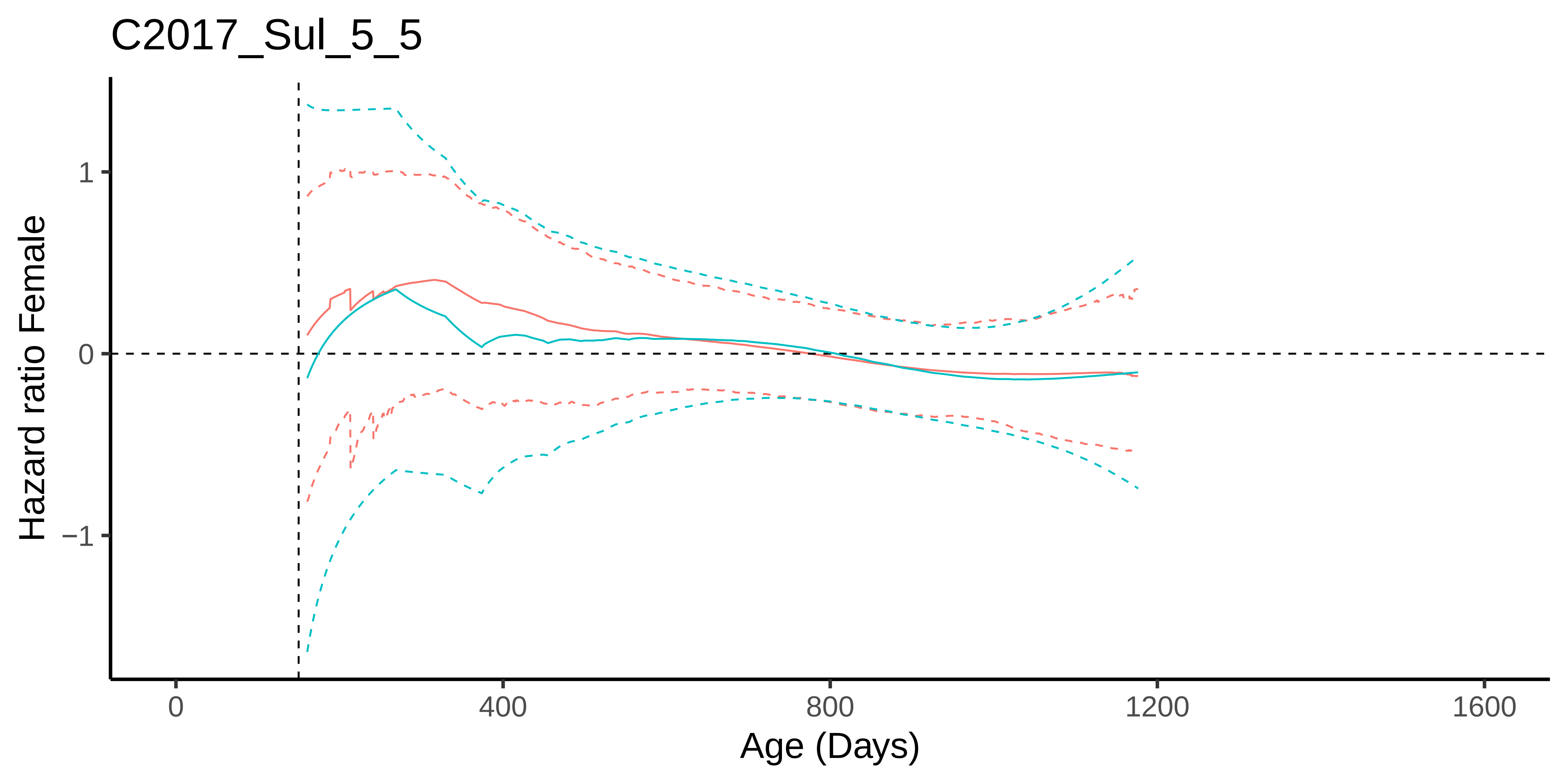
C2017_Sul_5_5
Hazard ratio Female
1
0
−1
0
400
800
1200
1600
Age (Days)

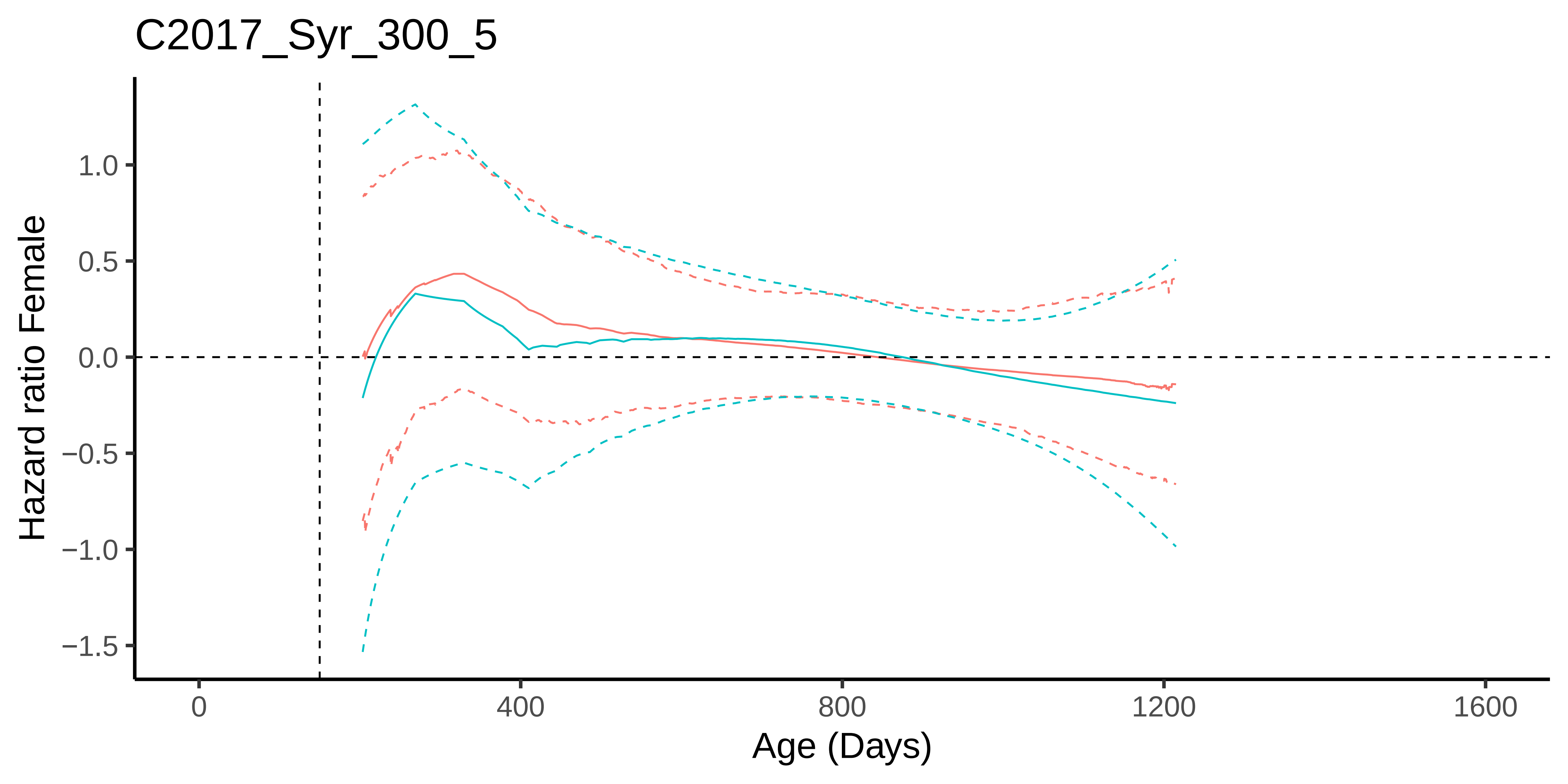
C2017_Syr_300_5
Hazard ratio Female
1.0
0.5
0.0
−0.5
−1.0
−1.5
0
400
800
1200
1600
Age (Days)
